# NUDIX Hydrolases Target Specific Inositol Pyrophosphates and Regulate Phosphate Homeostasis and Bacterial Pathogen Susceptibility in Arabidopsis

**DOI:** 10.1101/2024.10.18.619122

**Authors:** Robin Schneider, Klea Lami, Isabel Prucker, Sara Christina Stolze, Annett Strauß, Julie Marie Schmidt, Simon M. Bartsch, Kevin Langenbach, Esther Lange, Kevin Ritter, David Furkert, Natalie Faiß, Sandeep Kumar, M. Shamim Hasan, Athanasios Makris, Lukas Krusenbaum, Stefanie Wege, Yemisrach Zewdu Belay, Simon Kriescher, Jeremy The, Michael Harings, Florian M. W. Grundler, Martina K. Ried-Lasi, Heiko Schoof, Philipp Gaugler, Marília Kamleitner, Dorothea Fiedler, Hirofumi Nakagami, Ricardo F. H. Giehl, Thomas Lahaye, Saikat Bhattacharjee, Henning J. Jessen, Verena Gaugler, Gabriel Schaaf

**Affiliations:** Department of Plant Nutrition, Institute of Crop Science and Resource Conservation, University of Bonn, 53115 Bonn, Germany; Department of Chemistry and Pharmacy and CIBSS-Centre for Integrative Biological Signalling Studies, Albert-Ludwigs University Freiburg, 79104 Freiburg, Germany; Protein Mass Spectronomy, Max Planck Institute for Plant Breeding Research, 50829 Cologne, Germany; Department of General Genetics, Center of Plant Molecular Biology (ZMBP) University of Tübingen, 72076 Tübingen, Germany; Leibniz-Forschungsinstitut für Molekulare Pharmakologie (FMP), 13125 Berlin, Germany; Humboldt-Universität zu Berlin, Institut für Chemie, 12489 Berlin, Germany; Laboratory of Signal Transduction and Plant Resistance, UNESCO-Regional Centre for Biotechnology (RCB), NCR Biotech Science Cluster, 3rd Milestone, Faridabad-Gurgaon Expressway, Faridabad, 121001, Haryana, India; Molecular Phytomedicine, Institute of Crop Science and Resource Conservation, University of Bonn, 53115 Bonn, Germany; Department of Molecular Signal Processing, Leibniz Institute of Plant Biochemistry, 06120 Halle (Saale), Germany; Department of Crop Bioinformatics, Institute of Crop Science and Resource Conservation, University of Bonn, 53115 Bonn, Germany; Department of Physiology & Cell Biology, Leibniz-Institute of Plant Genetics and Crop Plant Research (IPK), 06466 Gatersleben, Germany

## Abstract

Inositol pyrophosphates (PP-InsPs) are important signaling molecules that regulate diverse cellular processes in eukaryotes, including energy homeostasis, phosphate (P_i_) signaling, and phytohormone perception. Yet, in plants, the enzymes responsible for their turnover remain largely unknown. Using a non-hydrolysable PP-InsP analog in a pull-down approach, we identified a family of Arabidopsis NUDIX hydrolases (NUDTs) that group into two closely related subclades. Through *in vitro* assays, heterologous expression systems, and higher-order gene-edited mutants, we explored the substrate specificities and physiological roles of these hydrolases. Using a combination of strong anion exchange (SAX)-HPLC, PAGE, and capillary electrophoresis electrospray ionization mass spectrometry (CE-ESI-MS), we found that their PP-InsP pyrophosphatase activity is enantiomer-selective and Mg^2+^-dependent. Specifically, subclade I NUDTs preferentially hydrolyze 4-InsP_7_, while subclade II NUDTs target 3-InsP_7_, with minor activity against other PP-InsPs, including 5-InsP_7_. In higher-order mutants of subclade II NUDTs, we observed defects in both P_i_ and iron homeostasis, accompanied by increased levels of 1/3-InsP_7_ and 5-InsP_7_, with a markedly larger increase in 1/3-InsP_7_. Ectopic expression of NUDTs from both subclades induced local P_i_ starvation responses (PSRs), while RNA-seq analysis comparing wildtype (WT) and subclade II *nudt12/13/16* loss-of-function plants indicates additional PSR-independent roles, potentially involving 1/3-InsP_7_ in the regulation of plant defense. Consistently, *nudt12/13/16* mutants displayed enhanced resistance to *Pseudomonas syringae* infection, indicating a role in bacterial pathogen susceptibility. Expanding beyond subclade II NUDTs, we demonstrated susceptibility of the 3PP-position of PP-InsPs to enzymatic activities unrelated to NUDTs, and found that such activities are conserved across plants and humans. Additionally, we found that NUDT effectors from pathogenic ascomycete fungi exhibit a substrate specificity similar to subclade I NUDTs. Collectively, our findings reveal new roles for NUDTs in PP-InsP signaling, plant nutrient and immune responses, and highlight a cross-kingdom conservation of PP-InsP-metabolizing enzymes.

## Introduction

Inositol pyrophosphates (PP-InsPs), such as InsP_7_ and InsP_8_, are small signaling molecules consisting of a phosphorylated *myo*-inositol ring and one or more pyrophosphate groups, which regulate a wide range of cellular processes in eukaryotes. In metazoans, these messengers control DNA repair, insulin signaling, blood clotting, ribosome biogenesis, spermiogenesis, and telomere length maintenance (Stephens et al. 1993; Menniti et al. 1993; Shears 2015, 2018; Thota and Bhandari 2015). In plants, PP-InsPs regulate jasmonate and auxin signaling, as well as a nutrient status-dependent accumulation of storage lipids (Laha et al. 2015, 2016, 2022; Couso et al. 2016). An emerging and somewhat unifying role of these messengers in fungi, metazoans and plants is their critical involvement in energy homeostasis and phosphate (P_i_) signaling (Lee et al. 2007; Szijgyarto et al. 2011; Wilson et al. 2013; Wild et al. 2016; Dong et al. 2019; Zhu et al. 2019; Li et al. 2020; Ried et al. 2021; Wang et al. 2021; Riemer et al. 2021; Gulabani et al. 2022; Qin et al. 2023; Chabert et al. 2023; Raj et al. 2024). Phosphorus (P) is an essential macronutrient and a key determinant of agricultural yield (Tilman et al. 2002; MacDonald et al. 2011; Hawkesford et al. 2023). Plants have evolved complex sensing and signaling mechanisms to adjust the plant’s P demand with external P_i_ availability. In Arabidopsis, the MYB transcription factors Phosphate Starvation Response 1 (PHR1) and PHR1-like 1 (PHL1) regulate the expression of P_i_ starvation-induced (PSI) genes (Rubio et al. 2001; Bustos et al. 2010). Under P_i_ deficiency, PHR1 activates PSI genes by binding to cognate binding elements (P1BS) in their promoter region. Under P_i_ sufficiency, stand-alone SYG1/Pho81/XPR1 (SPX) proteins negatively regulate PHR transcription factors, as shown in various plant species (Liu et al. 2010; Shi et al. 2014; Lv et al. 2014; Puga et al. 2014; Wang et al. 2014; Qi et al. 2017; Zhong et al. 2018; Osorio et al. 2019; Ried et al. 2021). Intriguingly, SPX proteins function as cellular receptors for certain PP-InsPs, such as 5-InsP_7_ and InsP_8_ (Wild et al. 2016; Gerasimaite et al. 2017; Dong et al. 2019; Zhu et al. 2019; Li et al. 2020; Ried et al. 2021; Pipercevic et al. 2023; Yan et al. 2024). When bound to InsP_8_, SPX proteins interact with the unique coiled-coil motif of PHR1, interfering with its oligomerization and promoter-binding activity (Ried et al. 2021). Consequently, mutants defective in these PP-InsPs display constitutively up-regulated P_i_ starvation responses (PSRs) both locally and systemically, suggesting that plants are largely unable to sense P_i_ directly but sense certain PP-InsPs as a proxy for P_i_ (Dong et al. 2019; Zhu et al. 2019; Riemer et al. 2021). However, the regulation of these PSRs and the precise mechanisms and regulation of P_i_ sensing in plants still remains poorly understood.

Inositol polyphosphates (InsPs) and PP-InsPs have a high negative charge density, and many of their isomers are present at low cellular abundance, making them particularly challenging to analyze. Moreover, most methods fail to accurately separate PP-InsP isomers with identical molecular masses. A major breakthrough came with the development of capillary electrophoresis electrospray ionization mass spectrometry (CE-ESI-MS) (Qiu et al. 2020), which enabled the identification of three major InsP_7_ (also referred to as PP-InsP_5_) isomers in plant extracts: 1-InsP_7_ (carrying the PP-moiety at the 1-position) and/or 3-InsP_7_ (hereafter referred to as 1/3-InsP_7_), 4-InsP_7_ and/or 6-InsP_7_ (hereafter referred to as 4/6-InsP_7_), and 5-InsP_7_ (Riemer et al. 2021). Because InsP_6_, like *myo*-inositol, is a *meso* compound with a plane of symmetry dissecting the 2 and 5 positions, the 1/3 and 4/6 phosphates are enantiomeric and cannot be distinguished in the absence of chiral selectors (Blüher et al. 2017; Ritter et al. 2023) (Figure S1). In addition, CE-ESI-MS allowed identification of a novel PP-InsP_4_ isomer of unknown isomer identity in Arabidopsis roots (Riemer et al. 2021; Ritter et al. 2025).

The biosynthetic pathways of PP-InsPs are only partially resolved. Even though 5-InsP_7_ appears to be present in all eukaryotes examined to date, its synthesis from phytate (InsP_6_) in different organisms is mediated by distinct, sequence-unrelated enzyme families. In plants, 5-InsP_7_ synthesis is catalyzed by inositol (1,3,4) trisphosphate 5/6-kinases (ITPKs). This activity has been detected *in vitro* via enzymatic assays with recombinant protein, through heterologous expression of various Arabidopsis and rice ITPKs in yeast (Laha et al. 2019; Whitfield et al. 2020), and *in planta* for Arabidopsis ITPK1 and ITPK2 (Riemer et al. 2021; Laha et al. 2022).

The synthesis of InsP_8_ from 5-InsP_7_ appears to be largely conserved in eukaryotes and is catalyzed by members of a protein family distinct from the aforementioned InsP_6_ kinases (Mulugu et al. 2007). In Arabidopsis, this reaction is catalyzed by two PPIP5K isoforms, namely VIH1 and VIH2 (Desai et al. 2014; Laha et al. 2015; Dong et al. 2019; Zhu et al. 2019). Notably, only a few PPIP5K enzymes, such as budding yeast Vip1 and human PPIP5K2, have been shown to specifically generate 1-InsP_7_ and 1,5-InsP_8_ (Mulugu et al. 2007; Fridy et al. 2007; Wang et al. 2011; Capolicchio et al. 2014; Dollins et al. 2020). For most other plant and mammalian PPIP5K isoforms, it remains unclear whether they catalyze the formation of the phosphoanhydride bond at the 1- or 3-position of the fully phosphorylated *myo*-inositol ring. This ambiguity is further complicated by the potential involvement of inositol phosphate isomerases, such as ITPKs, in generating these 1/3-InsP_7_ and 1/3,5-InsP_8_ species (Sweetman et al. 2007; Josefsen et al. 2007; Caddick et al. 2008). Consequently, the enantiomeric identity of 1/3-InsP_7_ and 1/3,5-InsP_8_ remains unresolved in eukaryotes.

Our understanding of PP-InsP turnover in plants also remains incomplete. Similar to yeast and mammalian PPIP5Ks, AtVIH1 and AtVIH2 are bifunctional enzymes, harboring an N-terminal ATP-grasp kinase domain and a C-terminal phosphatase-like domain (Mulugu et al. 2007; Fridy et al. 2007; Wang et al. 2011; Laha et al. 2015; Zhu et al. 2019). The phosphatase domain of Arabidopsis VIH2 hydrolyzes 1-InsP_7_, and 5-InsP_7_ to InsP_6_ *in vitro* while the fission yeast PPIP5K homolog Asp1 and budding yeast Vip1 can convert 1-InsP_7_ and 1,5-InsP_8_ to InsP_6_ and 5-InsP_7_, respectively (Wang et al. 2015; Pascual-Ortiz et al. 2018; Zhu et al. 2019; Dollins et al. 2020). The synthesis and turnover of specific PP-InsPs are tightly linked to the cellular adenylate energy charge (i.e., the ATP/ADP ratio). For example, ATP modulates the balance between the kinase and phosphatase activities of Arabidopsis VIH proteins *in vitro* (Zhu et al. 2019), indicating that PP-InsP levels are sensitive to energy availability. Similarly, the ITPK1-mediated synthesis of 5-InsP_7_ depends on high ATP concentrations, reflecting its relatively high K_m_ for ATP. Under low-energy conditions, ITPK1 can catalyze a reverse reaction, transferring the β-phosphate from 5-InsP_7_ to ADP, thereby regenerating ATP and producing InsP_6_. This reverse phosphotransferase activity likely represents a rapid and energy-efficient mechanism to suppress PSRs when cellular energy is limited. Findings from Whitfield et al. (2020) and Riemer et al. (2021) highlight a conserved principle across eukaryotes, including mammals, where structurally distinct InsP_6_ kinases exhibit similar energy-dependent regulatory behavior. Unlike PPIP5Ks, the dual catalysis of ITPK1 does not rely on distinct functional domains, but is executed by a single atypical ATP-grasp fold kinase domain (Zong et al. 2022).

Recently, a family of specific inositol phosphohydrolases, termed *Arabidopsis thaliana* Plant and Fungi Atypical Dual Specificity Phosphatases (PFA-DSP1–5), has been identified. Similarly to the activity of the budding yeast homologue Siw14 (Steidle et al. 2016; Wang et al. 2018), all five PFA-DSP isoforms displayed robust and highly specific pyrophosphatase activities towards the 5-β-phosphates of 5-InsP_7_ and 1/3,5-InsP_8_ (Wang et al. 2022; Gaugler et al. 2022; Laurent et al. 2024). However, the enzymes responsible for hydrolyzing other InsP_7_ isomers, such as 1-InsP_7_ and/or 3-InsP_7_, and 4-InsP_7_ and/or 6-InsP_7_, remain largely unresolved in plants.

To this end, we took advantage of a non-hydrolysable InsP_7_ analog, 5PCP-InsP_5_ (Wu et al. 2016), in which the oxygen of a phosphoanhydride bond is substituted by a carbon, to carry out affinity pull-down experiments and identified Arabidopsis NUDIX proteins as potential interactors of InsP_7_. NUDIX is an acronym for ‘NUcleoside DIphosphate linked to some other moiety X,’ reflecting the diverse nucleoside diphosphate substrates targeted by NUDIX-type (NUDT) hydrolases. The budding yeast Ddp1 (Safrany et al. 1999; Lonetti et al. 2011; Andreeva et al. 2019) and the five human DIPP proteins (DIPP1, 2α, 2β, 3α and 3β) (Kilari et al. 2013) are members of the NUDIX family and were shown to hydrolyze preferentially the β-phosphate of 1-InsP_7_. Recent independent publications also explore broader roles of NUDIX hydrolases in PP-InsP metabolism (Chalak et al. 2024; Laurent et al. 2024; Freed et al. 2025). However, these studies did not investigate the isomer-specific turnover of 3-InsP_7_ or 4/6-InsP_7_, nor the involvement of NUDTs in plant defense, which we characterize here.

We provide evidence that members of two different subclades of these phosphohydrolases display distinct PP-InsP pyrophosphatase activities that are highly specific and enantiomer-selective: subclade I members NUDT4, NUDT17, NUDT18 and NUDT21 show a preference for the 4-PP moiety, and subclade II members NUDT12, NUDT13 and NUDT16 show a preference for the 3-PP moiety of PP-InsPs. We further demonstrate that subclade II members play roles in P_i_ and iron (Fe) homeostasis, regulate defense gene expression, and contribute to basal resistance against bacterial pathogens. In addition, we show that the 3PP-position of PP-InsPs is a preferred substrate for conserved enzymatic activities unrelated to NUDTs.

## Results

### *Arabidopsis thaliana* NUDT hydrolases physically interact with a non-hydrolysable InsP_7_ analog

To identify potential PP-InsP interacting proteins, we performed an affinity pull-down assay using 5PCP-InsP_5_ beads, a reagent consisting of a non-hydrolysable InsP_7_ analog coupled to a solid phase resin (Wu et al. 2016; Furkert et al. 2021), which were incubated with Arabidopsis shoot or root extracts. Bound proteins were eluted with a high molar excess of InsP_6_, and both beads and eluates were digested with trypsin. The resulting peptides were analyzed separately for eluate and on-bead fractions by liquid chromatography–mass spectrometry (LC-MS/MS) to identify proteins interacting with InsP_7_.

The peptide analysis recovered several known proteins involved in InsP/PP-InsP metabolism such as MIPS3 (Luo et al. 2011), IPK2ɑ and IPK2β (Stevenson-Paulik et al. 2002, 2005; Xia et al. 2003), ITPK3 (Sweetman et al. 2007), VIH1, and VIH2. Additional known InsP_6_/PP-InsP-binding proteins were also enriched, including the jasmonate co-receptor F-box protein COI1 (Sheard et al. 2010), various TIR1/COI1 homologs (AFB1, AFB2, AFB3), components of the TIR1/COI1-dependent SKP1-CUL1-F-box protein (SCF) ubiquitin E3 ligase complex (ASK1, ASK2, CUL1, CUL2) (Laha et al. 2022), several CSN subunits (Walia et al. 2025), and the presumed InsP_8_ binding proteins SPX1 (Dong et al. 2019) and SPX2 (Ried et al. 2021) (Table S1).

In addition to these known interactors, peptides of five different NUDIX (NUDT) hydrolases were identified in shoot and root extracts. NUDT4, NUDT17 and NUDT21, were detected in shoot samples, while NUDT16 and NUDT18 were recovered from root extracts (Table S1). Phylogenetic analysis of the 29 Arabidopsis NUDT hydrolases revealed that these five candidates cluster together, along with two additional members, NUDT12 and NUDT13, in a distinct clade (Figure 1A). This clade can be subdivided into two subclades: subclade I, comprising NUDT4, 17, 18, and 21, and subclade II, comprising NUDT12, 13, and 16. Pairwise protein sequence comparisons illustrate the high sequence similarity within and between these subclades (Figure 1B).

**Figure 1:**
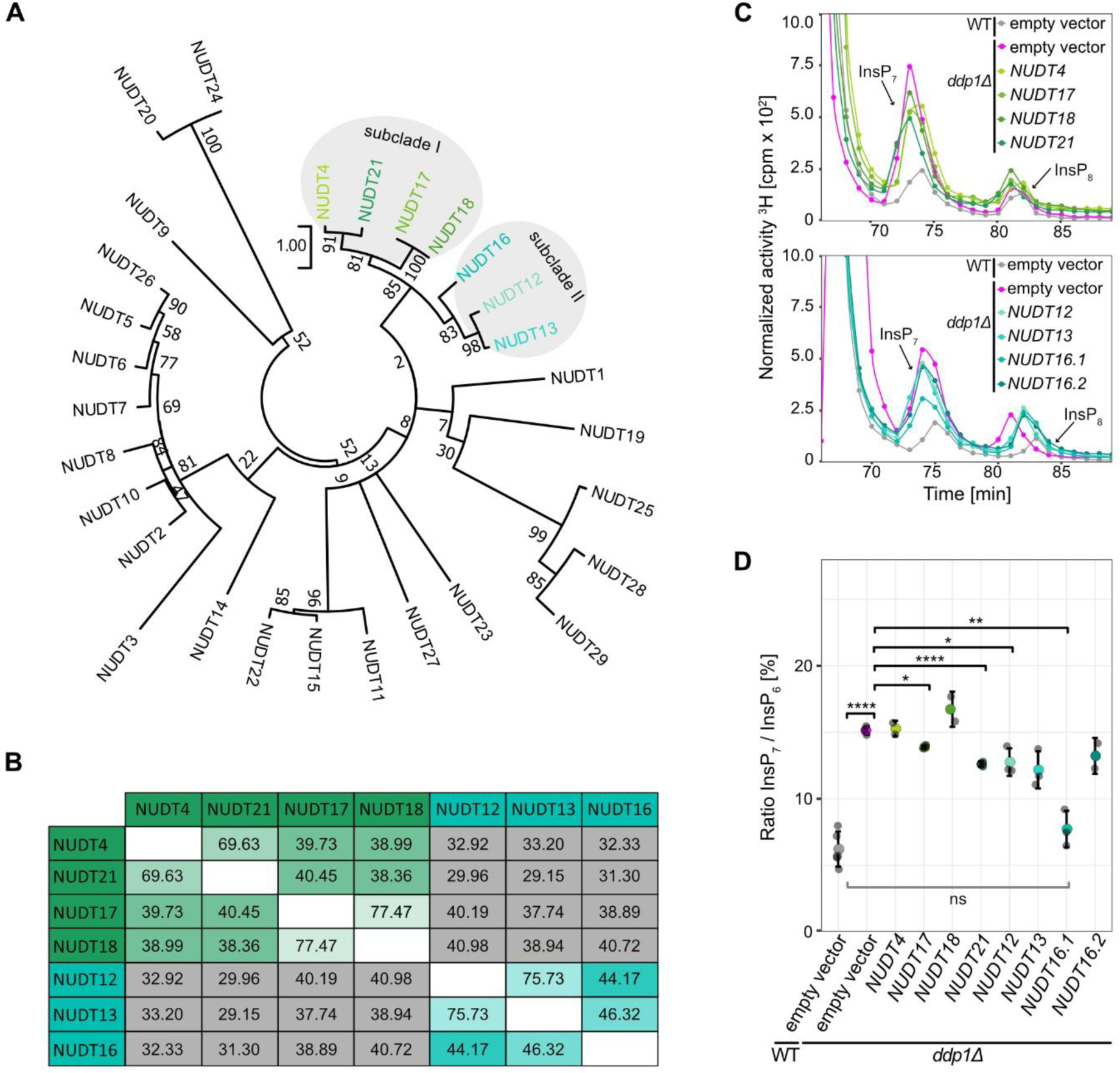
Identified Arabidopsis NUDT hydrolases group in two subclades and heterologous expression partially complements *ddp1Δ*-associated defects in InsP7 levels in yeast. Phylogenetic tree of 29 Arabidopsis NUDT hydrolases **(A)**. The tree was created with MEGA11 and the Jones-Taylor-Thornton method with 500 bootstrap replications. Arabidopsis NUDT4, -17, -18, and -21 can be grouped as a subclade of the phylogenetic tree (subclade I). NUDT12, -13, and -16 can also be grouped as a subclade (subclade II) and together with subclade I as one clade in the phylogenetic tree. **(B)** Pairwise comparison analysis of Arabidopsis subclade I (green) and subclade II (turquoise) NUDTs. Values in the matrix refer to percent identity between respective protein pairs generated in CLC Main Workbench 23.0.3. Color gradients indicate higher (lighter tones) or lower (darker tones) degrees of protein identity. Green and turquoise boxes highlight comparisons within subclades I and II, respectively, while grey boxes compare proteins from different subclades. **(C)** SAX-HPLC profiles of radiolabeled *ddp1Δ* yeast transformed with either empty vector (pAG425GPD-*ccdB*), or pAG425GPD carrying the respective *NUDT* gene. Depicted are representative SAX-HPLC runs of one experiment. The experiment was repeated with similar results (n = 3 or 4, except for NUDT4, NUDT18, NUDT16.2, n = 2). Combined data are shown in Figure 1B. (D) Relative amounts of InsP7 of wild-type (WT) and *ddp1Δ* yeast transformed with empty vector, and *ddp1Δ* transformed with pAG425GPD carrying the *NUDT* genes are shown as InsP7/InsP6 ratios. InsP6 and InsP7 levels were determined by SAX-HPLC analyses and data were processed with R. Data represents mean ± SD. Asterisks indicate values that are significantly different from *ddp1Δ* according to Student’s *t* test (*p* < 0.05 (*); *p* < 0.01 (**); *p* < 0.001 (***); *p* < 0.0001 (****)).

The physiological roles of these hydrolases and their potential substrates were unknown at the onset of this work. We therefore decided to systematically investigate all seven NUDT hydrolases in this clade for their possible involvement in PP-InsP metabolism, including those not directly recovered in the pull-down assay.

### Heterologous expression of Arabidopsis subclade I and II NUDTs partially complement yeast *ddp1Δ* defects

Considering that NUDTs share homology with yeast Ddp1 and mammalian DIPP proteins, all of which display pyrophosphatase activity against 1-InsP_7_, we investigated whether heterologous expression of subclade I and II NUDTs rescues the metabolic defects of the yeast *ddp1Δ* mutant. This mutant strongly accumulates 1-InsP_7_ as compared to its isogenic WT strain (Lonetti et al. 2011) (Figure 1C and D). SAX-HPLC analysis of extracts from [³H]-*myo*-inositol-labeled yeast transformants showed that, with the exception of *NUDT4* and *NUDT18*, expression of all subclade I and II NUDTs partially restored InsP_7_ degradation in the *ddp1Δ* background (Figure 1C and D). The lack of complementation by *NUDT4* may be in part explained by its low expression detected in *ddp1Δ* protein extracts (Figure S2).

Because plants likely contain at least one form of 4/6-InsP_7_ and one form of 1/3-InsP_7_, whose respective enantiomer identities remain unresolved, we resorted to *in vitro* assays to further investigate the substrate specificities of subclade I and II NUDTs.

### *In vitro* biochemical analyses reveal distinct and highly specific PP-InsP pyrophosphatase activities for Arabidopsis subclade I and II NUDT proteins

To biochemically characterize Arabidopsis NUDT hydrolases, we used chemically synthesized, enantiomerically pure InsP_7_ and InsP_8_ isomers (Capolicchio et al. 2013, 2014). Recombinant NUDT proteins were expressed with an N-terminal hexahistidine (His_6_) maltose-binding protein (MBP) tag, and a His_6_-MBP fusion alone was used as a negative control. *In vitro* reactions were carried out with InsP_6_ or different InsP_7_ isomers in the presence of Mg^2+^ (Figure 2A, Figure S3). PAGE analysis revealed that subclade I hydrolases (NUDT4, NUDT17, NUDT18, and NUDT21) catalyzed nearly complete hydrolysis of 4-InsP_7_ (Figure 2A). The resulting product co-migrated with InsP_6_ on PAGE and was confirmed by CE-ESI-MS analysis, using [^13^C_6_] InsP_6_ as a standard, to match the mass and migration time of InsP_6_ (Figure S4).

**Figure 2:**
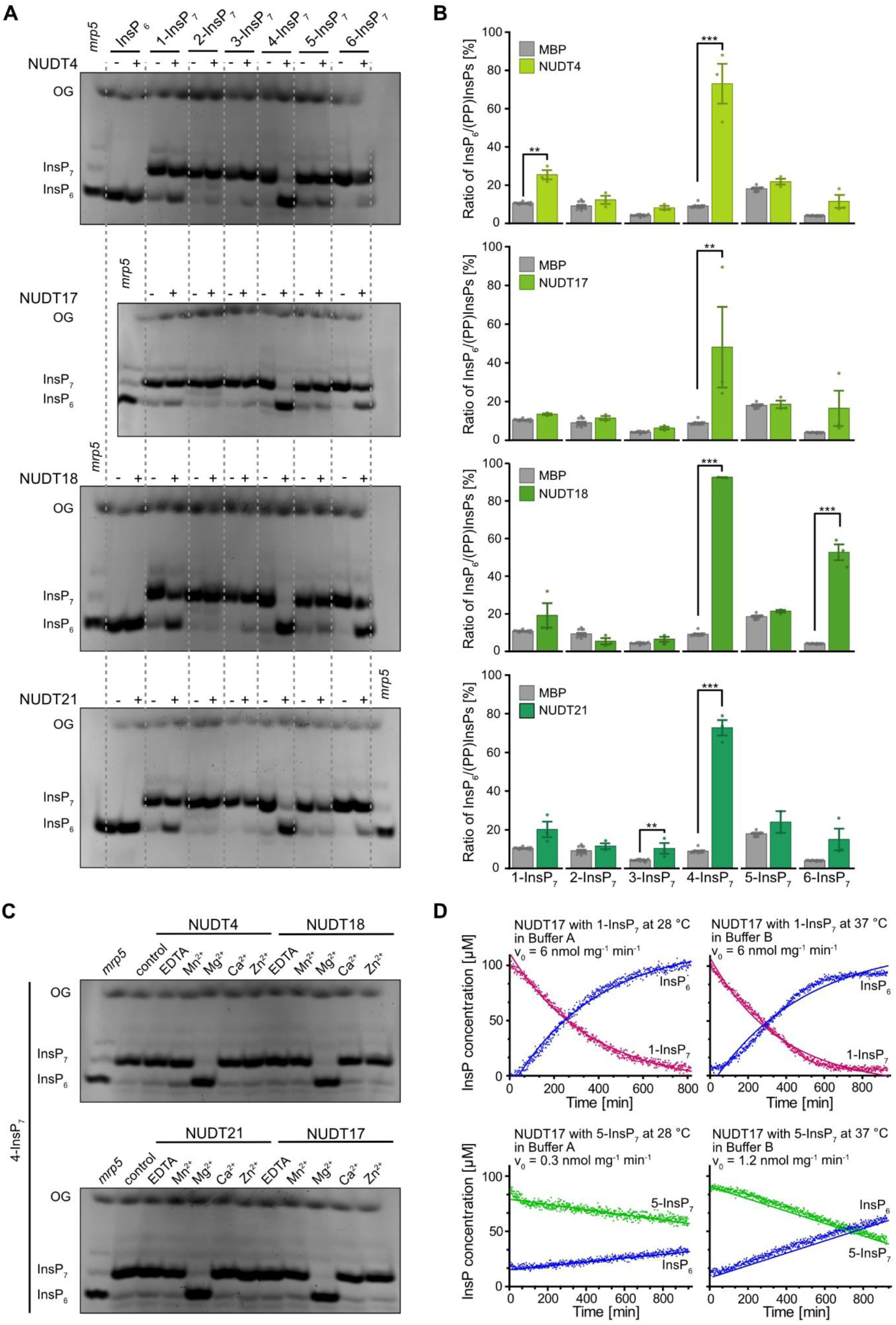
Subclade I NUDTs display Mg^2+^-dependent PP-InsP phosphatase activity with high specificity for 4-InsP7 *in vitro.* Recombinant His6-MBP-tagged NUDT4, NUDT17, NUDT18 or NUDT21 was incubated with 0.33 mM InsP7 and (**A** and **B**) 1 mM MgCl2 or (**C**) either 1 mM EDTA, MnCl2, MgCl2, CaCl2, or ZnCl2 at 28 °C (indicated with the plus symbol). His8-MBP served as a negative control (indicated with the minus symbol). After 1 h, the reaction products were (**A** and **C**) separated by 33 % PAGE, and visualized by toluidine blue. A TiO2-purified Arabidopsis *mrp5* seed extract was used as a marker for InsP6, InsP7 and InsP8. (B) For CE-ESI-MS analyses, samples were spiked with an isotopic standards mixture ([^13^C6] 1,5-InsP8, [^13^C6] 5-InsP7, [^13^C6] 1-InsP7, [^13^C6] InsP6, [^13^C6] 2-OH InsP5). Data are presented as relative ratios of InsP6 to all measured InsP/PP-InsPs and represent mean ± SEM (n ≥ 11 for MBP and n = 3 for NUDT samples, with the exception of sample NUDT21 + 5-InsP7, where only 2 samples were available due to CE-ESI-MS measurement issues). Representative extracted-ion electropherograms are shown in Figure S3. Asterisks indicate values that are significantly different determined by a Dunnett’s test against MBP with Šidák correction k = 4 (*p* ≤ 0.05 (*); *p* ≤ 0.01 (**); *p* ≤ 0.001 (***)). OG: orange G. The enzymatic activity of NUDT17 was also assessed using NMR spectroscopy (**D**) with [¹³C6]1-InsP7 and [¹³C6]5-InsP7 as substrates. Time-course assays were conducted in either buffer A (50 mM HEPES, pH 7.0; 10 mM NaCl; 0.1% (v/v) β-mercaptoethanol; 1 mM MgCl2; 5% (v/v) glycerol; and 0.2 mg/mL BSA) at 28 °C or buffer B (50 mM HEPES, pH 7.3; 150 mM NaCl; 1 mM DTT; 0.5 mM MgCl2; and 0.2 mg/mL BSA) at 37 °C. For reactions displaying a linear progression, rates were determined via linear regression. In cases where the progress curve exhibited a hyperbolic shape, nonlinear regression was applied, and reaction velocities were calculated from the first derivative of a one-phase decay model. All reaction rates, whether derived from linear or nonlinear fits, were normalized to the enzyme’s mass concentration.

Consequently, CE-ESI-MS quantification showed a significant increase of InsP_6_ from 4-InsP_7_ in reactions with subclade I NUDT hydrolases compared to the negative control (Figure 2B). Notably, there was also a robust hydrolysis of 6-InsP_7_ in the presence of NUDT18 (Figure 2A and B) with minor activities of NUDT4 against 1-InsP_7_ and of NUDT21 against 3-InsP_7_ (Figure 2A and B). In contrast, none of the subclade I NUDTs hydrolyzed InsP_6_ or 5-InsP_7_ under these conditions (Figure 2A and B, Figure S3).

To test the cation dependence of 4-InsP_7_ hydrolysis, we substituted Mg^2+^ with alternative divalent cations (Ca^2+^, Zn^2+^, or Mn^2+^) or removed metal ions using EDTA. No hydrolysis was observed under any of these conditions (Figure 2C), indicating that Mg^2+^ is essential for subclade I-mediated 4-InsP_7_ hydrolysis.

To further quantify enzyme activities, we performed pseudo-2D spin-echo difference NMR assays using NUDT17. These assays are limited to [^13^C]-labeled PP-InsP isomers, which are not yet available for all substrates. Reactions were performed under our established low-salt conditions (10 mM NaCl, 28 °C), as well as under high-salt conditions (150 mM NaCl, 37 °C) used by Laurent et al. (2024) for studying NUDT17 and PFA-DSP1. In both conditions, NUDT17 showed consistent but low activity toward 1-InsP_7_ (6 nmol·min^-1^·mg^-1^; Figure 2D). In contrast, activity against 5-InsP_7_ was substantially lower, being reduced by 20-fold under low-salt conditions (0.3 nmol·min^-1^·mg^-1^) and by ∼5-fold under high-salt conditions (1.2 nmol·min^-1^·mg^-1^), suggesting limited catalytic efficiency toward this substrate. In fact, the low hydrolysis rates obtained with 5-IP_7_ resemble the rate of spontaneous (non-enzymatic) hydrolysis, further underscoring the weak enzymatic activity of NUDT17 toward this isomer. Together, these data indicate that 4-InsP_7_ is the preferred substrate of NUDT17, while 1- and 5-InsP_7_ are likely minor or physiologically less relevant targets.

In contrast to subclade I NUDTs, the four hydrolases of subclade II (NUDT12, NUDT13, NUDT16.1 and NUDT16.2) displayed a strong preference for 3-InsP_7_. The product co-migrated with InsP_6_ by PAGE (Figure 3A) and was confirmed by CE-ESI-MS (Figure 3B). Notably, NUDT16.2 displayed comparatively low activity, requiring a 26-fold higher protein concentration than NUDT16.1 to achieve comparable hydrolysis levels (Figure S5). Other PP- InsPs were poor substrates: NUDT12 showed only minor activity against all tested PP-InsPs except 5-InsP_7_; NUDT13 and NUDT16.1 displayed weak activity toward 1-, 4-, and 6-InsP_7_; and NUDT16.2 hydrolyzed 1- and 6-InsP_7_ to a limited extent (Figure 3A and B; Figure S6). Like subclade I proteins, subclade II hydrolases required Mg^2+^ for activity, and no hydrolysis was observed with Ca^2+^, Zn^2+^, Mn^2+^, or EDTA (Figure 3C). It is important to note that the *in vitro* assay results shown in Figure 2B and Figure 3B represent the relative ratios of InsP_6_ to all detected InsP/PP-InsP species in order to take into account the background InsP_6_ signal originating from InsP_7_ and InsP_8_ synthesis. However, we cannot entirely exclude the possibility that residual InsP_6_ may influence enzyme activity, either through competitive inhibition or by altering binding dynamics.

**Figure 3:**
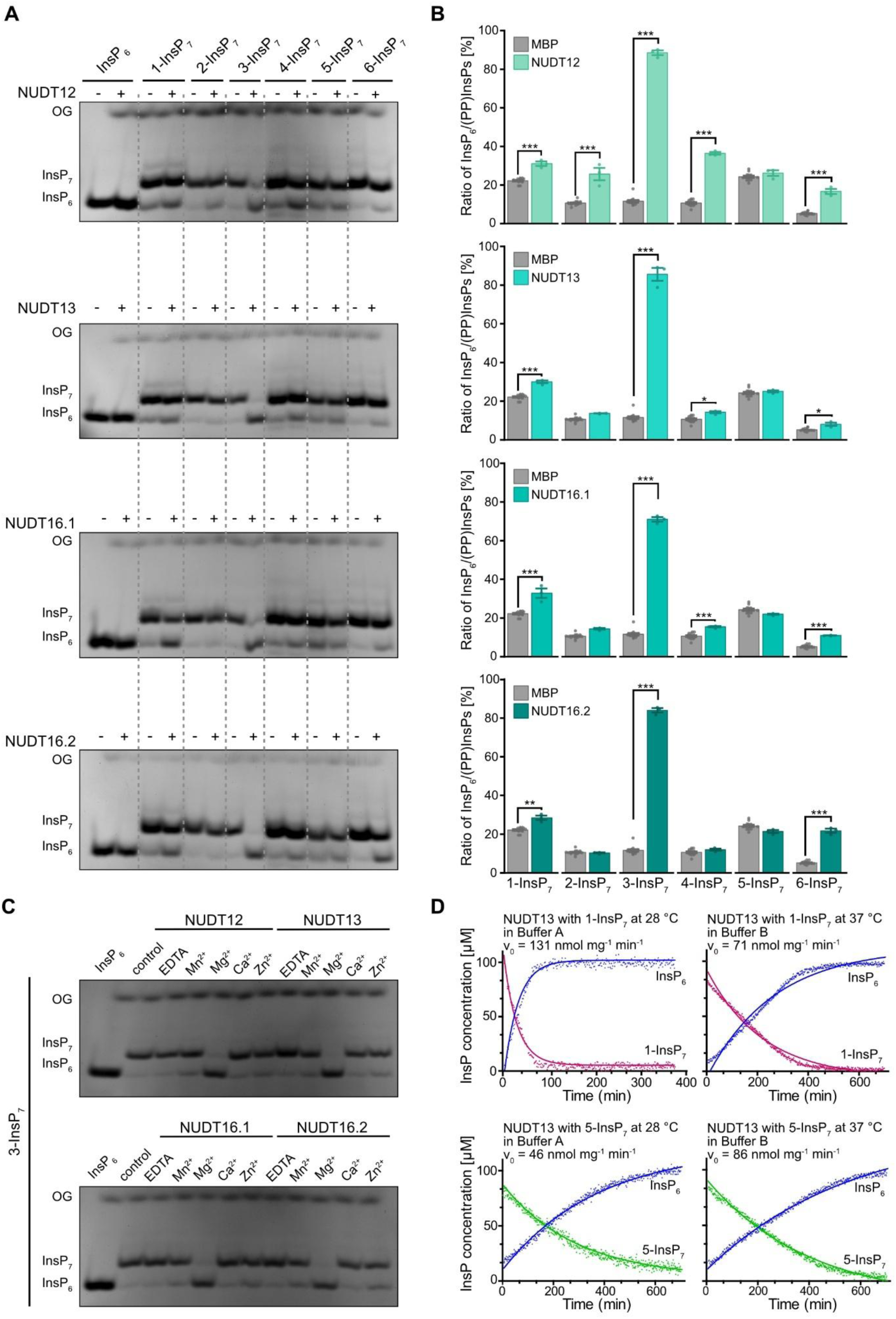
Subclade II NUDTs display Mg^2+^-dependent PP-InsP phosphatase activity with high specificity for 3-InsP7 *in vitro.* Recombinant His6-MBP-tagged NUDT12, NUDT13, NUDT16.1 or NUDT16.2 was incubated with 0.33 mM InsP7 and (**A** and **B**) 1 mM MgCl2 or (**C**) either 1 mM EDTA, MnCl2, MgCl2, CaCl2, or ZnCl2 at 28 °C (indicated with the plus symbol). His8-MBP served as a negative control (indicated with the minus symbol). After 1 h, the reaction products were (**A** and **C**) separated by 33 % PAGE and visualized by toluidine blue or (**B**) spiked with isotopic standards mixture ([^13^C6] 1,5-InsP8, [^13^C6] 5-InsP7, [^13^C6] 1-InsP7, [^13^C6] InsP6, [^13^C6] 2-OH InsP5) for CE-ESI-MS analyses. Data represent mean ± SEM (n = 12 for MBP and n = 3 for NUDT samples). Representative extracted-ion electropherograms are shown in Figure S5. Asterisks indicate values that are significantly different determined by a Dunnett’s test against MBP with Šidák correction k = 4 (*p* ≤ 0.05 (*); *p* ≤ 0.01 (**); *p* ≤ 0.001 (***)). OG: orange G. Note that NUDT16.2 had a 16-times higher concentration (16 µM) than NUDT16.1 (0.6 µM). Time-course reactions with [^13^C6]1-InsP7 and [^13^C6]5-InsP7 as substrates were carried out in either buffer A (50 mM HEPES, pH 7.0; 10 mM NaCl; 0.1% (v/v) β-mercaptoethanol; 1 mM MgCl2; 5 % (v/v) glycerol; 0.2 mg/mL BSA) at 28 °C or buffer B (50 mM HEPES, pH 7.3; 150 mM NaCl; 1 mM DTT; 0.5 mM MgCl2; 0.2 mg/mL BSA) at 37 °C and monitored by NMR spectroscopy (**D**). Linear progress curves were analyzed using linear regression. For hyperbolic curves, nonlinear regression based on a one-phase decay model was applied, with initial velocities calculated from its first derivative. All rates were normalized to the mass concentration of NUDT13.

As for subclade I, we used pseudo-2D spin-echo difference NMR to assess NUDT13 activity. Under low-salt conditions at 28 °C, NUDT13 exhibited modest activity toward 1-InsP_7_ (131 nmol·min^-1^·mg^-1^) and ∼3-fold lower activity toward 5-InsP₇ (Figure 3D), consistent with PAGE-based endpoint measurements. Under high-salt conditions at 37 °C, activities toward 1-InsP_7_ and 5-InsP_7_ were comparable (71 vs. 86 nmol·min^-1^·mg^-1^ respectively). However, overall activity levels remained moderate, reinforcing the conclusion that 5-InsP_7_ and 1-InsP_7_ are not preferred substrates of NUDT13.

### Subclade II NUDT hydrolases display β-phosphate specificity also with 1,5-InsP_8_ and 3,5-InsP_8_, *in vitro*

We then tested whether subclade I and II NUDTs can also hydrolyze the enantiomers of InsP_8_ 1,5-InsP_8_ and 3,5-InsP_8_ in the presence of Mg^2+^ *in vitro* (Figure 4A-D, Figure S7, Figure S8). Subclade I NUDTs, which specifically hydrolyze the 4-β-phosphate of InsP_7_, exhibited only limited hydrolytic activity toward 1,5-InsP_8_, which was observed for NUDT4, NUDT18, and NUDT21 (Figure 4A and B). Additionally, NUDT4 displayed minor activity against 3,5-InsP_8_. In contrast, subclade II hydrolases (NUDT12, NUDT13, and NUDT16) robustly hydrolyzed the 3-β-phosphate of 3,5-InsP_8_. This activity was enantiomer-specific for NUDT12 and NUDT13, while NUDT16 (both splice forms) also hydrolyzed 1,5-InsP_8_ to 5-InsP_7_ to a detectable extent (Figure 4C and D).

**Figure 4:**
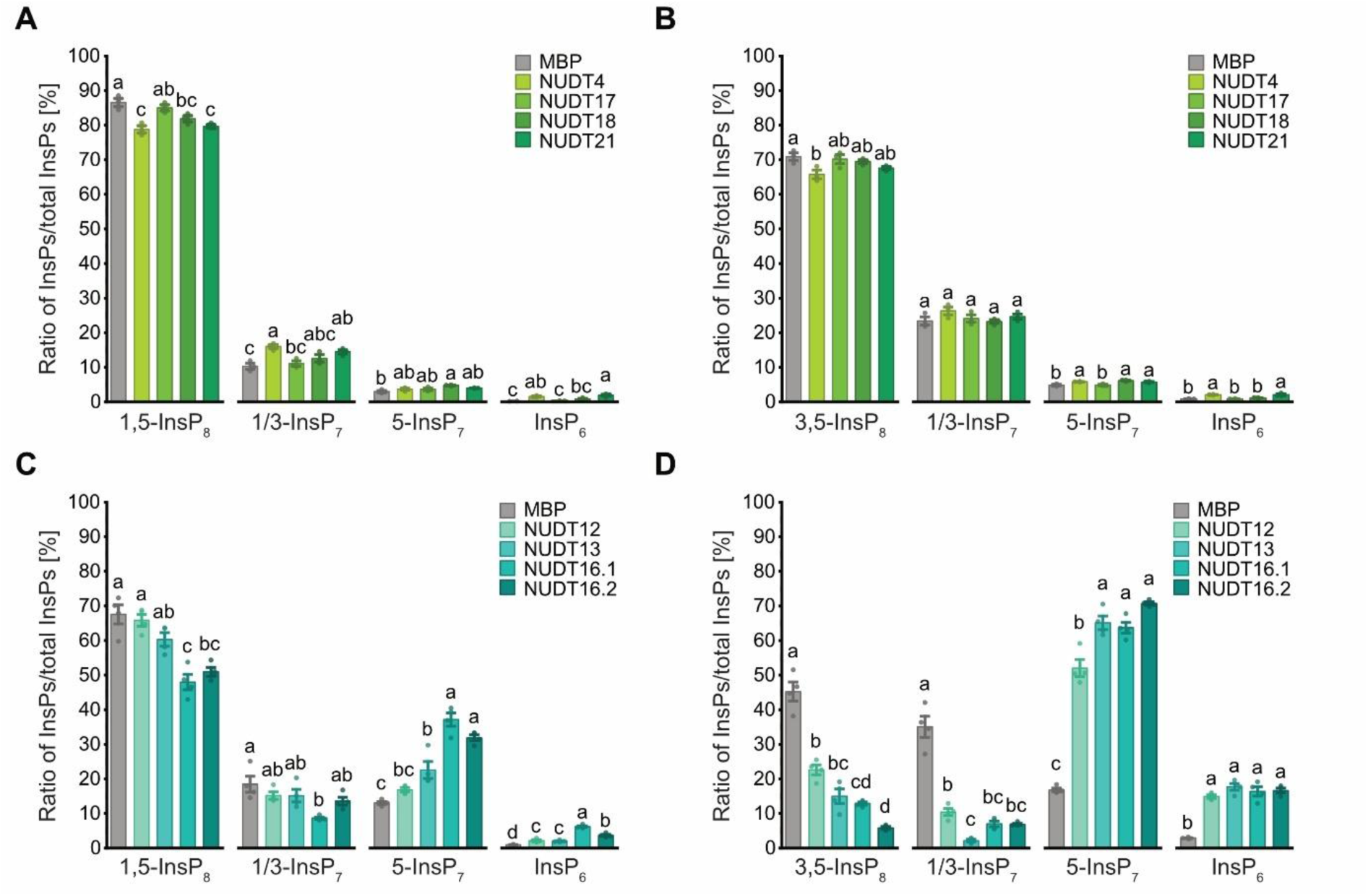
Arabidopsis NUDT hydrolases display differential hydrolytic activities against 1,5-InsP8 and 3,5-InsP8 *in vitro*. Recombinant His6-MBP-tagged (**A** and **B**) NUDT4, NUDT17, NUDT18, NUDT21, (**C** and **D**) NUDT12, NUDT13, NUDT16.1 or NUDT16.2 protein was incubated with 0.33 mM (**A** and **C**) 1,5-InsP8 or (B and D) 3,5-InsP8 and 1 mM MgCl2 at 28 °C. His8-MBP served as a negative control. After 1 h, the reaction products were spiked with isotopic standards mixture ([^13^C6] 1,5-InsP8, [^13^C6] 5-InsP7, [^13^C6] 1-InsP7, [^13^C6] InsP6, [^13^C6] 2-OH InsP5) and subjected to CE-ESI-MS analyses. Data represent mean ± SEM with (**A** and **B**) n = 3 and (**C** and **D**) n = 4. Representative extracted-ion electropherograms are shown in Figure S6 and Figure S7. Different letters indicate values that are significantly different determined with one-way ANOVA followed by Dunn-Šidák test with a significance level of 0.05. The peak area of 1/2/3-InsP7 and of 4/5/6-InsP7 was not further resolved but, since 1,5-InsP8 and 3,5-InsP8 isomers were used in that reaction, we conclude that the 1/2/3-InsP7 peak area represents 1/3-InsP7 and the 4/5/6-InsP7 peak area represents 5-InsP7, respectively.

### At high protein concentrations, NUDT hydrolases lose some substrate specificity

We next investigated whether increasing protein concentration and doubling the incubation time of our *in vitro* assays would result in a loss of substrate specificity. Under these modified conditions, overall substrate preferences remained largely consistent with our earlier findings (Figure S5, Figure S9). However, additional, lower-level hydrolytic activities were observed.

For instance, all subclade I NUDTs partially hydrolyzed 5-InsP_7_, as well as both 1,5 and 3,5-InsP_8_ (Figure S9). Likewise, subclade II members NUDT12, NUDT13 and NUDT16.1 retained robust activity toward 3-InsP_7_ but also showed low-level activity against other InsP_7_ isomers, including those pyrophosphorylated at positions 1, 2, 4, and 6. Weak hydrolytic activity against 5-InsP_7_ was also observed (Figure S5).

To assess potential off-target activity in more distantly related NUDIX proteins, we included NUDT7, previously shown to act as an ADP-ribose/NADH phosphohydrolase (Ogawa et al. 2005). As expected, NUDT7 did not hydrolyze any PP-InsP substrates, even at high protein concentrations (∼7 µM) (Figure S5), indicating that the observed pyrophosphatase activity is specific to the subclade I and II enzymes.

Collectively, these findings reveal that subclade I and II NUDT hydrolases exhibit distinct substrate specificities, with a preference for 4-InsP_7_ and 3-InsP_7_, respectively, and limited activity toward other PP-InsPs, especially under more physiologically relevant conditions.

### Higher order mutants of subclade II NUDTs display an increase in 1/3-InsP_7_ and 5-InsP_7_

To investigate the physiological role of subclade I and II NUDT hydrolases *in planta*, we established single and higher order loss-of-function mutants in *Arabidopsis thaliana.* The single T-DNA insertion lines *nudt17-1* and *nudt18-1*, as well as two independent T-DNA insertion lines of *nudt4* (*nudt4-1* and *nudt4-2*) and *nudt21* (*nudt21-1* and *nudt21-2*) showed no significant difference in PP-InsP levels in comparison to the WT when grown hydroponically under optimal P_i_ conditions (Figure S10). Similarly, a *nudt21-1 nudt4-1* double mutant also exhibited no detectable change in PP-InsP profiles (Figure S10).

To overcome a possible functional redundancy of NUDT hydrolases, we generated higher-order mutants by targeting all NUDT genes from either subclade I (*NUDT4*, *NUDT17*, *NUDT18* and *NUDT21*) or subclade II (*NUDT12*, *NUDT13* and *NUDT16*) using CRISPR/Cas9. For each gene, guide RNAs were designed to target sites near the start codon or within the conserved 23-amino acid NUDT domain (Figure S11A, Figure S12A). The resulting higher-order mutants *nudt4/17/18/21* and *nudt12/13/16* harbor mutations in all four and three targeted NUDT hydrolases, respectively (Figure S11B, Figure S12B). As depicted in Figure S11C, the resulting mutations in *nudt4/17/18/21* lead to a frameshift in all targeted open reading frames, except for line 1, where an in-frame deletion resulted in a truncated NUDT21 lacking residues 32-74. Most edits resulted from single base pair mutation or larger deletions (Figure S11C). In the *nudt12/13/16* triple mutant, *NUDT13* and *NUDT16* carried frameshift mutations with premature stop codons, while the start codon of *NUDT12* was disrupted (Figure S12C).

None of the higher order subclade I or II *nudt* lines displayed any obvious developmental phenotype under standard growth conditions (Figure 5A). We first analyzed [^3^H]-*myo*-inositol-labeled extracts from WT and the higher order mutants, *nudt4/17/18/21* and *nudt12/13/16* using SAX-HPLC. In *nudt4/17/18/21* (line 2), no consistent differences in any InsP or PP-InsP species were detected (Figure 5B). In contrast, *nudt12/13/16* plants displayed a significant increase in InsP_7_ (Figure 5C). Because SAX-HPLC analyses cannot resolve between different InsP_7_ isomers we employed CE-ESI-MS analyses of Nb_2_O_5_-purified InsPs from plants grown on ½ MS medium.

**Figure 5:**
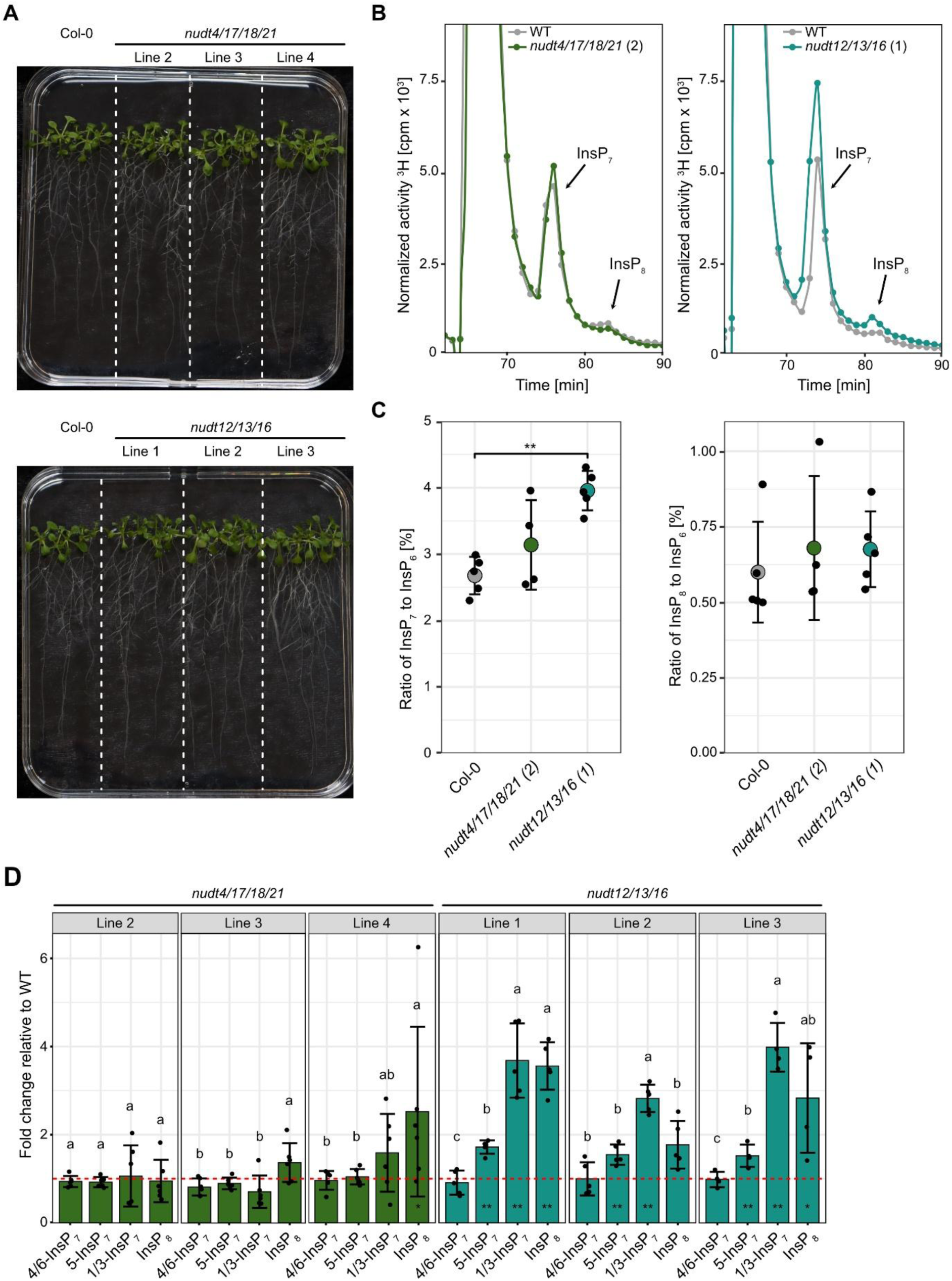
Higher order subclade II *nudt* mutant lines display increased 1/3-InsP7 and 5-InsP7 levels. (**A**) Representative images of 2-week-old wild-type (WT) plants and independent higher-order *nudt* mutant lines from subclades I and II grown on ½ MS agar plates supplemented with sucrose. (**B**) SAX-HPLC profiles of 2- weeks old WT and subclade I and subclade II *nudt* mutant lines. Lines were grown in liquid ½ MS medium supplemented with sucrose. Labeling with [^3^H]-*myo*-inositol was done for 5 days. Depicted are representative SAX-HPLC runs of one line for each subclade. The experiment was repeated with similar results (n = 4 or 5). Combined data are shown in Figure 5C. (**C**) Relative amounts of InsP7 and InsP8 of WT, subclade I and subclade II NUDT mutant lines are shown as InsP7/InsP6 or InsP8/InsP6 ratios. InsP6, InsP7 and InsP8 levels were determined by SAX-HPLC analyses and data were processed with R. Asterisks indicate significantly different InsP7/InsP6 or InsP8/InsP6 ratios compared to WT determined with Wilcoxon test for unpaired samples (*p* ≤ 0.01 (**)). (**D**) Fold changes in InsP8 and various InsP7 isomers, normalized to InsP6, in subclade I and subclade II *nudt* mutant lines relative to wild-type (WT), with the red dotted line indicating the WT reference (fold change = 1). Plants were grown on ½ MS agar plates supplemented with sucrose for 2 weeks. Asterisks denote statistically significant differences from WT based on a Wilcoxon test for unpaired samples (\**p* ≤ 0.05; \*\**p* ≤ 0.01; \*\*\**p ≤ 0.001*). Compact letters indicate statistical groupings determined by pairwise Wilcoxon tests for paired samples; InsP fold changes sharing the same letter are not significantly different at α = 0.05.

Under our standard growth conditions, we did not observe changes in any inositol pyrophosphate species that were consistent in all three independent subclade I *nudt4/17/18/21* lines (Figure 5D, left panels). In contrast, all subclade II *nudt12/13/16* lines showed a robust ∼2.5-fold increase in 1/3-InsP_7_ (Figure 5D, right panels), indicating that the *in vitro* hydrolytic activity against the 1/3-PP bond is of physiological relevance *in planta*. Additionally, we observed an ∼1.5-fold increase in 5-InsP_7_ and, for line 1 and line 3, a significant increase of ∼2-fold in InsP_8_. Furthermore, all lines showed a minor but significant reduction in InsP_6_ (Supplementary File 2).

### Transient expression of subclade I NUDT hydrolases in *Nicotiana benthamiana* reveals PP-InsP pyrophosphatase activity and induction of phosphate starvation responses *in vivo*

Since the *nudt4/17/18/21* lines did not show consistent alterations in PP-InsPs, possibly due to redundancy with other enzymatic activities, we further explored the potential PP-InsP pyrophosphatase activity of NUDTs by heterologous transient expression in *N. benthamiana* leaves. To this end, we generated constructs encoding translational fusions of NUDTs from both subclades with a C-terminal 4xMyc epitope tag, under control of the strong viral CaMV *35S* promoter.

In agreement with our *in vitro* findings (Figure 2), transient expression of *NUDT17*, *-18*, and *- 21* significantly reduced 4/6-InsP_7_ in *N. benthamiana* leaves, as revealed by CE-ESI-MS analyses (Figure 6A, Figure S13). In addition, expression of all subclade I *NUDTs* also resulted in reduced 5-InsP_7_ levels, consistent with our *in vitro* data showing 5-InsP_7_ hydrolysis at higher protein concentrations (Figure 6B, Figure S9). Similarly, transient expression of subclade II *NUDTs*, with the exception of *NUDT13*, led to reduced levels of 4/6-InsP_7_ and 5-InsP_7_ in *N. benthamiana* (Figure 6A and B, Figure S13, Supplementary File 2). Quantification of InsP_8_ and 1/3-InsP_7_ species was not possible in samples from this transient expression system, as concentrations were close to or below the detection limit of CE-ESI-MS measurements.

**Figure 6:**
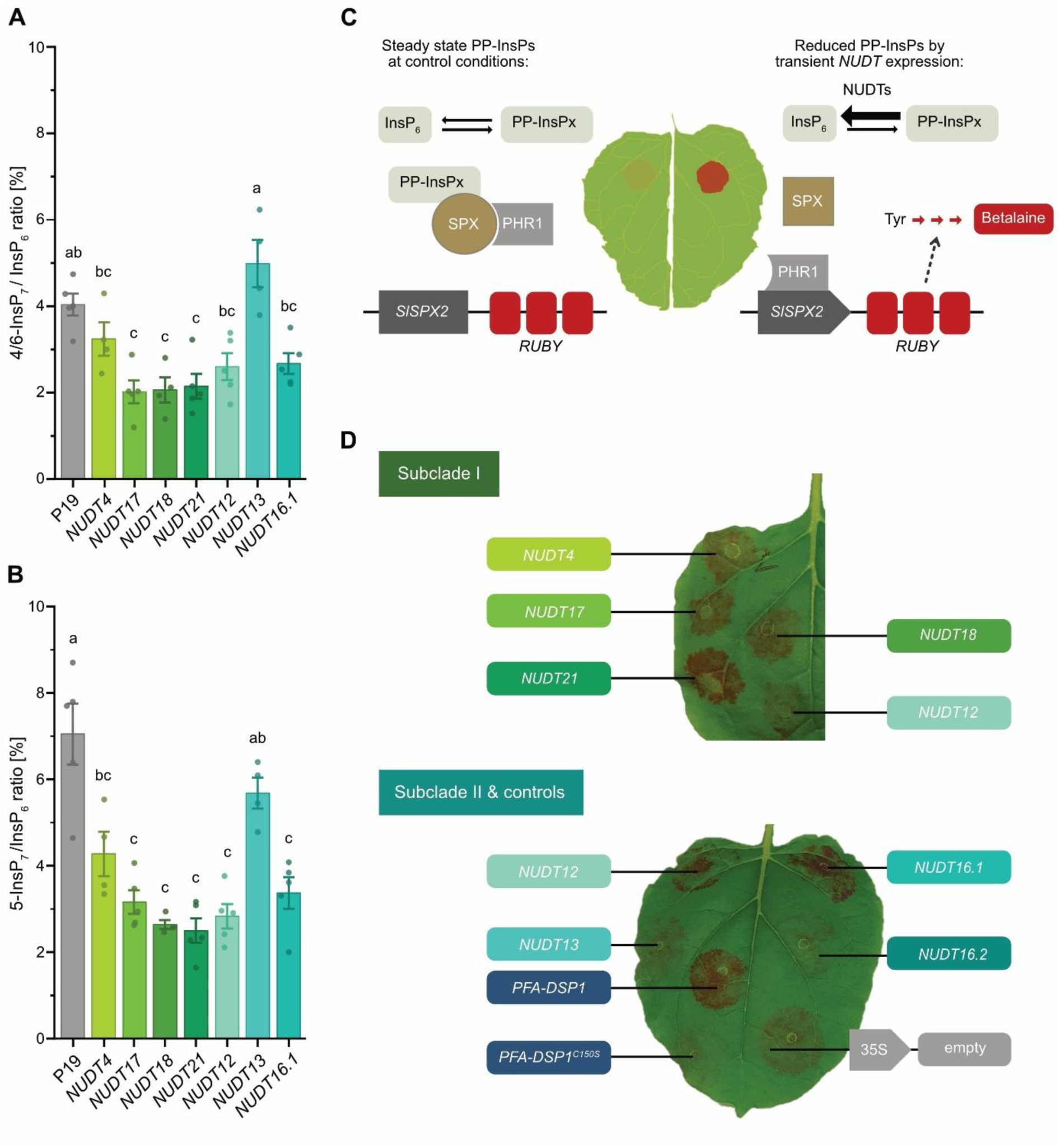
Transient expression of NUDTs in *N. benthamiana* reveals PP-InsP activity and regulation of phosphate starvation responses. (**A** and **B**) Heterologous expression of subclade I and II NUDTs results in 4/6-PP-InsP7 and 5-PP-InsP7 turnover *in planta* and induces local PSR. The silencing inhibitor P19 alone or together with NUDTs were transiently expressed in 6-weeks old *N. benthamiana* leaves. 2-3 days post infiltration (dpi) (PP-)InsPs were purified with Nb2O5 beads, spiked with isotopic standards mixture ([^13^C6] 1,5-InsP8, [^13^C6] 5-InsP7, [^18^O2] 4-InsP7, [^13^C6] 1-InsP7, [^13^C6] InsP6, [^13^C6] 2-OH InsP5) and subjected to CE-ESI-MS analyses. Data represent mean ± SEM (n ≥ 4). Different letters indicate values that are significantly different determined with one-way ANOVA followed by Dunn-Šidák test with a significance level of 0.05. Representative extracted-ion electropherograms are shown in Figure S12. (C) Schematic representation of PSR reporter assay in *N. benthamiana* leaves, in which a reduced PP-InsP concentration enables PHR1 to bind a P1BS binding site in the *SlSPX2* promoter (Solyc12g009480; LOC101257836) driving the expression (dotted arrow) of a *RUBY* reporter cassette (encoding three different proteins CYP76AD1, DODA and a glycosyltransferase) that catalyze the stepwise synthesis (red arrows) of a red betalain pigment. (D) Transient co-expression of the *RUBY* reporter with subclade I or subclade II *NUDT* hydrolase genes under the transcriptional control of the viral CaMV *35S* promoter. Co-expression with *PFA-DSP1* served as a positive control while co-expression with *PFA-DSP1^C150S^* encoding the catalytic inactive protein or an empty vector served as negative controls. The picture was taken 2 dpi (upper leaf) or 3 dpi (bottom leaf). Note that subclade I *NUDT12* was expressed in both leaves for comparison. Three independent replicates are shown in Figure S13.

Expression data extracted from the ePlant browser platform (Kilian et al. 2007) indicate that *NUDT21* and *NUDT17* are locally and transiently upregulated upon wounding. Given the ^k^nown role o^f^ 5-InsP_7_ ^i^n PS^R^, we tested whether ectopic expression of subclade I and II *NUDTs* could activate a local PSR *in planta*. To this end, we co-expressed a *RUBY* reporter (He et al. 2020) under transcriptional control of a *Solanum lycopersicum SPX2* promoter fragment with *NUDT* hydrolases in *N. benthamiana* leaves (Figure 6C). Heterologous expression of Arabidopsis *PFA-DSP1*, which encodes a *bona fide* 5PP-InsP pyrophosphatase hydrolyzing the 5-β-phosphate of 5-InsP_7_, as well as 1,5-InsP_8_ and 3,5-InsP_8_ (Gaugler et al. 2022), served as a positive control (Figure 6D, Figure S14).

As expected, PFA-DSP1 induced strong RUBY reporter activity, while its catalytically inactive variant PFA-DSP1^C150S^ (Gaugler et al. 2022), failed to do so (Figure 6D, Figure S14), confirming that PSR induction was dependent on the PFA-DSP1 enzymatic activity. Notably, heterologous expression of all subclade I *NUDTs* caused robust red staining, reporting local activation of PSR (Figure 6D). Among subclade II members, NUDT12 and the active splice variant of NUDT16 (variant 1) also induced red staining, albeit generally with lower intensity (Figure 6D, Figure S14). In contrast, expression of *NUDT13* or the less active splice variant 2 of *NUDT16* failed to activate PSR as compared to the empty vector or PFA-DSP1^C150S^ negative controls (Figure 6D, Figure S14).

### Subclade II NUDTs are involved in P_i_- and Fe-homeostasis

To investigate the physiological consequences of elevated 1/3-InsP_7_ and 5-InsP_7_ levels observed in *nudt12/13/16* lines, we first assessed plant growth and mineral nutrient profiles. Under standard growth conditions, neither the higher-order *nudt4/17/18/21* nor the *nudt12/13/16* lines displayed any obvious developmental phenotype (Figure 5A). However, given that 5-InsP_7_ can bind and activate SPX domain proteins (Wild et al. 2016) and has been proposed to partially compensate for the loss of InsP_8_ in suppressing PSR in Arabidopsis (Riemer et al. 2021), we performed mineral nutrient profiling under P_i_-sufficient conditions.

Inductively coupled plasma optical emission spectroscopy (ICP-OES) analysis of shoots from *nudt12/13/16* plants grown on a peat-based substrate revealed significantly decreased P concentrations in both independent mutant lines compared to WT (Figure 7A). This is consistent with a role of 5-InsP_7_ in PSR regulation and suggests a potential disruption in P_i_ homeostasis. Among macronutrients, no consistent differences were observed. However, strikingly, shoot iron (Fe) concentrations were reduced by approximately 53% in both mutant lines, indicating an additional role for subclade II NUDTs in Fe homeostasis (Figure 7B).

**Figure 7:**
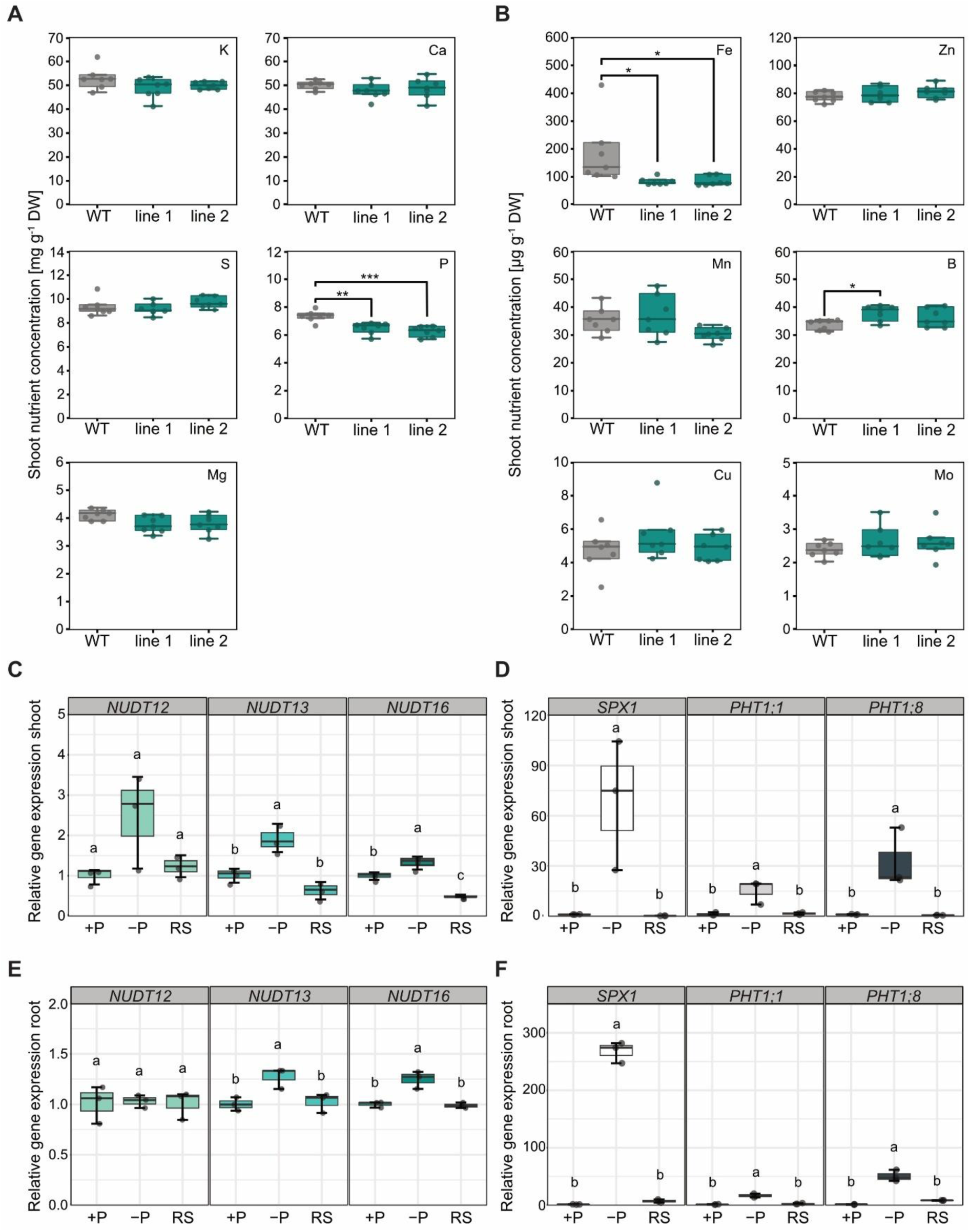
Subclade II NUDTs are involved in P and Fe homeostasis and are upregulated by P-deficiency. (**A** and **B**) Shoot concentration of the indicated mineral macronutrients (**A**) and micronutrients. (**B**) Each line is represented by seven biological replicates (n = 7). Asterisks indicate values that are significantly different determined with one-way ANOVA followed by Dunn-Šidák test (*p* ≤ 0.05 (*); *p* ≤ 0.01 (**); *p* ≤ 0.001 (***)). Plants were grown for 4-5 weeks on peat-based substrate. (**C-F**) Arabidopsis wild-type (WT) plants were grown under three conditions: phosphorus-sufficient (+P), phosphorus-deficient (–P), and phosphorus-deficient followed by 12 hours of phosphate resupply (RS). Gene expression in shoots and roots was analyzed by qPCR. Expression levels of *NUDT12*, *NUDT13*, and *NUDT16* (panels **C** and **E**), and *SPX1*, *PHT1;1*, and *PHT1;8* (panels **D** and **F**) are presented as mean ± SEM (n = 3). Different letters denote statistically significant differences, determined by one-way ANOVA followed by Dunn-Šidák post hoc test (*p* < 0.05).

To determine whether *NUDT12*, *NUDT13*, and *NUDT16* are themselves responsive to changes in P_i_ availability, we performed expression analysis in plants grown hydroponically under P_i_-sufficient and -deficient conditions, followed by P_i_ resupply, as described previously (Riemer et al. 2021). The expression of *NUDT13* and *NUDT16* was significantly upregulated in response to P_i_ deficiency: *NUDT13* showed approximately 2-fold induction in both shoots and roots, while *NUDT16* was induced ∼1.5-fold (Figure 7C and E). *NUDT12* expression remained largely unchanged.

In parallel, we also analyzed the transcript levels of well-characterized PSI marker genes *SPX1*, *PHT1;1*, and *PHT1;8*. As expected, all three PSI markers showed strong induction under P_i_-deficiency, with *SPX1* transcript levels increasing approximately 75-fold in shoots and 250-fold in roots (Figure 7D and F). *PHT1;8* was induced around 30-fold in shoots and 50-fold in roots, while *PHT1;1* showed a more moderate response (Figure 7D and F). Although *NUDT13* and *NUDT16* are not as strongly induced as classical PSI genes, the significant upregulated gene expression under phosphate deficiency still supports a regulation by P_i_ availability.

### Transcriptome analyses suggests subclade II NUDT-targeted PP-InsPs act primarily as negative regulators of gene expression and also modulate PSR-independent functions

To further explore the physiological roles of subclade II NUDTs, and by extension, 1/3-InsP_7_, we performed RNA-seq analyses on shoots from two independent *nudt12/13/16* triple mutant lines and WT plants grown under P_i_-sufficient conditions on a peat-based substrate (Figure 8A). Parallel, CE-ESI-MS analysis confirmed a selective accumulation of 1/3-InsP_7_ in the mutants under these conditions, with no detectable increase in 5-InsP_7_ (Figure 8B), in contrast to plants grown on ½ MS medium (Figure 5D).

**Figure 8:**
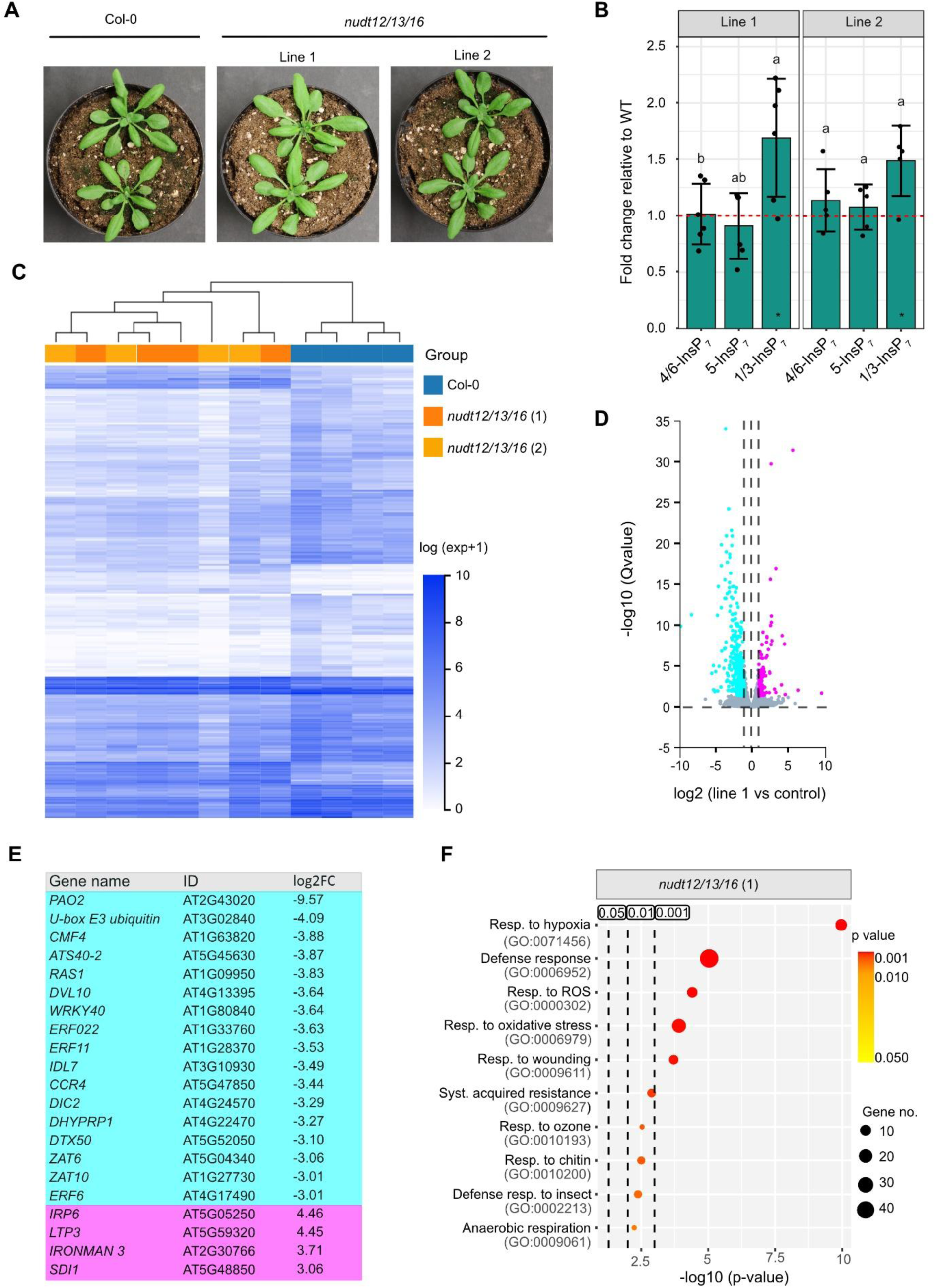
Shoot transcriptome analyses of *nudt12/13/16* triple mutants suggest subclade II NUDT-dependent PP-InsPs act primarily as negative regulators of gene expression. (**A**) Representative images of 3-week-old wild-type (WT) plants and independent higher-order nudt mutant lines from subclade II grown on peat-based substrate. (**B**) CE-ESI-MS analysis of inositol phosphates (InsPs) from plants grown under the same conditions. Fold changes of the indicated InsP7 isomers, normalized to InsP6, are shown for subclade II nudt mutants relative to WT (indicated by the red dotted line). Asterisks denote statistically significant differences from WT (fold change ≠ 1) based on the Wilcoxon test for unpaired samples (\**p* ≤ 0.05; \*\**p* ≤ 0.01; \*\*\**p* ≤ 0.001). Compact letters indicate statistical groupings among InsP isomers, determined by pairwise Wilcoxon tests for unpaired samples. InsPs sharing the same letter do not differ significantly at α = 0.05. (**C**) Clustered Heatmap of differentially expressed genes (DEGs) filtered by Q<0.05 and log2 fold change (|log2FC| > 1). The horizontal axis represents the log2 (expression value + 1) for the samples based on TPM (Transcripts Per Million) expression value, while the vertical axis represents genes. In the default color scheme, dark blue indicates higher expression levels and white indicates lower expression levels. Mutant lines are represented in orange and light orange, and are clustered together, while the control is depicted in blue. Each line is represented by four biological replicates (n = 4). (**D**) Volcano plot comparing triple mutant line 1 to WT. The X-axis represents the log2FC values, and the Y-axis shows -log10-transformed significance values. Magenta dots indicate upregulated DEGs, cyan dots indicate downregulated DEGs, and gray dots indicate non-DEGs. (**E**) The table displays genes that showed strongest differential expression (|log2FC| > 3) of only downregulated and upregulated genes with known annotations. A complete list with all genes in this category of strongest differential expression is shown in Table S3. All plants used in these analyses, including wildtype controls, were grown on a peat-based substrate for 2-3 weeks. (**F**) Shown is a GO enrichment analysis of DEGs with Q < 0.05 and (|log2FC| > 1), based on Biological Processes (BP) for *nudt12/13/16* triple mutant line 1. The X-axis represents the statistical significance of these GO terms in -log10 (p-value), while the Y-axis lists the GO terms with their respective ID. Bubble size indicates the number of DEGs annotated to each GO term, and the graph highlights three significance thresholds: 0.05, 0.01, and 0.001. The color of the bubbles corresponds to the p-value, with red indicating more significant enrichment (Resp.: response, ROS: reactive oxygen species, syst.: systematic, termin.: termination, poly.: polymerase, transc.: transcription).

Across both mutant lines, we identified 256 genes that were differentially expressed compared to WT (Figure 8C, D, and E, Supplementary File 2). Notably, the vast majority (206) were downregulated, suggesting that PP-InsPs targeted by subclade II NUDTs such as 1/3-InsP_7_, may function predominantly as a negative regulator of gene expression. Only 50 genes were upregulated in the mutants.

Among the differentially expressed genes (DEGs), several known to be associated with PSRs, including *PAP1*, *ATL80*, and *ZAT6*, were significantly downregulated (Tables S2 and S3). Additionally, a subset of Fe-responsive genes also showed altered expression (Tables S3 and S4), consistent with the observed changes in Fe accumulation (Figure 7B). However, PSR- and Fe-related transcripts accounted for only a small proportion of total DEGs.

Gene Ontology (GO) enrichment analysis revealed a striking overrepresentation of terms related to defense and associated processes, including responses to wounding, reactive oxygen species (ROS), chitin, insects, and systemic acquired resistance in both independent lines (Figure 8F, Figure S15). Other enriched terms pointed to broader stress-associated processes, such as responses to hypoxia, oxidative stress, ozone, and anaerobic respiration, many of which are also linked to ROS signaling pathways (Figure 8F, Figure S15).

These data suggest that 1/3-InsP_7_, regulated by subclade II NUDTs, plays a broader role beyond PSR, potentially acting as a negative regulator of immune and stress-associated transcriptional programs.

### The 3PP- position of PP-InsPs is also vulnerable to enzymatic activities beyond subclade II NUDTs

Because RNA-seq data of *nudt12/13/16* lines indicated misregulation of gene expression unrelated to PSR pathways, we speculated that the accumulation of PP-InsPs other than 5-InsP_7,_ particularly 1/3-InsP_7_, might contribute to these transcriptional phenotypes. This raised the question of whether 3-InsP_7_ could also serve as a substrate for other enzymes implicated in PP-InsP metabolism. Recent work by Withfield and colleagues (Whitfield et al. 2024) demonstrated that both potato and Arabidopsis ITPK1 can phosphorylate 1-InsP_7_ and 3-InsP_7_ to 1,5-InsP_8_ and 3,5-InsP_8_, respectively. To further assess this intriguing activity, we tested all six synthetic InsP_7_ isomers using recombinant Arabidopsis ITPK1. In agreement with those findings, we observed robust phosphorylation of 3-InsP_7_, consistent with the formation of 3,5-InsP_8_. In contrast, only minor activity was detected against 1-InsP_7_, and no phosphorylation was observed for the remaining InsP_7_ isomers, indicating a strong substrate preference of ITPK1 (Figure S16A). Interestingly, human ITPK1, for which we had previously identified InsP_6_ kinase activity (Laha et al. 2019), also phosphorylated 3-InsP_7_ under similar conditions (Figure S16B), suggesting that this enzymatic activity is evolutionarily conserved. We also investigated whether ITPK1 can catalyze a reverse phosphotransferase reaction—specifically, converting 3,5-InsP_8_ and ADP into 3-InsP_7_ and ATP—as previously described (Whitfield et al. 2024). Indeed, human ITPK1 exhibited robust phosphotransferase activity in this context, as evidenced by the formation of InsP_7_ (presumably 3-InsP_7_) (Figure S16B). While Arabidopsis ITPK1 displayed negligible phosphotransferase activity under our assay conditions, the kinase domain of Arabidopsis VIH2 was capable of catalyzing a comparable reaction with 3,5-InsP_8_ and ADP, consistent with its known preference for generating or hydrolyzing 5-InsP_7_ (Figure S16B). In summary, our findings, along with those of Whitfield and colleagues (Whitfield et al. 2024) highlight that the 3-β-phosphate of PP-InsPs represents a preferred site of enzymatic activity *in vitro* across multiple enzymes from plants and humans. Those results further underscore the potential importance of distinguishing 3-InsP_7_ from its enantiomer 1-InsP_7_ *in planta*. Such a distinction will be essential to fully understand the physiological functions of these isomers.

### NUDT effectors of pathogenic ascomycete fungi display a substrate specificity reminiscent of subclade I NUDTs

Because the RNA-seq data of *nudt12/13/16* lines revealed a substantial enrichment of defense-related gene expression changes (Figure 8E and F), we speculated that plant pathogens might manipulate PP-InsP homeostasis as a virulence strategy. Supporting this idea, recent work identified NUDT hydrolase-type effectors in several plant-pathogenic fungi, which were shown to contribute to virulence (McCombe et al. 2025). The authors proposed that these effectors promote local PSRs at infection sites through their presumed 5-InsP_7_ pyrophosphatase activity.

To further assess the biochemical properties of these fungal NUDT effectors, we expressed and purified His_6_-tagged versions of candidate proteins from *Magnaporthe oryzae* (the causal agent of rice blast), *Colletotrichum graminicola* (causing anthracnose leaf blight in maize and wheat), *Colletotrichum higginsianum* (a fungus that infects many species of the Brassicaceae, including *A. thaliana*) and *Colletotrichum tofieldiae* (a root endophyte infecting *A. thaliana*). In agreement with McCombe et al., all tested fungal NUDTs displayed robust *in vitro* hydrolytic activity against 5-InsP_7_ (Figure 9A), consistent with their proposed role in modulating host P_i_ signaling. Strikingly, the substrate specificities of these effectors resembled those of Arabidopsis subclade I NUDTs. While 2-InsP_7_ and 3-InsP_7_ were largely resistant to hydrolysis, all four fungal NUDTs efficiently hydrolyzed 4-InsP_7_ and 6-InsP_7_ and showed moderate activity toward 1-InsP_7_ (Figure 9A and B, Figure S17). To better understand the functional relevance of this activity, particularly in the context of pathogenicity, we generated a catalytically inactive mutant of the *M. oryzae* NUDT, analogous to the variant described by McCombe et al. (2025), who reported reduced virulence associated with this mutation. Under our assay conditions, this mutant failed to hydrolyze not only 5-InsP_7_ but also 1-InsP_7_, 4-InsP_7_, and 6-InsP_7_ (Figure 9B).

**Figure 9:**
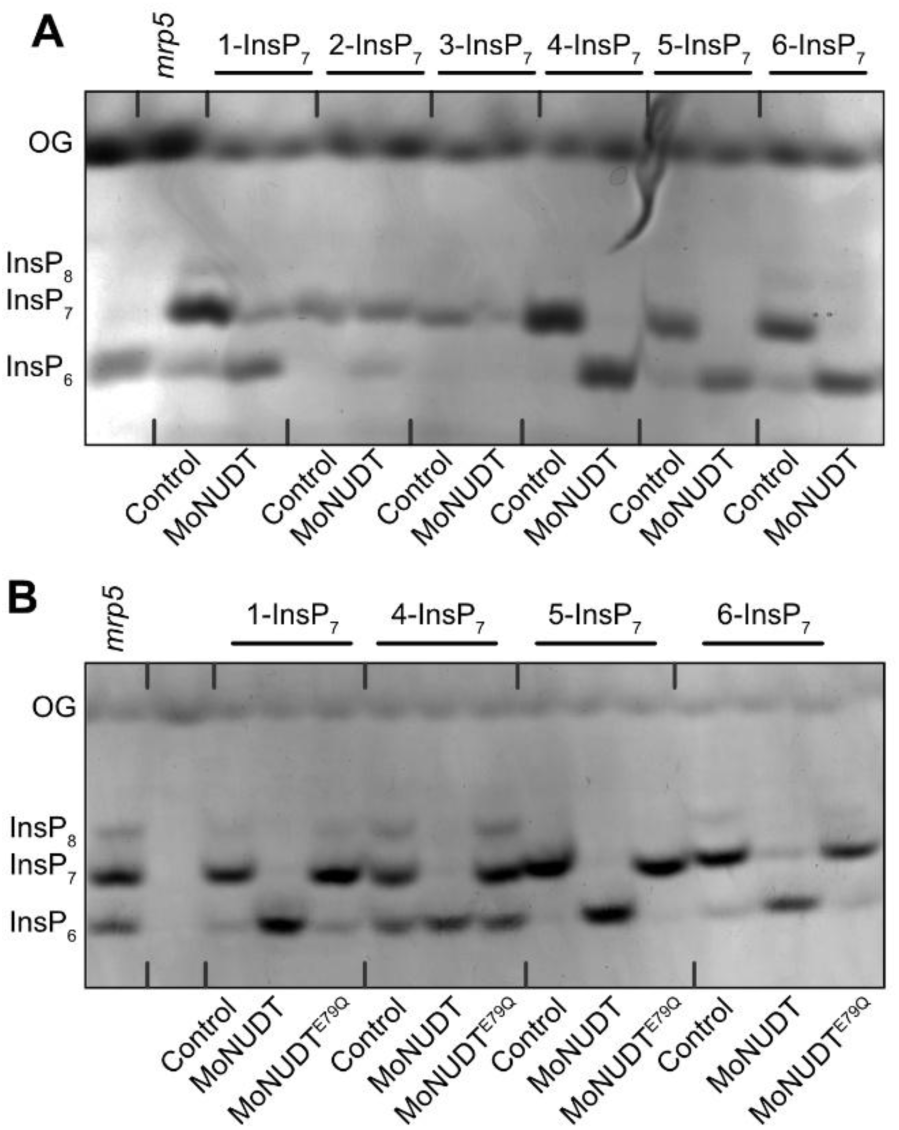
*Magnaporthe oryzae* NUDT displays substrate specificity similar to clade I NUDTs. Recombinant His6- tagged MoNUDT (**A**) and its catalytically inactive mutant (E79Q) (**B**) were incubated with 0.25 mM of the indicated InsP7 isomers. Control lanes (–) represent reactions without protein. After 45 min, reaction products were separated by 33 % PAGE and visualized using toluidine blue staining. A TiO2-purified Arabidopsis *mrp5* seed extract was included as a reference for InsP6, InsP7, and InsP8 standards. OG: Orange G loading dye.

### Subclade II NUDT hydrolases do not affect nematode parasitism but suppress antibacterial immunity in Arabidopsis

To explore potential roles of subclade II NUDTs in plant-pathogen interactions, we first examined transcript abundance of *NUDT12*, *NUDT13*, and *NUDT16* during Arabidopsis infection by cyst nematodes (CN) using previously published transcriptomic data (Siddique et al. 2022).

This analysis revealed that *NUDT16* transcript levels increased significantly during the migratory stage of infection (10 hours post infection, hpi), but decreased during the sedentary phase (12 days postinfection, dpi) (Figure S18A). In contrast, *NUDT13* expression remained unchanged during the migratory stage, but was significantly downregulated during the sedentary stage. *NUDT12* showed no significant expression changes at either stage of infection. These differential expression patterns suggest a possible role of subclade II NUDTs in response to CN infection. To assess whether these expression changes reflect a functional contribution to CN parasitism, we evaluated nematode susceptibility parameters in the *nudt12/13/16* triple mutant. No significant differences were observed in the average number of females, males, or total nematodes at 14 dpi compared to WT controls (Figure 10A). Moreover, high-resolution imaging revealed no alterations in female nematode size or the dimensions of associated syncytia (Figure 10B). These data suggest that NUDT12, NUDT13, and NUDT16 are unlikely to play a direct role in CN parasitism.

**Figure 10:**
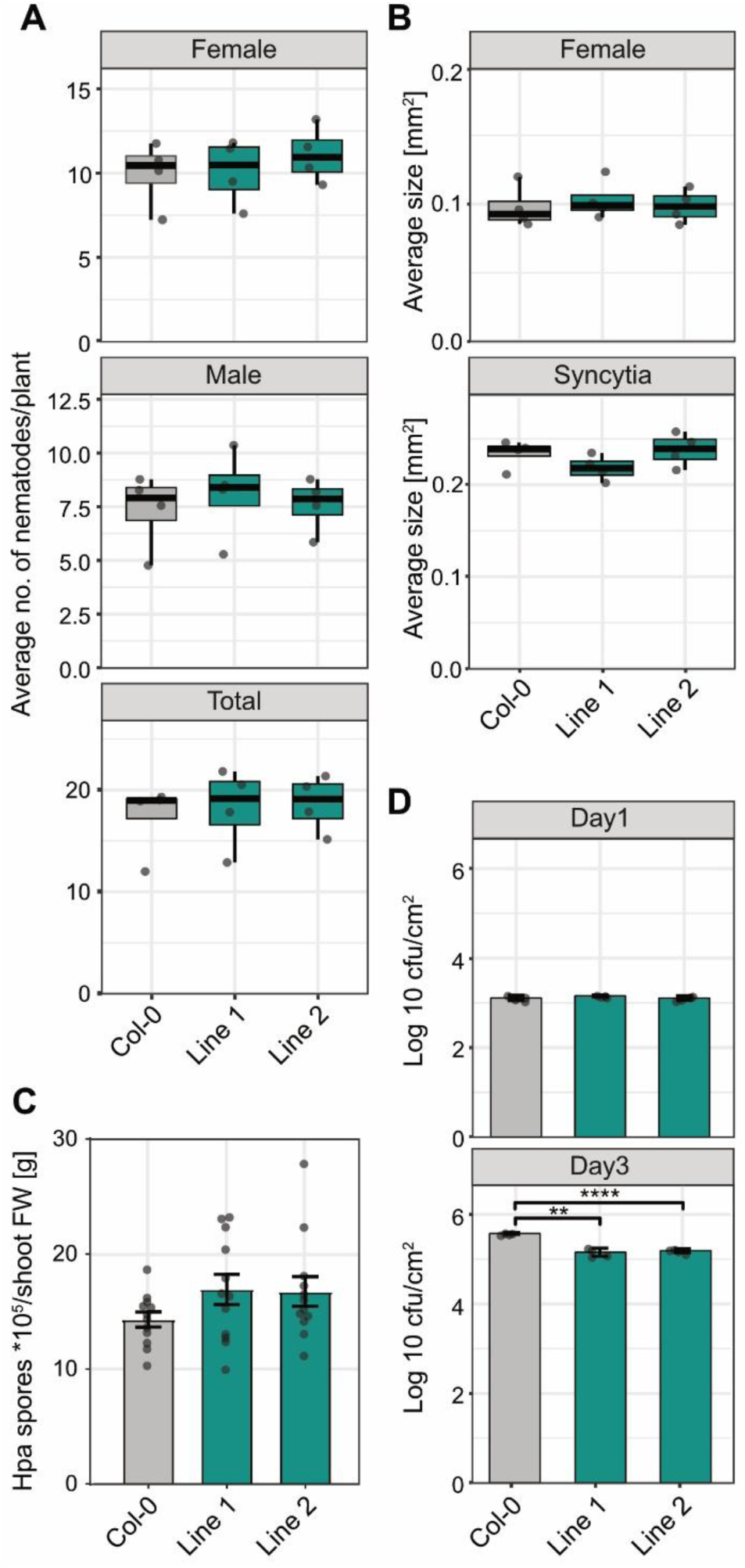
Subclade II NUDTs do not regulate responses to parasitic nematode or biotrophic oomycete infection, but is involved in plant defenses against pathogenic bacteria. **(A)** Average numbers of female, male, and total nematodes per root system at 14 days post-inoculation (dpi) in wild-type (WT) and *nudt12/13/16* triple mutant lines 1 and 2. **(B)** Average size of female nematodes and their associated syncytia at 14 dpi. Data were analyzed using the Wilcoxon test for unpaired samples (*ns*, *p* > 0.05; \**p* ≤ 0.05; \*\**p* ≤ 0.01; *\*\*\*p ≤ 0.001*). Dots represent the mean of four independent biological replicates. (**C**) Bar graphs represent the mean number of spores ± SEM per gram shoot fresh weight (FW) isolated from Arabidopsis WT (Col-0) or the indicated mutants 7 dpi with *Hyaloperonospora parabidopsidis* isolate Noco2. n = 12. For statistical analysis, an ordinary one-way ANOVA with Dunnett’s multiple comparisons test was performed. The experiment was performed three times independently with similar results. **(D)** Bar graph showing bacterial infection levels in leaves at 0 and 3 days after inoculation with *P. syringae* in WT and *nudt12/13/16* triple mutant lines 1 and 2. Data were analyzed using a two-tailed t-test (*ns*, *p* > 0.05; \**p* ≤ 0.05; \*\**p* ≤ 0.01; *\*\*\*p ≤ 0.001*). Dots represent the mean of four independent biological replicates.

To assess whether the *nudt12/13/16* triple mutant lines exhibit altered susceptibility to *Hyaloperonospora arabidopsidis* (*Hpa*), an obligate biotrophic oomycete and causing agent of downy mildew disease in Arabidopsis, we performed a spore quantification assay. Under the tested conditions, the mutants displayed a consistent trend toward slightly elevated spore counts compared to the WT (Figure 10C). This trend was observed across all three independent experiments. However, the differences were not statistically significant (*p* > 0.05 in all cases). When compared to both, the known hyper-susceptible mutant *eds1* (Parker et al. 1996) and the hypo-susceptible mutant *crp5* (Bowling et al. 1997), the tested lines showed no significant deviation from WT levels of *Hpa* proliferation. These results suggest that, under the conditions used in this study, the mutants do not exhibit altered susceptibility to *Hpa* infection.

We next evaluated the potential role of these NUDTs in plant immunity against bacterial pathogens by examining the proliferation of *Pseudomonas syringae* pv. *tomato* DC3000 in leaves of *nudt12/13/16* plants. Bacterial titers were comparable between mutant and WT plants at 1 dpi, indicating similar initial colonization (Figure 10D, upper panel). However, by 3 dpi, bacterial growth was significantly reduced in both *nudt12/13/16* lines relative to WT (Figure 10D, lower panel). The strongest effect was observed in line 1, which showed a pronounced reduction in bacterial titer (*p* < 0.0001), while line 2 also displayed a moderate but significant decrease (*p* < 0.01). Thus, these results indicate that loss of subclade II NUDTs enhances resistance to bacterial pathogens.

## Discussion

### NUDT pyrophosphatases are novel players involved in plant PP-InsP turnover

PP-InsPs were first identified more than 30 years ago in mammalian cells, when treatment of AR4-2J rat pancreatic tumor cells with high doses of sodium fluoride (NaF) led to the accumulation of these molecules (Menniti et al. 1993). NaF was presumed to inhibit endogenous pyrophosphatases, thereby revealing the rapid turnover of PP-InsPs. The authors estimated that approximately 50% of the intracellular InsP_6_ pool (50–100 µM) could be converted into low-abundance PP-InsPs (0.1–2 µM) within an hour. More recent ^18^O-water labeling experiments in yeast confirmed similarly rapid turnover, showing that 5-InsP_7_ and 1,5- InsP_8_ reach labeling equilibrium within 15–20 minutes (Kim et al. 2024) further underscoring the dynamic nature of these signaling molecules in eukaryotes.

Despite growing evidence for critical roles of PP-InsPs in plant signaling, particularly in defense, auxin perception, and PSRs, their metabolic turnover in plants has remained poorly understood. PFA-DSP-type phosphohydrolases have been shown to cleave the 5-β- phosphate of 5-InsP_7_ and 1,5-InsP_8_ *in vitro* with high specificity (Wang et al. 2022; Gaugler et al. 2022) (Figure 11), but their *in vivo* roles in PP-InsP catabolism are still unresolved. Moreover, enzymes responsible for the hydrolysis of other isomers, such as 3-, 4-, or 6-InsP_7_, had not been identified until now.

**Figure 11:**
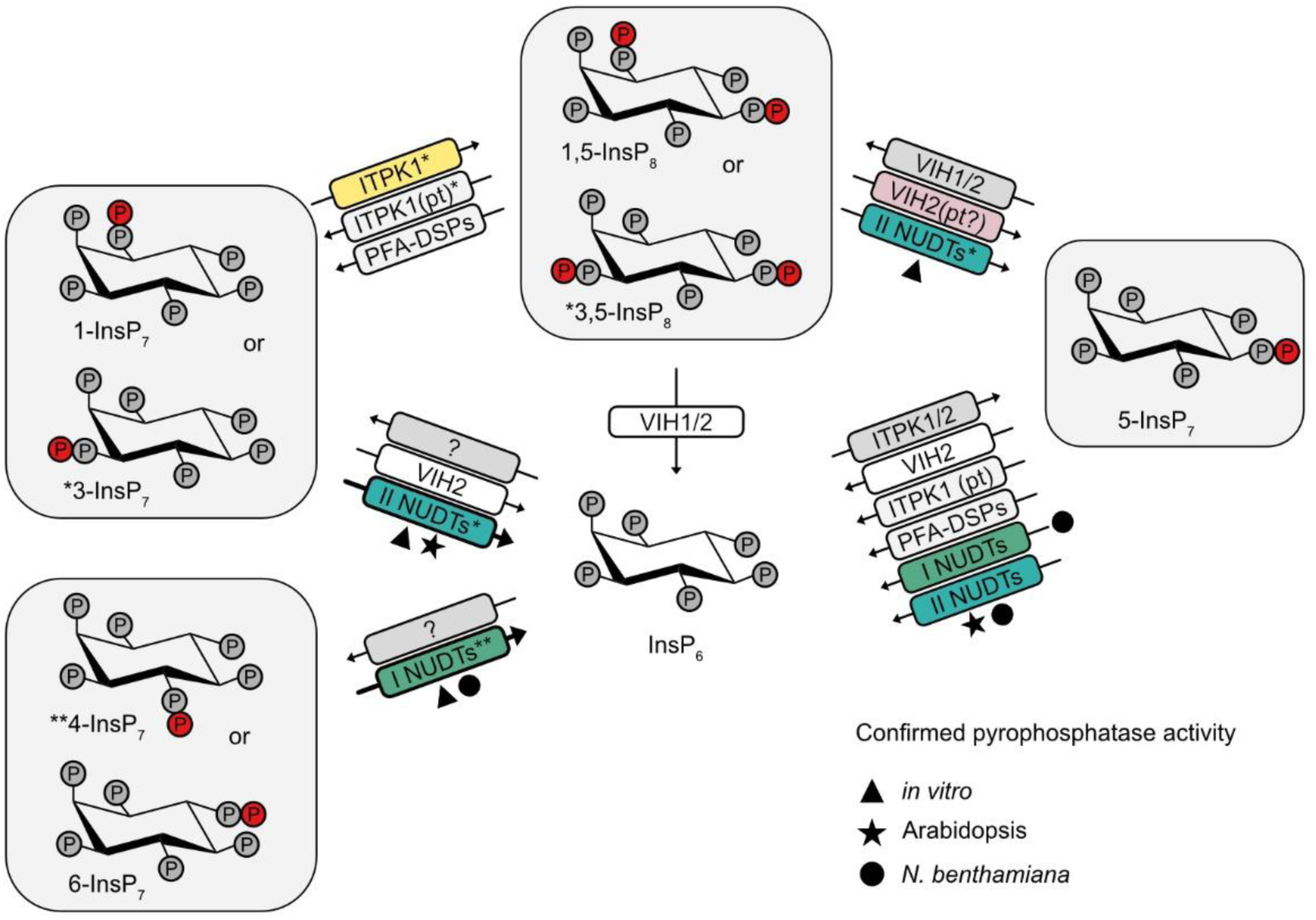
Inositol pyrophosphate metabolism in Arabidopsis. Enzymes whose catalytic activities have been confirmed at least under *in vitro* conditions are shown. Proteins highlighted in color indicate that their respective activities were identified in this work. Subclade I and II NUDTs (depicted as I NUDT and II NUDT, respectively) displayed specific pyrophosphatase activities either *in vitro*, via enzymatic assays using recombinant protein (black triangle), or *in planta*, through analyses of Arabidopsis loss-of-function mutants (black star) or via transient expression of NUDTs in *Nicotiana benthamiana* leaves (black circle). Asterisks indicate that the identity of the preferred enantiomers of distinct InsP7 species and/or of InsP8 targeted by NUDTs and ITPK1 are known. Enzymes labeled with (*) and (**) target the 3 and 4 positions, respectively. pt: phosphotransfer.

To uncover candidate PP-InsP-interacting proteins, we performed an affinity pull-down assay using 5PCP-InsP_5_ beads, a stable, non-hydrolysable analog of InsP_7_. This approach recovered known PP-InsP-related proteins including VIH1 and VIH2 (Laha et al. 2015; Zhu et al. 2019), SPX2 (Ried et al. 2021), and COI1 (Laha et al. 2015, 2016). We also isolated IPK2ɑ and IPK2β, which have been shown to be involved in the synthesis of lower InsP species (Stevenson-Paulik et al. 2002, 2005). Even though in the latter work a link of those enzymes with PP-InsPs was not determined, recent findings suggest that IPK2α may catalyze the formation of 4/6-InsP_7_ from InsP_6_ *in vitro*, implicating this protein in PP-InsP synthesis (Yadav et al. 2023). Importantly, our pull-down also identified five Arabidopsis NUDIX hydrolases, NUDT4, -16, - 17, -18, and -21, as potential PP-InsP interactors. These enzymes were recently proposed to function as inositol pyrophosphate phosphatases (Chalak et al. 2024; Laurent et al. 2024; Freed et al. 2025), and our study provides independent and complementary evidence for their specific involvement in PP-InsP turnover *in planta*. More specifically, we found that these NUDTs exhibit distinct and highly specific substrate preferences: subclade I members preferentially hydrolyze 4-InsP_7_, while subclade II enzymes show robust activity toward 3-InsP_7_ (Figures 2, Figure 3). Given that subclade II NUDTs display only minor activity toward 1-InsP_7_, and that higher-order mutants of this clade accumulate 1/3-InsP_7_ to a high degree, our findings suggest that Arabidopsis may produce 3-InsP_7_ in addition to its canonical enantiomer, 1-InsP_7_, previously identified in yeast and mammals (Mulugu et al. 2007; Fridy et al. 2007; Dollins et al. 2020).

Supporting this line of investigation, Whitfield et al.(2024), as well as data presented in this study, show that 3-InsP_7_ is a preferred substrate for the InsP_6_ kinase ITPK1, a specificity that is conserved from plants to humans (Figure S16). Although current evidence for the presence of 3-InsP_7_ is indirect, the clear enantiomer selectivity of subclade I and II NUDTs offers a promising tool for resolving the true identities of plant-derived 1/3- and 4/6-InsP_7_ isomers. This could be achieved by combining *in vitro* digests of plant-purified PP-InsPs with synthetic [^13^C_6_]-labeled InsP_7_ standards. The fact that 4/6-InsP_7_ has recently been detected in mammalian tissues (Qiu et al. 2023), and the unresolved presence of 1/3-InsP_7_ isomers in animals further underscore the broader utility of such tools.

Finally, the human DIPP1 (also known as NUDT3), which is the NUDIX hydrolase most closely related to yeast Ddp1 and to both Arabidopsis NUDT subclades, shows robust *in vitro* activity toward both 1-InsP_7_ and 3-InsP_7_ (Dollins et al. 2020), suggesting that enzymatic discrimination between these enantiomers may be evolutionarily conserved.

Our *in vitro* and *in vivo* data supporting the specific catalytic activities of Arabidopsis NUDTs are partially corroborated by previous studies, but they also highlight some apparent discrepancies that warrant further discussion. While Laurent et al. (2024) reported elevated 1/3-InsP_7_ in *nudt17/18/21* mutants, they also observed unchanged 1/3-InsP_7_ and reduced 5-InsP_7_ levels upon NUDT17 overexpression, with similar reductions in the corresponding mutants, highlighting the need for cautious interpretation. Our results do not support a role for NUDT4/17/18/21 in 1/3-InsP_7_ hydrolysis *in planta*. Besides, the absence of a metabolic phenotype in *nudt4/17/18/21* mutants may reflect redundancy with other NUDT family members or unrelated enzymes. Further studies employing combinatorial genetics and refined analytical tools will be required to determine the physiological relevance of 4-InsP_7_ pyrophosphatase (or other hydrolytic) activities linked to subclade I NUDTs in plants.

### Target specificities of subclade II NUDTs may adapt to environmental changes

We were initially intrigued by the modest, yet significant increase in 5-InsP_7_ levels observed in independent triple mutants lacking subclade II NUDTs (Figure 5D), particularly given their relatively poor activity toward this isomer under standard *in vitro* conditions (Figure 3A and B). However, transient expression of either *NUDT12* or the splice variant *NUDT16.1* in *N. benthamiana* led to a measurable reduction in 5-InsP_7_ levels (Figure 6B), suggesting that these enzymes can hydrolyze 5-InsP_7_ under certain conditions.

This observation is consistent with our *in vitro* data showing that all subclade II NUDTs exhibit some 5-InsP_7_ pyrophosphatase activity when present at higher protein concentrations (Figure S5), a feature absent in the more distantly related NUDT7. We propose that this represents a “side activity” of physiological relevance that may become functionally important under specific environmental or metabolic conditions.

Supporting this idea, both independent *nudt12/13/16* mutant lines consistently exhibited decreased shoot P levels (Figure 7A), a phenotype aligning with the established role of 5-InsP_7_ as a negative regulator of PSRs in yeast and plants (Wild et al. 2016; Riemer et al. 2021). Notably, a subtle but reproducible increase in 5-InsP_7_ was detected in both triple mutant lines, in addition to the robust accumulation of 1/3-InsP_7_ (Figure 5D). This suggests that 5-InsP_7_ turnover by subclade II NUDTs, while likely a secondary activity, may still contribute to PP-InsP homeostasis and associated nutrient signaling *in planta*. It is worth noting, however, that the 5-InsP_7_ pyrophosphatase activity of NUDT17 is approximately more than 30-fold lower than that of the *bona fide* 5-InsP_7_ phosphohydrolase PFA-DSP1 (Laurent et al. 2024). This large difference in catalytic efficiency calls into question whether NUDT17 plays a major role in 5-InsP_7_ turnover in tissues where PFA-DSPs are expressed. Laurent et al. (Laurent et al. 2024) also analyzed an NUDT17 overexpression line and reported altered levels of 5-InsP_7_ and 1,5-InsP_8_, as well as variable effects on P_i_ accumulation under high phosphate conditions. Their findings suggest a potential link between NUDT17 activity and P_i_ signaling, although some results, including similar 5-InsP_7_ reductions in both knockout and overexpression lines, highlight the need for further clarification.

### Subclade II NUDTs contribute to PP-InsP-dependent regulation of plant-pathogen interactions but not of parasitic nematodes

RNA-seq experiment with two independent *nudt12/13/16* triple mutants revealed that that the majority of DEGs were downregulated, consistent with the concept that PP-InsPs, such as 5-InsP_7_ and InsP_8_, act as negative regulators of gene expression (Dong et al. 2019; Zhu et al. 2019; Ried et al. 2021; Riemer et al. 2021). Among the DEGs were a number of PSR-related genes (Table S2), potentially explaining the reduced shoot phosphorus content observed in the mutant lines (Figure 7A).

However, several of the most strongly downregulated or upregulated genes (log2FC < -3 or > +3, Figure 8C – E, Table S3) suggest broader, PSR-independent roles for PP-InsPs in plant signaling. Notably, a subset of these DEGs included genes involved in ethylene signaling and senescence (e.g., *ATS40-2*, *ERF022*, *ERF11*, *ERF6*), while others were linked to plant immune response (e.g., *WRKY40*, *PAO2*, *IDL7*, *DHYPRP1*) (Table S3), supporting the enriched GO categories associated with defense-related pathways (Figure 8F, Figure S15B-D).

To explore whether these transcriptomic changes are functionally linked to pathogen susceptibility, we assessed the performance of *nudt12/13/16* mutants in biotic stress assays. Despite dynamic changes in *NUDT16* and *NUDT13* transcript levels during early and late stages of cyst nematode (CN) infection (Figure S18A), *nudt12/13/16* mutants showed no differences in nematode infection parameters or syncytium development compared to WT controls (Figure 10A and B). These findings suggest that subclade II NUDTs are unlikely to contribute directly to Arabidopsis responses against CN parasitism.

In addition, the sporulation of the obligate biotrophic oomycete *H. arabidopsidis* was similar in the *nudt12/13/16* mutants and the WT (Figure 10C, Figure S18B), however, a slight but non-significant increase in spore number per gram of shoot fresh weight was observed in both tested *nudt12/13/16* mutant lines in all independent experiments (Figure 10C, Figure S18B).

By contrast, we observed significantly enhanced resistance to *Pseudomonas syringae* pv. t*omato* DC3000 in both *nudt12/13/16* lines (Figure 10D) suggesting that subclade II NUDTs negatively regulate antibacterial immunity in Arabidopsis, possibly through their modulation of PP-InsP levels, particularly 1/3-InsP_7_ and maybe also 5-InsP_7_.

Taken together, our RNA-seq and pathogen assays strongly support a role for subclade II NUDTs, and by extension their PP-InsP substrates, as negative regulators of immune gene expression and bacterial defense. Although the molecular mechanisms remain to be elucidated, these findings provide functional evidence linking PP-InsP turnover to immunity, beyond the well-established PSR context.

Particularly compelling is the recent discovery that certain fungal effector proteins exhibit 5-InsP_7_ pyrophosphatase activity (McCombe et al. 2025). In our *in vitro* assays, NUDT-type effectors from diverse ascomycete pathogens hydrolyzed 4-InsP_7_ and 6-InsP_7_ robustly, and to a lesser extent 1-InsP_7_, while showing no detectable activity against 3-InsP_7_ (Figure 9, Figure S17). These substrate specificities mirror those of Arabidopsis subclade I NUDTs and suggest that fungal pathogens may exploit PP-InsP turnover to manipulate host signaling. As proposed by McCombe and colleagues, such hydrolytic activities may locally deplete 5-InsP_7_, thereby activating PSR and weakening immune responses. However, under our assay conditions, the fungal effector mutant studied by McCombe et al. lacked activity not only against 5-InsP_7_, but also against 1-, 4-, and 6-InsP_7_. This suggests that its impaired virulence may result not solely from an inability to hydrolyze 5-InsP_7_ and trigger PSR, but potentially from broader deficits in PP-InsP hydrolysis that affect additional immunity-linked signaling pathways. Thus, alterations in 4/6-InsP_7_, or a combination of multiple PP-InsP species, may contribute to defense modulation and pathogen success.

### PP-InsPs represent potential regulators of iron homeostasis

In addition to PSR- and immunity-related genes, several Fe-regulated genes were differentially expressed in *nudt12/13/16* mutants (Table S4). Notably, *IRONMAN 3* and *IRP6*, key regulators of iron deficiency responses, were strongly upregulated. This may reflect changes in cytosolic iron availability, potentially due to chelation by elevated levels of 1/3-InsP_7_ or 5-InsP_7_. Supporting this idea, genes involved in metal detoxification, such as *ZAT6* and *ZAT10* (phytochelatin regulators), and *FER1*, encoding the Fe storage protein ferritin (Briat et al. 2010), were downregulated. Consistent with impaired Fe deficiency signaling, we also observed decreased shoot Fe content in mutant lines (Figure 7B). Whether these changes result from altered Fe binding or reflect downstream effects of PP-InsP signaling remains to be clarified.

## Material and methods

### Plant materials and growth conditions

Seeds of *Arabidopsis thaliana* WT (Col-0) and T-DNA insertion lines *nudt4-1* (SALK_102051.27.90.x), *nudt4-2* (SAIL_1211_A06), *nudt17-1* (SAIL_500_D10), *nudt18-1* (SALKseq_117563.1), *nudt21-1* (SALK_031788.56.00.x), *nudt21-2* (SALKseq_132932.101) and *mrp*5 (GK-068B10) were obtained from The European Arabidopsis Stock Centre (http://arabidopsis.info/). To identify homozygous mutant lines, the plants were genotyped by PCR using the primers indicated in Table S5.

For sterile growth, *Arabidopsis thaliana* seeds were surface sterilized by washing them three times with 70 % (v/v) ethanol mixed with 0.05 % (v/v) Triton X-100 and once with 100 % (v/v) ethanol. Sterilized seeds were sown onto ½ MS plates containing 0.5 % (w/v) sucrose and 0.7 % (w/v) Phytagel (Sigma-Aldrich), pH 5.7. The seeds were stratified for 2 days at 4 °C and afterwards transferred to a growth chamber with a 16 h/8 h day/night period, equipped with white LEDs (160 μmol^-1^ m^-2^) (“True daylight”, Polyklima) and a temperature of 22 °C/20 °C to amplify seeds.

For the affinity pull-down with the 5-PCP-InsP_5_ resin, the plants were grown for 3 weeks in a growth incubator (Percival) with an 8 h/16 h day/night period, equipped with fluorescent light bulbs (120 μmol^-1^ m^-2^) and a temperature of 22 °C/20 °C. The roots and shoots of approximately 150 seedlings were collected and frozen in liquid nitrogen.

For the quantification of inositol pyrophosphate content by CE-ESI-MS, the control and mutant lines were grown on ½ MS plates, transferred to a growth chamber with a 16 h/8 h day/night period, light by white LEDs (160 μmol^-1^ m^-2^) (“True daylight”, Polyklima) and a temperature of 22 °C/20 °C. After 14 days, the shoots were collected and transferred to 2 mL centrifuge tubes and frozen in liquid nitrogen. The remaining plant material from the plates was transferred to pots containing peat substrate (Einheitserde Vermehrungserde) in block randomized design, and plants were grown for another week. 100 mg material was harvested and frozen in liquid nitrogen. The material was later grinded into powder and used for RNA extraction.

For nutrient content analyses, WT and triple mutant plants were grown for 4-5 weeks in pots containing peat-based substrate (Einheitserde Vermehrungserde) in block randomized design.

The P_i_-dependent regulation of inositol (pyro)phosphates was evaluated using Arabidopsis plants hydroponically grown as described previously (Riemer et al. 2021).

### Constructs

The open reading frames (ORFs) of *NUDT4* (At1g18300), *NUDT7* (At4g12720), *NUDT12* (At1g12880), *NUDT13* (At3g26690), *NUDT16.1* (At3g12600.1), *NUDT16.2* (At3g12600.2), *NUDT17* (At2g01670), *NUDT18* (At1g14860), *NUDT21* (At1g73540) were amplified from cDNA of *Arabidopsis thaliana* seedlings by PCR, fused to Gateway compatible *attB*1 and *att*B2 sites with primers listed in Table S5 and cloned in pDONR221 (Invitrogen) with BP clonase II (Invitrogen) following the manufacturer’s instructions. To express recombinant proteins fused N-terminally to a His_6_-maltose-binding protein (MBP) epitope tag, the ORFs were recombined into the bacterial expression vector pDest-566 (Addgene plasmid #11517, http://n2t.net/addgene:11517; RRID:Addgene_11517, kindly donated by Dominic Esposito) by LR clonase II (Invitrogen) following the manufacturer’s instructions. His-tagged MBP used as control for the *in vitro* assays was expressed with a modified pET28b vector encoding a His_8_- MBP protein (Laha et al. 2015). His_8_-MBP-VIH2 and His_8_-MBP-HsITPK were expressed with the previously described modified pETB28b vector (Laha et al. 2015, 2019).

The fungal *NUDT* genes *MoNUDT*, *CgNUDT*, *ChNUDT* and *CtNUDT* were ordered in a pTWIST ENTR vector (Twist Biosciences, San Francisco, CA, USA) based on the sequences from McCombe et al. (McCombe et al. 2025). The ORFs were then recombined into the bacterial expression vector pDest-527 (Addgene plasmid #11518, http://n2t.net/addgene:11518; RRID:Addgene_11518, kindly donated by Dominic Esposito) by LR clonase II (Invitrogen) following the manufacturer’s instructions. Site-directed mutagenesis to generate the catalytically inactive E79Q variant of *MoNUDT* was performed on the respective plasmids with the primers listed in Table S5.

For the *RUBY*-reporter construct, 1 kb *SlSPX2* promoter fragment upstream of the *SlSPX* start codon including the 5’UTR was fused upstream of the *RUBY* reporter. The given *SlSPX2* promoter-*RUBY* fusion was placed in a LII ɑ (4-5) plasmid backbone.

For yeast expression, NUDTs were fused to a C-terminal V5 epitope tag (primers listed in Table S5). The amplified genes harboring the *V5* sequence were cloned with BP clonase II in pDONR221 and then recombined into the destination vector pAG425GPD-*ccdB* (Alberti et al. 2007) (Addgene plasmid #14154, http://n2t.net/addgene:14154; RRID:Addgene_14154, kindly donated by Susan Lindquist) with LR clonase II following the manufacturer’s instructions.

For expression in *N. benthamiana*, the genes were recombined into the binary expression vector pGWB517 (Nakagawa et al. 2007). Constructs harboring *PFA-DSP1* and *PFA-DSP1^C150S^* were used as control (Gaugler et al. 2022).

### Generation of high order NUDT mutants in plants with CRISPR/Cas9

CRISPR-Cas9 constructs were generated as described earlier (Ursache et al. 2021), with sgRNAs listed in Table S6. For each of the *nudt4/nudt17/nudt18/nudt21* quadruple and the *nudt12/nudt13/nudt16* triple mutants, two constructs were generated: one with the *sgRNA-NUDT17.1*, *sgRNA-NUDT18.2*, *sgRNA-NUDT21.1* and *sgRNA-NUDT4.2* in the expression vector pRU295 (Addgene plasmid #167694, http://n2t.net/addgene:167694; RRID:Addgene_167694, kindly donated by Niko Geldner), and one construct with *sgRNA-NUDT18.1*, *sgRNA-NUDT17.2*, *sgRNA-NUDT4.1* and *sgRNA-NUDT21.2* in the expression vector pRU294 (Addgene plasmid #16768, http://n2t.net/addgene:167689; RRID:Addgene_167689, kindly donated by Niko Geldner). To generate the triple mutant, *sgRNA-NUDT12.1*, *sgRNA-NUDT13.2*, *sgRNA-NUDT16.1* and *sgRNA-NUDT16.2* were cloned in the expression vector pRU295 and sgRNA*-NUDT13.1* and *sgRNA-NUDT12.2* were cloned in the expression vector pRU294.

The triple *nudt12/nudt13/nudt16* and quadruple *nudt4/nudt17/nudt18/nudt21* mutants were then generated by co-transforming *Arabidopsis thaliana* WT plants with *Agrobacterium tumefaciens* strain GV3101 harboring the respective pRU294 or pRU295 constructs. By co-transforming the plants via floral dip, it was possible to insert CRISPR-Cas9 constructs targeting two specific sequences for each NUDT. Three days after the first infiltration, the plants were transformed again to increase the transformation efficiency.

The seeds were screened for a red and green fluorescence signal using a fluorescence stereomicroscope (OLYMPUS SZX16, Japan). The seeds were sterilized and germinated on ½ MS plates as described above. After two weeks, the seedlings were transferred to soil and plant gDNA was extracted from leaves with Phire Plant Direct PCR kit (Thermo Fisher) following the manufacturer’s instructions. The promoter and gene regions of the targets were amplified with the primers listed in Table S5 and analyzed by Sanger Sequencing.

### Yeast strains, transformation and growth

The BY4741 WT (MATa *his*3Δ *leu*2Δ *met*15Δ *ura*3Δ) and *ddp1Δ* (MATa *his*3Δ *leu*2Δ *met*15Δ *ura*3Δ *ddp1Δ::KanMX*) were obtained from Euroscarf. The yeast cells were transformed with the Li-acetate method and grown on 2x CSM-Leu plates for 3 days at 28 °C. Yeast transformants were grown in the respective media overnight at 28 °C in a rotating wheel before radioactive labeling with [^3^H]-*myo*-inositol.

### Yeast protein extraction and immunodetection

Multiple yeast transformants were cultured in 4 mL of YPD medium (3% glucose) or selective SD medium at 28 °C for up to 24 hours. Cultures were then reinoculated into fresh medium and grown for one additional day. Cells were harvested and resuspended in 500 μL of extraction buffer (300 mM sorbitol, 150 mM NaCl, 50 mM Na_2_HPO_4_, 1 mM EDTA, pH 7.5) supplemented with 100 mM β-mercaptoethanol and a 1:50 dilution of fungal protease inhibitor cocktail (Sigma-Aldrich). Cell lysis was performed by bead beating with 150–200 μL of 0.5 mm glass beads. The lysate was centrifuged, and the supernatant was mixed with sample buffer and boiled for 10 minutes. Protein samples were resolved by SDS-PAGE and analyzed by immunoblotting.

V5-tagged proteins were detected using a mouse anti-V5 primary antibody (Invitrogen, R960-25; 1:2000 dilution), followed by an Alexa Fluor Plus 800-conjugated goat anti-mouse antibody (Invitrogen; 1:20,000). Gal4 was used as a loading control and detected with a rabbit polyclonal anti-Gal4 antibody (Santa Cruz; 1:1000) and a StarBright Blue 700-conjugated goat anti-rabbit antibody (Bio-Rad; 1:2500). Detection was performed using the multiplex function of the ChemiDoc MP imaging system (Bio-Rad).

### Affinity pull-down with 5PCP-InsP_5_ resin

The plant material was powdered with mortar and pestle. 3 mL of lysis buffer (50 mM Tris pH 7.5, 100 mM NaCl, 5 mM β-mercaptoethanol, 1 mM MgCl_2_, 0.1 % Tween 20, 10 % glycerol and cOmplete™ ULTRA EDTA free protease inhibitor tablets (Roche, Basel, Switzerland)) was added to the ground shoots and 1 mL was added to the ground roots. The suspension was pelleted at 4 °C for 15 min at 19000 x g and the supernatant was transferred to a new tube and centrifuged again. The resulting extract was filtered through a 20 µm syringe filter to remove remaining particles.

A 200 µL suspension of 5-PCP-InsP_5_ beads or control beads was washed three times for 5 min in 50 mM Tris pH 7.5, 100 mM NaCl, 1 mM MgCl_2_, 10 % glycerol (Wu et al., 2016). After equilibration, the beads were incubated with the prepared plant lysates for 2 h at 4 °C on an overhead shaker. The beads were then washed three times for 10 min with the equilibration buffer and then incubated overnight at 4 °C with 100 µL InsP_6_-containing wash buffer (50 mM Tris, 100 mM NaCl, 1 mM MgCl_2_, 20 mM InsP_6_ (SiChem) on a rotating shaker. Subsequently, the beads were pelleted and washed again with wash buffer for 10 min. The beads were then further analyzed by LC-MS/MS.

InsP_6_-washed beads were submitted to an on-bead digestion. In brief, dry beads were re-dissolved in 50 µL digestion buffer 1 (50 mM Tris, pH 7.5, 2 M urea, 1 mM DTT), trypsin (125 ng) was added and beads were incubated for 30 min at 30 °C in a thermal shaker with 400 rpm. Next, beads were pelleted and the supernatant was transferred to a fresh tube. Digestion buffer 2 (50 mM Tris, pH 7.5, 2 M urea, 5 mM CAA) was added to the beads and mixed, beads were pelleted and the supernatant was collected and combined with the previous one. The combined supernatants were then incubated overnight at 32 °C in a thermal shaker with 400 rpm. Samples were protected from light during incubation. The digestion was stopped by adding 1 µL TFA and desalted with C18 Empore disk membranes according to the StageTip protocol. The proteins that were eluted with an excess of InsP_6_ were reduced with dithiothreitol (DTT), alkylated with chloroacetamide (CAA), and digested with trypsin. These digested samples were desalted using StageTips with C18 Empore disk membranes (3 M), dried in a vacuum evaporator, and dissolved in 2 % ACN, 0.1 % TFA (Rappsilber et al., 2003).

Dried peptides were re-dissolved in 2 % ACN, 0.1 % TFA (10 µL) for analysis. Samples were analyzed using an EASY-nLC 1000 (Thermo Fisher) coupled to a Q Exactive mass spectrometer (Thermo Fisher). Peptides were separated on 16 cm frit-less silica emitters (New Objective, 0.75 µm inner diameter) and packed in-house with reversed-phase ReproSil-Pur C18 AQ 1.9 µm resin (Dr. Maisch). Peptides were loaded on the column and eluted for 115 min using a segmented linear gradient of 5 % to 95 % solvent B (0 min: 5 % B; 0 - 5 min -> 5 % B; 5 - 65 min -> 20 % B; 65 - 90 min ->35 % B; 90 - 100 min -> 55 % B; 100 - 105 min -> 95 % B, 105-115 min -> 95 % B) (solvent A 0 % ACN, 0.1 % FA; solvent B 80 % ACN, 0.1 % FA) at a flow rate of 300 nL/min. Mass spectra were acquired in data-dependent acquisition mode with a TOP15 method. MS spectra were acquired in the Orbitrap analyzer with a mass range of 300–1750 m/z at a resolution of 70,000 FWHM and a target value of 3×106 ions. Precursors were selected with an isolation window of 2.0 m/z. HCD fragmentation was performed at a normalized collision energy of 25. MS/MS spectra were acquired with a target value of 105 ions at a resolution of 17,500 FWHM, a maximum injection time (max.) of 55 ms and a fixed first mass of m/z 100. Peptides with a charge of +1, greater than 6, or with unassigned charge state were excluded from fragmentation for MS2, dynamic exclusion for 30 s prevented repeated selection of precursors.

Raw data were processed using MaxQuant software (version 1.5.7.4, http://www.maxquant.org/) (Cox and Mann 2008) with label-free quantification (LFQ) and iBAQ enabled (Tyanova et al. 2016) and the match-between runs option disabled. MS/MS spectra were searched by the Andromeda search engine against a combined database containing the sequences from *A. thaliana* (TAIR10_pep_20101214; ftp://ftp.arabidopsis.org/home/tair/Proteins/TAIR10_protein_lists/) and sequences of 248 common contaminant proteins and decoy sequences. Trypsin specificity was required and a maximum of two missed cleavages allowed. Minimal peptide length was set to seven amino acids. Carbamidomethylation of cysteine residues was set as fixed, oxidation of methionine and protein N-terminal acetylation as variable modifications. Peptide-spectrum-matches and proteins were retained if they were below a false discovery rate of 1 %.

The MaxQuant output (msms.txt) was further processed using Skyline (https://skyline.ms) (MacLean et al. 2010). Peaks were inspected and, if necessary, re-integrated manually. To identify potential binding candidates, proteins were filtered for two or more unique peptides and idot products with > 0.95 confidence. Subsequently, peptides were assessed individually for enrichment in the 5-PCP-InsP_5_ condition versus the control condition based on peak areas. Background binders were identified as proteins with equal abundance in both the 5-PCP-InsP_5_ and control conditions.

### ICP-OES analyses

Shoots of 4-week-old plants were dried at 60 °C for 5 days, ground to powder and 0.150 g were digested with 3 mL nitric acid (HNO_3_, 65 %) in a microwave-accelerated reaction system (CEM, Matthews, NC, USA). Nutrient concentrations were measured using inductively coupled plasma optical emission spectroscopy (ICP-OES; iCAP Pro X ICP-OES Duo; Thermo Fisher Scientific, Waltham, MA, USA). To ensure precision and accuracy throughout the analyses standard materials (IPE100; Wageningen Evaluation Programs for Analytical Laboratories, WEPAL; Wageningen University, the Netherlands) were used.

### Protein preparation

Recombinant NUDT proteins fused to His_6_-MBP and free His_8_-MBP protein were expressed in *E. coli* BL21-CodonPlus(DE3)-RIL cells (Stratagene). The bacterial cultures were grown overnight and fresh 2YT medium (16 g/L tryptone, 10 g/L yeast extract, 5 g/L NaCl) with respective antibiotics was inoculated 1:500 with the overnight culture. In the case of CgNUDT and ChNUDT LB medium (10 g/L tryptone, 5 g/L yeast extract, 10 g/L NaCl) and in the case of CtNUDT TB medium (12 g/L tryptone, 24 g/L yeast extract, 0.4 % v/v glycerol, 0.017 M KH_2_PO_4,_ 0.072 K_2_HPO_4_) was used. The cultures were grown for 3 h at 37 °C while shaking (200 rpm) to reach an estimated OD_600_ of 0.6. Protein expression was induced by adding 0.1 mM (CgNUDT, ChNUDT, CtNUDT), 0.125 mM (NUDT7 and -12), 0.25 mM (NUDT4, -16, -17, -18, and -21) or 0.5 mM (NUDT13, MoNUDT, MoNUDT^E79Q^) isopropyl-d-1- thiogalactopyranoside (IPTG). The cultures were grown for 24 h at 12 °C (NUDT4, and -18) or at 16 °C (NUDT7, -13, -16, -17, and -21, CgNUDT, ChNUDT) or for 3 h at 37 °C (NUDT12, MoNUDT, CtNUDT). Bacterial cells were lysed by vortexing the cells 8 times for 1 min with lysis buffer (20 mM Na_2_HPO_4_, 300 mM NaCl, 2 mM DTT, 0.05 mM EDTA and 1 % Triton X-100, pH 7.4) and glass beads (0.1 – 0.25 mm). NUDT4, -17, -18, and -21 were purified using a FPLC (ÄKTA pure^TM^), while NUDT12, -13, -16, MoNUDT, MoNUDT^E79Q^, CgNUDT, ChNUDT and CtNUDT were batch purified using Ni-NTA agarose resin (Macherey-Nagel). For both methods, the same binding buffer (20 mM Na_2_HPO_4_, 500 mM NaCl, 25 mM imidazol, pH 7.4) and elution buffer (20 mM Na_2_HPO_4_, 500 mM NaCl, 500 mM imidazol, pH 7.4) were used. The recombinant proteins were concentrated and imidazole was diluted by using Vivaspin 20 (50 kDa, Merck) for FPLC-purified proteins or Amicon Ultra 0.5 mL Centrifugal Filters (50 kDa, Merck) for the batch-purified proteins and the elution buffer without imidazole. The purified proteins were mixed with 20 % glycerol and stored at either at -20 °C or -80 °C. Recombinant Arabidopsis and human ITPK1 and VIH2 proteins fused to His_8_-MBP were expressed and purified as described before (Laha et al. 2015, 2019). Concentrations were estimated by using SDS-PAGE followed by Coomassie blue staining and comparing the proteins with designated amounts of a BSA standard.

### *In vitro* (PP-)InsP assays

The (PP-)InsP pyrophosphatase assays were performed with recombinant His_6_-MBP-NUDT proteins or His_8_-MBP proteins as control and 50 mM HEPES (pH 7.0), 10 mM NaCl, 5 % (v/v) glycerol, 0.1 % (v/v) β-mercaptoethanol, either 1 mM EDTA, MnCl_2_, MgCl_2_, CaCl_2_, or ZnCl_2_ and 0.33 mM of various InsP_7_ or InsP_8_ isomers as indicated. The recombinant protein concentrations were: ∼4.9 µM His_6_-MBP-NUDT4, ∼3.7 µM His_6_-MBP-NUDT12, ∼1.9 µM His_6_-MBP-NUDT13, ∼0.6 µM His_6_-MBP-NUDT16.1, ∼15.8 µM His_6_-MBP-NUDT16.2, ∼2.0 µM His_6_-MBP-NUDT17, ∼5.1 µM His_6_-MBP-NUDT18 and ∼5.0 µM His_6_-MBP-NUDT21. Since our aim was to determine which InsP7 serves as the preferred or most susceptible substrate for a given NUDT and not to directly compare activities between enzymes, the recombinant protein concentrations were individually adjusted to identify substrate specificity while minimizing nonspecific activity. To obtain normalized, quantitative activity data, NMR-based enzyme assays were performed (see below). For the assays with His_6_-MBP-MoNUDT, His_6_-MBP-MoNUDT^E79Q^, His_6_-MBP-CgNUDT, His_6_-MBP-ChNUDT and His_6_-MBP-CtNUDT 0.25 mM of the various InsP_7_ isomers and 0.13 µg/µL protein were used.

PP-InsP pyrophosphatase assays with higher subclade II protein concentrations were performed with ∼7 µM His_6_-MBP-NUDT7, ∼7.5 µM His_6_-MBP-NUDT12 or His_6_-MBP-NUDT13, and ∼8 µM His_6_-MBP-NUDT16.1 or His_6_-MBP-NUDT16.2. The pyrophosphatase assays with a higher subclade I protein concentration were performed in a reaction buffer as described previously (Riemer et al. 2021) with 0.33 mM of various InsP_7_ or InsP_8_ isomers as indicated. The recombinant protein concentrations were ∼9 µM His_6_-MBP-NUDT4, ∼6 µM His_6_-MBP-NUDT17, ∼6 µM His_6_-MBP-NUDT18 and ∼8 µM His_6_-MBP-NUDT21. InsP_7_ and InsP_8_ isomers were chemically synthesized as described previously (Capolicchio et al. 2013, 2014). The reactions were incubated for 1 h at 28 °C or 2 h at 22 °C and either separated by 33 % PAGE and visualized by toluidine blue staining or analyzed by CE-ESI-MS as described previously (Qiu et al. 2020; Riemer et al. 2021). Kinase assays were performed as described previously (Riemer et al. 2021) with 0.2 µg/µL of protein.

### Nuclear magnetic resonance spectroscopy (NMR)

Phosphatase assays were performed as previously described by Laurent *et al*. (Laurent et al. 2024). Reaction mixtures contained 100 µM of the respective [¹³C_6_]-labeled PP-InsP substrate in a total volume of 600 µL, prepared using one of two buffer systems. Buffer system A consisted of 50 mM HEPES (pH 7.0), 10 mM NaCl, 0.1 % (v:v) β-mercaptoethanol, 1 mM MgCl_2_, 5 % (v:v) glycerol and 0.2 mg/mL BSA. Buffer system B contained 50 mM HEPES (pH 7.3), 150 mM NaCl, 1 mM DTT, 0.5 mM MgCl_2_ and 0.2 mg/mL BSA. Reaction mixtures were pre-incubated at 28 °C (buffer A) or 37 °C (buffer B), and reactions were initiated by the addition of the appropriate amount of enzyme. For MBP-NUDT13, 500 nM was used for 1-InsP_7_ in buffer B at 37 °C and for 5-InsP_7_ in buffer A at 28 °C; 250 nM for 5PP-InsP₅ in buffer B at 37 °C; and 1.9 µM for 1PP-InsP₅ in buffer A at 28 °C. For MBP-NUDT17, 2.5 µM was used for 5-InsP_7_ and 1-InsP_7_ in buffer A at 28 °C, as well as for 1-InsP_7_ in buffer B at 37 °C; and 2 µM for 5-InsP_7_ in buffer B at 37 °C. Reactions were continuously measured at the given temperature with a nuclear magnetic resonance (NMR) pseudo-2D spin-echo difference experiment (Harmel et al. 2019). To record NMR spectra, a Bruker AV-III spectrometer (Bruker Biospin) equipped with a cryo-QCI probe operating at 600 MHz for ^1^H and 151 MHz for ^13^C nuclei was used. TopSpin 3.5 was used for both measurements as well as NMR data analysis. An individual NMR spectrum was recorded every 86s, the number of scans per spectrum amounted to 128. Quantification of PP-InsP species was carried out by plotting the relative intensity changes of the C2 peaks of the respective PP-InsPs as a function of reaction time. Raw data was evaluated in GraphPad PRISM 5 by adding different trend lines depending on the reaction progress and the resulting progress curve. In case of a linear progress curve, linear regression was used to derive the reaction rate. For hyperbolically shaped progress curve, a nonlinear fit was used and velocity rates were calculated by using the first derivatization of the one phase decay model. Reaction rates derived from both the linear as well as non-linear regression were normalized by the phosphataseś mass concentration excluding the mass of the MBP tag.

### (PP-)InsP extraction and purification

The extraction and purification of InsPs of Arabidopsis T-DNA insertion lines were performed using TiO_2_ (titanium(IV) oxide rutile, Sigma Aldrich) beads as previously described (Riemer et al. 2021). InsPs from the triple mutant *nudt12/13/16*, quadruple mutant *nudt4/17/18/21* and *N benthamiana* were extracted with Nb_2_O_5_ beads (Niobium (V) pentoxide, Sigma Aldrich) with the following method: The beads were weighed for each sample and washed once in water (300 µL) and once in 1 M perchloric acid (300 µL) (PA). Liquid nitrogen-frozen plant material was homogenized using a pestle and immediately resuspended in 800 µL ice-cold PA (1 M). Samples were kept on ice for 10 min with short intermediate vortexing and then centrifuged for 10 min at 18000 g at 4 °C. The supernatants were transferred into fresh 1.5-mL tubes and centrifuged again for 10 min at 18000 g at 4 °C. To bind InsPs onto the beads, the supernatants were resuspended in the prewashed Nb_2_O_5_ beads and rotated at 4 °C for 45 min. The tubes were then centrifuged at 11000 g for 1 min at 4 °C and washed twice in 500 µL ice-cold PA (1 M) and once with 500 µL PA (0.1 M). To elute InsPs, beads were resuspended in 200 µL 10 % ammonium hydroxide and then rotated for 5 min at room temperature. After centrifuging, the supernatants were transferred into fresh 1.5-mL tubes. The elution process was repeated and the second supernatants were added to the first. Eluted samples were vacuum evaporated at 45 °C to dry completely.

### PAGE, CE-ESI-MS and HPLC analyses

Arabidopsis plants and yeast transformants were radioactively labeled with [^3^H]-*myo*-inositol (ARC), extracted and analyzed via SAX-HPLC as described before (Azevedo and Saiardi 2006; Laha et al. 2015; Gaugler et al. 2020). *In vitro* biochemical assays were analyzed by PAGE and CE-ESI-MS, and purified InsPs of Arabidopsis mutant lines or *N. benthamiana* leaves were analyzed by CE-ESI-MS as described in (Qiu et al. 2020, 2023; Riemer et al. 2021).

### RNA extraction and quantitative real-time PCR

Total RNA was extracted from homogenized root or shoot samples using the NucleoSpin RNA Mini Kit (Machery-Nagel), followed by on-column DNase treatment (QIAGEN), according to the manufacturers’ protocols. cDNA was synthesized from 0.5-1 µg RNA by reverse transcription using the RevertAid First Strand cDNA synthesis Kit (Thermo Fisher Scientific) and oligo(dT) primer. A 10- or 20-times diluted cDNA sample was then used for quantitative real-time (RT) PCR analysis with the CFX384 Touch Real-Time PCR Detection System (Bio-Rad Laboratories) and the iQ SYBR Green Supermix (Bio-Rad Laboratories). All reactions were repeated in two technical and three biological replicates. Recorded C_t_ values were exported from the Bio-Rad CFX Manager Software (Version 3.1, Bio-Rad Laboratories) and used for the calculation of relative expression values according to ΔΔCT method and using *UBQ2* (*At2g36170*) or *ACTIN2* (*At3g18780*) as the reference gene. The reactions were carried out using the primers listed in Table S5.

Total RNA for transcriptome analysis was extracted from shoot samples using the QIAGEN Kit followed by on-column DNase treatment (QIAGEN), according to the manufacturers’ protocols.

### RNA sequencing, bioinformatic analysis and data visualization

For RNA-seq analysis, total RNA was extracted as described above. Library construction and sequencing were performed at BGI Genomics (Poland). mRNA enrichment was performed on total RNA using oligo(dT)-attached magnetic beads. The enriched mRNA with poly(A) tails was fragmented using a fragmentation buffer, followed by reverse transcription using random N6 primers to synthesize cDNA double strands. The synthesized double-stranded DNA was then end-repaired and 5’-phosphorylated, with a protruding ’A’ at the 3’ end forming a blunt end, followed by ligation of a bubble-shaped adapter with a protruding ’T’ at the 3’ end. The ligation products were PCR amplified using specific primers. The PCR products were denatured to single strands, and then single-stranded circular DNA libraries were generated using abridged primer. The constructed libraries were quality-checked and sequenced after passing the quality control. Sequencing was performed on the DNBSEQ platform using paired-end 150 bp reads(PE150). The raw data obtained from sequencing was filtered using SOAPnuke (Chen et al. 2017) to remove adapters, reads containing ploy-N and low-quality reads, and the filtered data (clean reads) was aligned to the Arabidopsis genome (TAIR10) using HISAT2 (https://daehwankimlab.github.io/hisat2/) based on Burrows-Wheeler transform and Ferragina-Manzini (FM) index. Clean data was aligned to the reference gene set using Bowtie2 (v2.3.4.3). Gene expression quantification was performed using RSEM (v1.3.1) software, and gene expression clustering heatmaps across different samples were generated using pheatmap (v1.0.8) (10.32614/CRAN.package.pheatmap). DESeq2 (v1.4.5) (Love et al. 2014) or DEGseq (Wang et al. 2010) or PoissonDis (Audic and Claverie 1997)) was employed for differential gene (DEGs) detection, with criteria set as Q (http://bioconductor.org/packages/qvalue/) value < 0.05 or FDR ≤ 0.001 and (|log2FC| > 1). The subsequent analysis and data mining were performed on Dr. Tom Multi-omics Data mining system (https://biosys.bgi.com).

For the GO terms (http://www.geneontology.org/), the base mean expression for each gene was calculated as the average expression of mutant and control samples (base mean = (avg. Exp. of Mutant + avg. Exp of Control) / 2). Genes with a base mean greater than 20 were selected to form the background set. From this background, an interest set was defined by selecting genes with a log2 fold change (|log2FC| > 1), and Q-values < 0.05. GO enrichment analysis was performed using the topGO (https://bioconductor.org/packages/release/bioc/html/topGO.html) package for all three gene ontology categories: biological process (BP), cellular component (CC), and molecular function (MF). The weigh01 algorithm and Fisher’s exact test were employed as parameters, and the results were filtered to include the top 200 GO terms (nodes). The significance of each GO term was determined by the associated *p*-value, and the top 10 GO terms were plotted in the order of the *p*-values. Data analysis and visualization was done using R software (https://www.r-project.org/, R Core Team, 2023). GO enrichment analysis was conducted using the hypergeometric test, implemented via the Phyper function in R, with a significance threshold of Q ≤ 0.05.

### Tobacco growth conditions and infiltration procedure

*N. benthamiana* seeds were germinated in a growth chamber with a 16 h/8 h day/night period equipped with fluorescence light bulbs and a temperature of 22 °C/20 °C for 5-6 weeks prior to infiltration.

A single colony of transformed *Agrobacterium tumefaciens* strain GV3101 containing T-DNA vectors was inoculated in 2 mL of YEB media containing the respective antibiotics and cultivated overnight at 28 °C while shaking. On the following day, 500 µL of overnight culture was added to 5 mL of fresh YEB with respective antibiotics and grown for another 4-6 h at 28 °C. Cultures were harvested by centrifugation at room temperature with 4000 g for 10 min. The pellet was resuspended in 250 µL infiltration medium containing 10 mM MgCl_2_, 10 mM MES (pH 5.8) and 150 μM of freshly added acetosyringone. OD_600_ was determined using a 1:10 dilution and adjusted to OD_600_ = 0.8 in infiltration medium. The working solution for the RUBY assay was prepared by adding equal amounts of RUBY-reporter and NUDTs strains, and adding p19 (OD_600_ = 0.05). The NUDT overexpression experiment in the absence of the RUBY-reporter was done in a similar way. The mixtures were then infiltrated into the abaxial surface of 5–6-week-old *N. benthamiana* leaves using a 1 mL syringe without a needle. After infiltration, the plants were kept under previous growth conditions for 2-3 days before analyzing RUBY staining or extracting and purifying the inositol phosphates.

### Pathogen Assays

For nematode infection assays, WT Arabidopsis (Col-0), *nudt12/13/16-1*, and *nudt12/13/16-2* plants were grown on 0.2 % (w/v) Knop medium in Petri dishes, with two plants per dish. Growth conditions were maintained at ∼23 °C under a 16-hour light/8-hour dark photoperiod for 12 days prior to inoculation (Chopra et al. 2021). To obtain *Heterodera schachtii* juveniles, approximately 300 sterile cysts harvested from a monoculture on mustard (*Sinapis alba* ‘Albatros’) roots were placed in a funnel and incubated in 3 mM ZnCl_2_ in darkness for 5 days. Emerged second-stage juveniles (J2s) were collected and washed with sterile tap water before use. Nematode infection assays were performed following the protocol by Hasan et al. (Hasan et al. 2022). Each plant was inoculated under sterile conditions with 60–80 freshly hatched, active J2s. Each genotype included 20–30 plants per biological replicate, and the experiment was repeated across four independent biological replicates. At 14 days post-inoculation (dpi), the number of female and male nematodes per root system was recorded. Additionally, 40–50 female nematodes and their associated syncytia were imaged using an M165C stereomicroscope and measured with LAS v. 4.3 image analysis software (Leica Microsystems).

A separate set of experiments was conducted to assess susceptibility to the biotrophic oomycete pathogen *Hyaloperonospora arabidopsidis*. Arabidopsis seedlings (14 days old) of Col-0, *cpr5*, *eds1*, *nudt12/13/16* line 1 and *nudt12/13/16* line 2 were spray-inoculated with a suspension of *H. arabidopsidis* (*Hpa*) isolate Noco2 spores (5 × 10^4^ spores/mL) as described previously (Asai et al. 2015). After inoculation, plants were kept at 18 °C under a 10 h light/14 h dark photoperiod in trays covered with sealed lids to maintain high humidity. At 7 dpi, infected aerial parts were harvested, placed in 1 mL sterile water, and vortexed to release spores. Spores were counted using a hemocytometer and normalized to leaf fresh weight.

Bacterial growth assay was performed with the *P. syringae* strain *DC3000* and according to Gulabani et al. (Gulabani et al. 2022). Briefly, fully expanded leaves from 4-week-old plants grown under short-day conditions (16 h light/ 8 h dark photoperiod) were infiltrated with the bacterial strain at a density of 10^5^ cfu/ml, using a syringe without a needle. Bacterial accumulation was quantified by harvesting leaf discs of defined area at 0 and 3 days post-infiltration (dpi) and macerating them in 10 mM MgCl_2_. Serial dilutions of the extracts were plated on Pseudomonas F agar media plates containing 25 μg/mL Rifampicin and 100 ug/mL Cycloheximide. Bacterial growth is reported as Log_10_ cfu/cm^2^. Statistical analysis according to Student’s *t*-test is a pairwise comparison of bacterial accumulation in the mutants versus WT plants.

## Data availability

The mass spectrometry proteomics data have been deposited to the ProteomeXchange Consortium via the PRIDE (Perez-Riverol et al. 2022) partner repository with the dataset identifier PXD056943.

## Acknowledgments

The authors thank Brigitte Ueberbach, Li Schlüter, Nur Gömec, Anna M. Frentzen (Department of Plant Nutrition, Institute of Crop Science and Resource Conservation, University of Bonn), Jacqueline Fuge (Department of Physiology & Cell Biology, Leibniz-Institute of Plant Genetics and Crop Plant Research) and Anne Harzen (Plant Proteomics and Mass Spectronomy Group, Max Planck Institute for Plant Breeding Research) for excellent technical assistance.

The author(s) declare that financial support was received for the research, authorship, and/or publication of this article. This work was funded by grants from the Deutsche Forschungsgemeinschaft (SCHA 1274/4-1, SCHA 1274/5-1, and under Germany’s Excellence Strategy, EXC-2070–390732324, PhenoRob to G.S.; JE 572/4-1 and under Germany’s Excellence Strategy, CIBSS–EXC-2189–Project ID 390939984 to H.J.J.; LA 1338/18-1 to T.L.; and TRR356/I (491090170), TP-B08 and Project number 451218338 to M.K.R.-L.), by the Marie Skłodowska-Curie Action (Grant Agreement ID 101108767) to S.W., and by the Department of Biotechnology, Government of India (Grant number BT/PR45561/AGIII/103/1386/2023) to S.B.

## Author contributions

R.S., V.G., and G.S. conceived the study; R.S., V.G., Kl.L, G.S., T.L., R.F.H.G., S.B., F.M.W.G., M.K.R.-L., H.J.J., H.N., Do.F. designed experiments; R.S., Kl.L., I.P., V.G., A.S., S.C.S, S.M.B., Ke.L, E.L., J.M.S., K.R., Da.F., N.F.; M.S.H., A.M., Sa.K., S.W., Y.Z.B., Si.K., M.H., P.G., and R.F.H.G. performed experiments; R.S., Kl.L., I.P., V.G., S.C.S, A.S., J.T., H.S., L.K., P.G., M.K., H.N., and R.F.H.G., G.S. wrote the manuscript with input from all authors. All authors have read and agreed to the published version of the manuscript.

