## Supporting information for "NUDIX Hydrolases Target Specific Inositol Pyrophosphates and Regulate Phosphate Homeostasis and Bacterial Pathogen Susceptibility in Arabidopsis"

Schneider et al.

Supporting Information

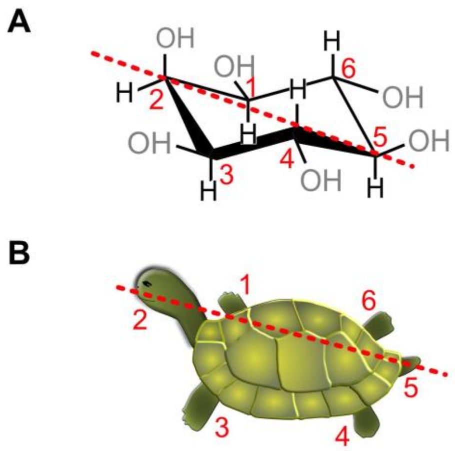

Figure S1: Structure of *myo*-inositol. (A) Spatial representation as chair conformation. (B) Schematic representation based on Agranoff´s turtle.

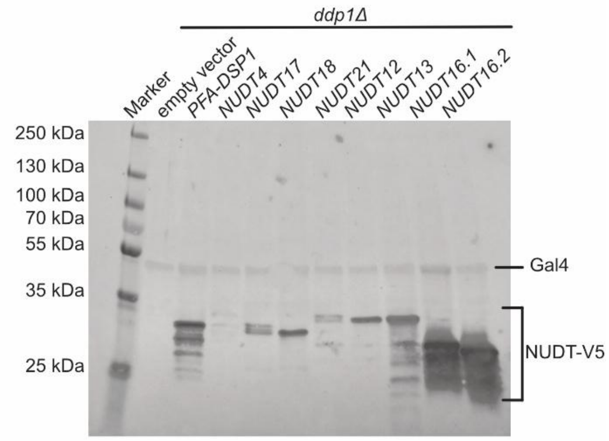

Figure S2: NUDTs from subclades I and II exhibit differential expression in the *ddp1Δ* yeast strain. Immunoblot analysis was performed on protein extracts from *ddp1Δ* yeast transformed with either an empty pAG425-GPD-*ccdB* plasmid, pAG425 carrying *PFA-DSP1* (positive control), or pAG425 constructs encoding *NUDTs* fused to a C-terminal V5 tag. Detection of V5-tagged proteins was carried out using an anti-V5 primary antibody (Invitrogen; 1:2000 dilution) and an Alexa Fluor Plus 800-conjugated anti-mouse secondary antibody (Invitrogen, goat; 1:20000 dilution). Gal4 protein levels served as a loading control and were simultaneously detected using a polyclonal anti-Gal4 antibody (Santa Cruz; 1:1000 dilution) along with a StarBright Blue 700-conjugated anti-rabbit secondary antibody (Bio-Rad, goat; 1:2500 dilution). Signals were visualized using the multiplex mode of the ChemiDoc MP imaging system (Bio-Rad).

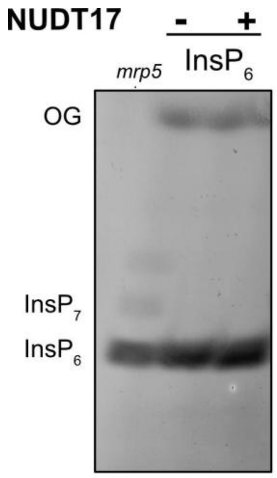

Figure S3: NUDT17 does not hydrolyze $InsP_6$ *in vitro*. Recombinant $His_6$-MBP-NUDT17 was incubated with 0.33 mM $InsP_6$ and 1 mM $MgCl_2$ at 28 °C. Recombinant $His_8$-MBP served as a negative control (indicated with the minus). After 1 h, the reactions were analyzed by 33 % PAGE and visualized by toluidine blue staining. OG: orange G.

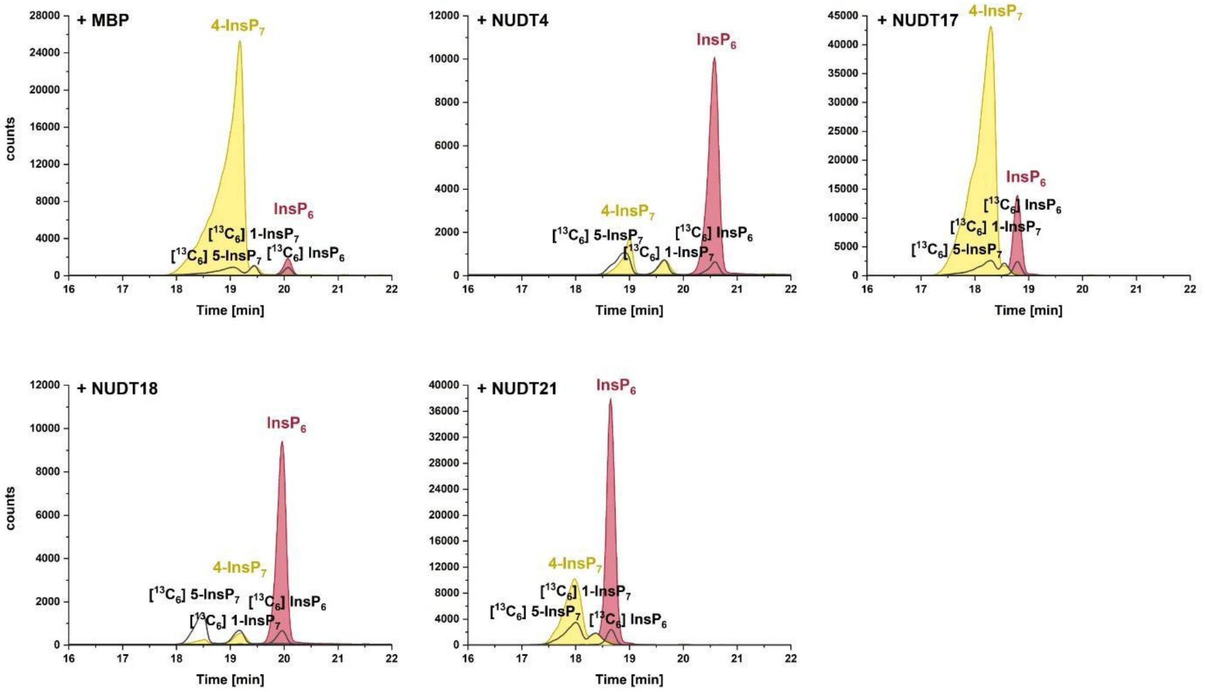

Figure S4: Arabidopsis NUDT hydrolases of subclade I display 4-$InsP_7$ pyrophosphatase activity *in vitro*. Recombinant $His_6$-MBP-NUDT proteins were incubated with 0.33 mM 4-$InsP_7$ and 1 mM $MgCl_2$ at 28 °C. $His_8$-MBP served as a negative control. $His_6$-MBP-NUDT4, -17, -18, or -21 were incubated with 4-$InsP_7$. After 1 h, the reaction products were spiked with isotopic standards mixture ([$^{13}C_6$] 1,5-$InsP_8$, [$^{13}C_6$] 5-$InsP_7$, [$^{13}C_6$] 1-$InsP_7$, [$^{13}C_6$] $InsP_6$, [$^{13}C_6$] 2-OH $InsP_5$) and subjected to CE-ESI-MS analyses.

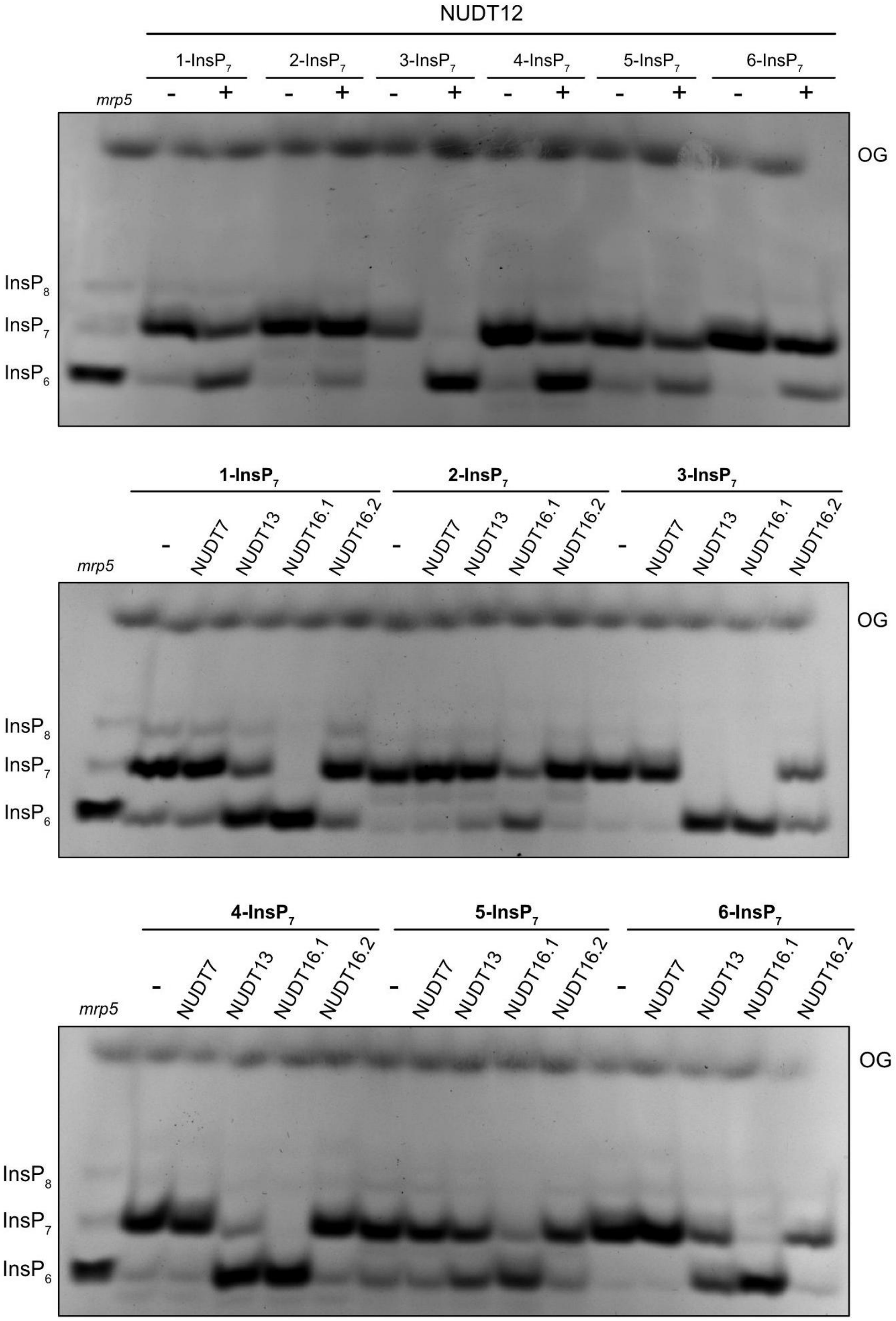

Figure S5: Subclade II NUDT hydrolases lose substrate specificity at higher concentrations *in vitro*. Recombinant $His_6$-MBP-tagged NUDT7 (~7 µM), NUDT12 (~7.5 µM), NUDT13 (~7.5 µM), NUDT16.1 (~8 µM) or NUDT16.2 (~8 µM) was incubated with 0.33 mM $InsP_7$ and 1 mM $MgCl_2$ at 28 °C (indicated with the plus symbol). $His_8$-MBP served as a negative control (indicated with the minus symbol). After 1 h, the reaction products were separated by 33 % PAGE and visualized by toluidine blue. OG: orange G.

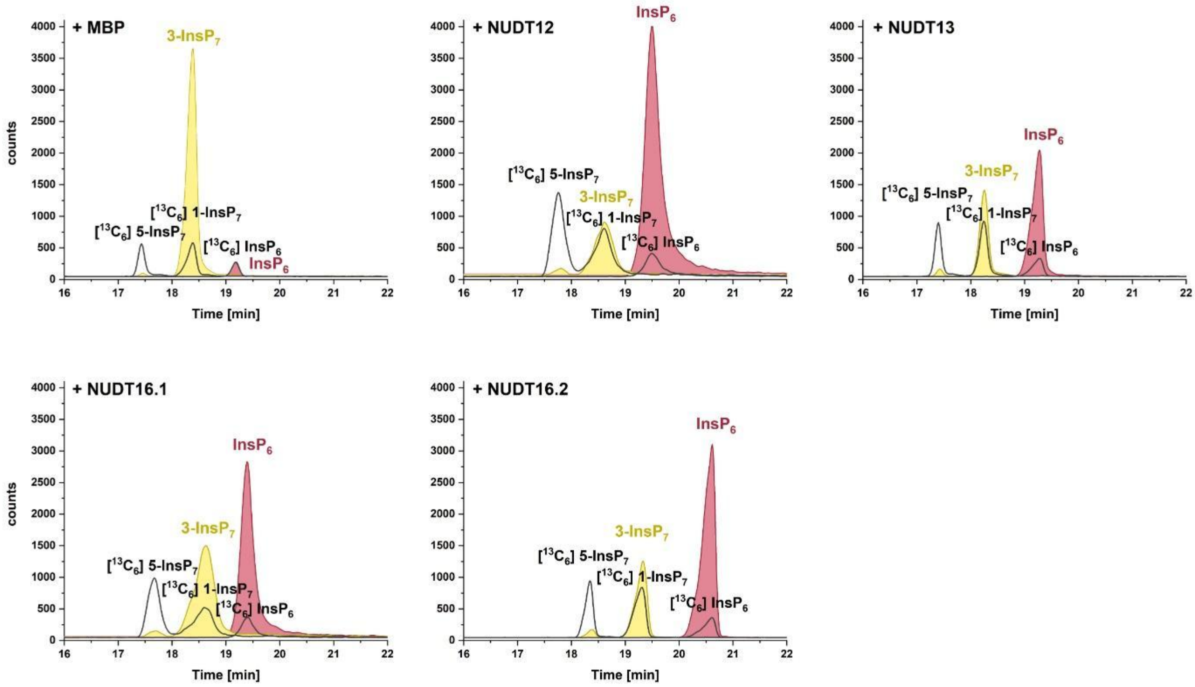

Figure S6: Arabidopsis NUDT hydrolases of subclade II display 3-InsP$_7$ pyrophosphatase activity *in vitro*. Recombinant His$_6$-MBP-NUDT proteins were incubated with 0.33 mM 3-InsP$_7$ and 1 mM MgCl$_2$ at 28 °C. His$_8$-MBP served as a negative control. His$_6$-MBP-NUDT12, -13, -16.1, or -16.2 were incubated with 3-InsP$_7$. After 1 h, the reaction products were spiked with isotopic standards mixture ([$^{13}$C$_6$] 1,5-InsP$_8$, [$^{13}$C$_6$] 5-InsP$_7$, [$^{13}$C$_6$] 1-InsP$_7$, [$^{13}$C$_6$] InsP$_6$, [$^{13}$C$_6$] 2-OH InsP$_5$) and subjected to CE-ESI-MS analyses.

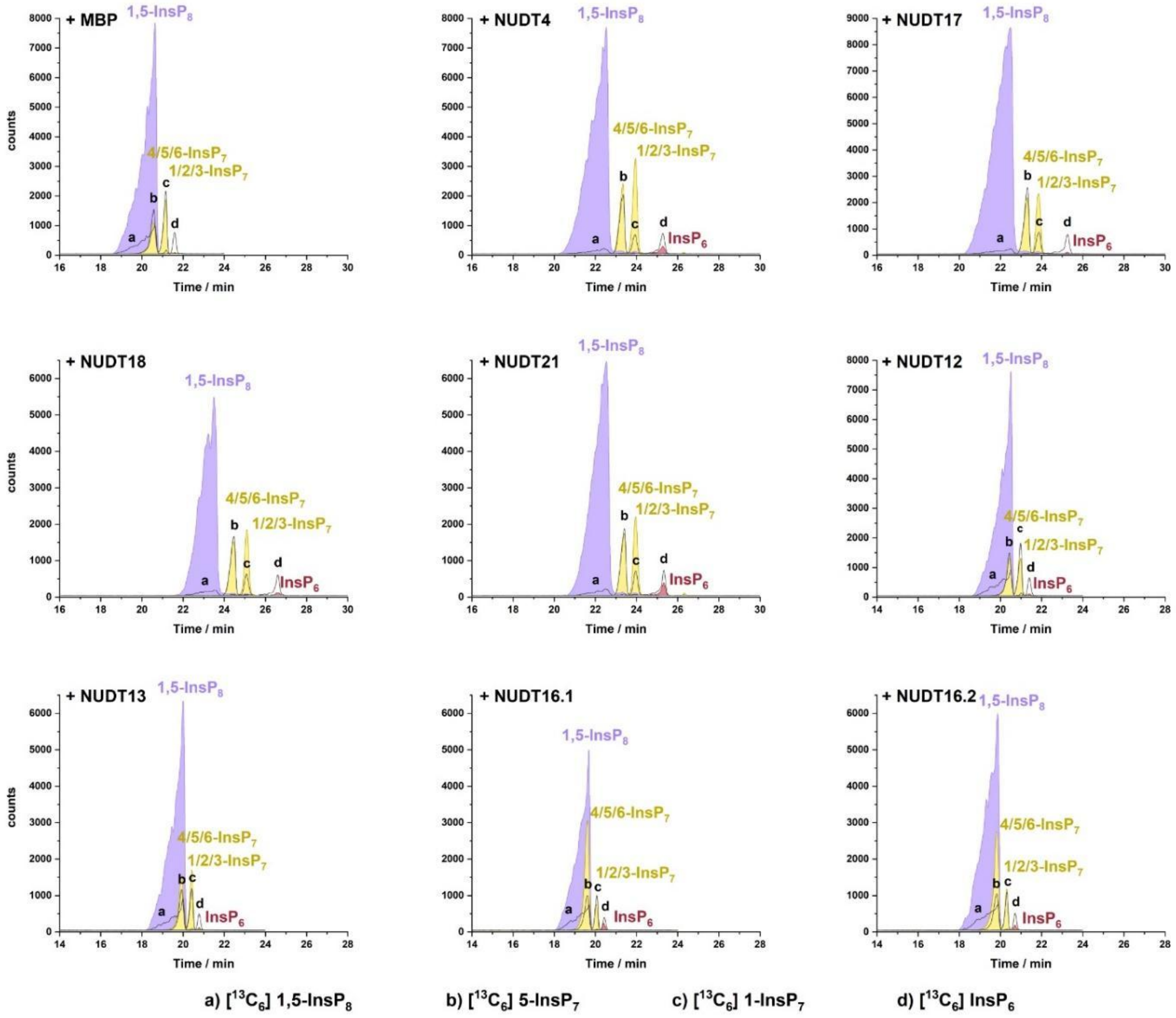

Figure S7: Arabidopsis NUDT hydrolases show a weak hydrolysis activity toward 1,5-$InsP_8$ *in vitro*. Recombinant $His_6$-MBP-NUDT proteins were incubated with 0.33 mM 1,5-$InsP_8$ and 1 mM $MgCl_2$ at 28 °C. $His_8$-MBP served as a negative control. After 1 h, the reaction products were spiked with isotopic standards mixture ([$^{13}C_6$] 1,5-$InsP_8$, [$^{13}C_6$] 5-$InsP_7$, [$^{13}C_6$] 1-$InsP_7$, [$^{13}C_6$] $InsP_6$, [$^{13}C_6$] 2-OH $InsP_5$) and subjected to CE-ESI-MS analyses.

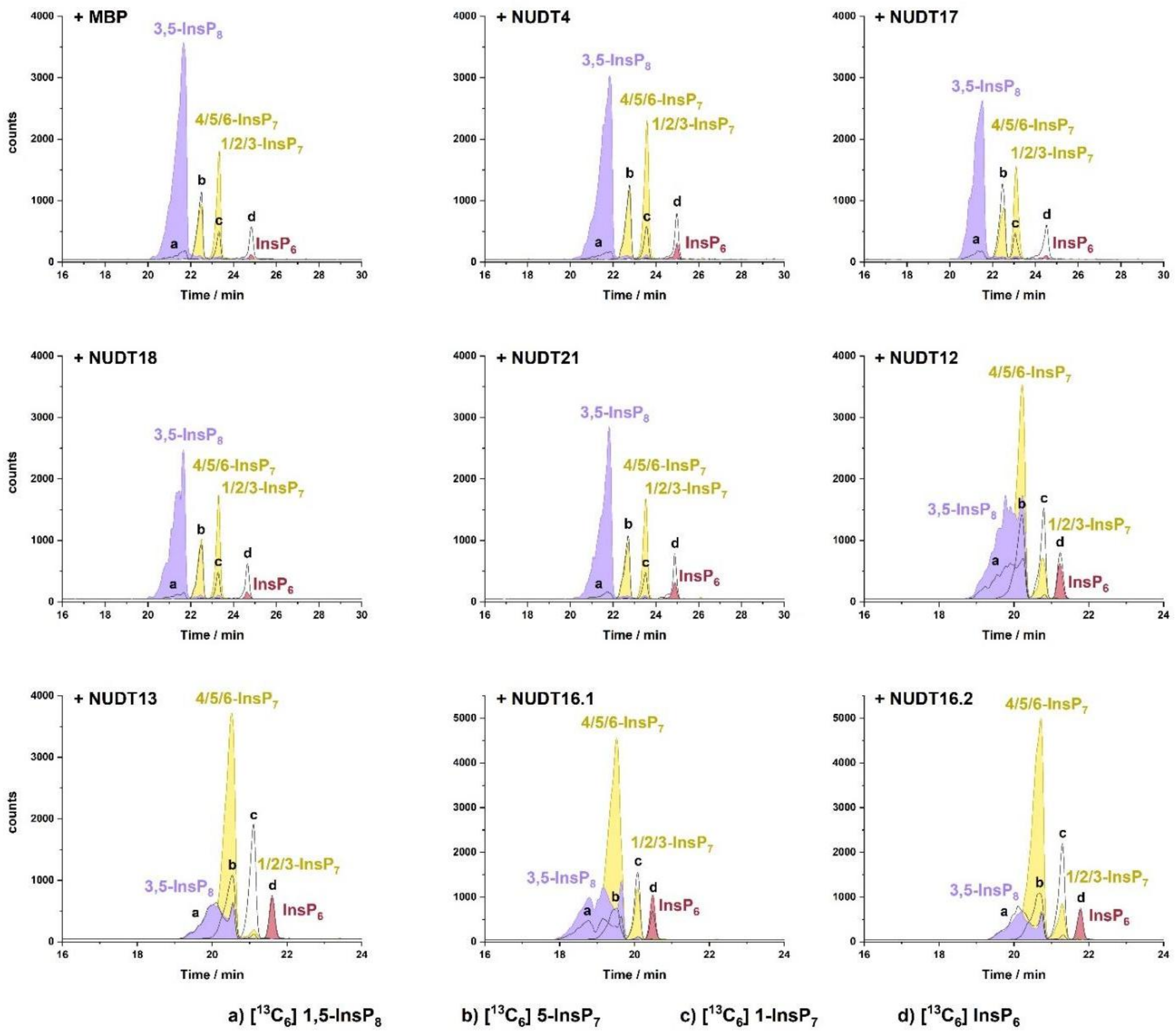

Figure S8: Arabidopsis NUDT hydrolases display differential hydrolysis activity toward 3,5-$InsP_8$ *in vitro*. Recombinant $His_6$-MBP-NUDT proteins were incubated with 0.33 mM 3,5-$InsP_8$ and 1 mM $MgCl_2$ at 28 °C. $His_8$-MBP served as a negative control. After 1 h, the reaction products were spiked with isotopic standards mixture ([$^{13}C_6$] 1,5-$InsP_8$, [$^{13}C_6$] 5-$InsP_7$, [$^{13}C_6$] 1-$InsP_7$, [$^{13}C_6$] $InsP_6$, [$^{13}C_6$] 2-OH $InsP_5$) and subjected to CE-ESI-MS analyses.

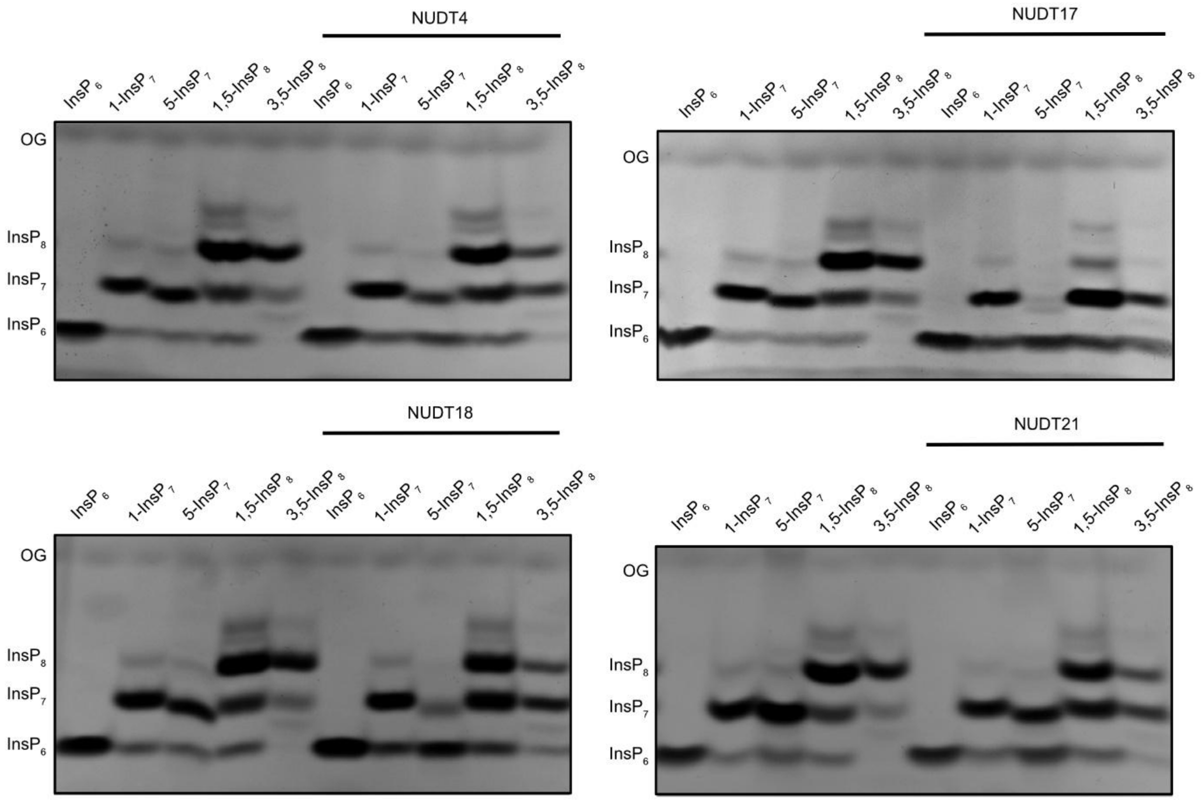

Figure S9: Subclade I NUDT hydrolases lose substrate specificity at higher concentrations *in vitro*. Recombinant $His_6$-MBP-tagged NUDT4 (~9 µM), NUDT17 (~6 µM), NUDT18 (~6 µM) or NUDT21 (~8 µM) was incubated with 0.33 mM $InsP_7$ and 1 mM $MgCl_2$ at 22 °C. $His_8$-MBP served as a negative control (placed at the left side of each gel). After 2 h, the reaction products were separated by 33 % PAGE and visualized by toluidine blue. OG: orange G.

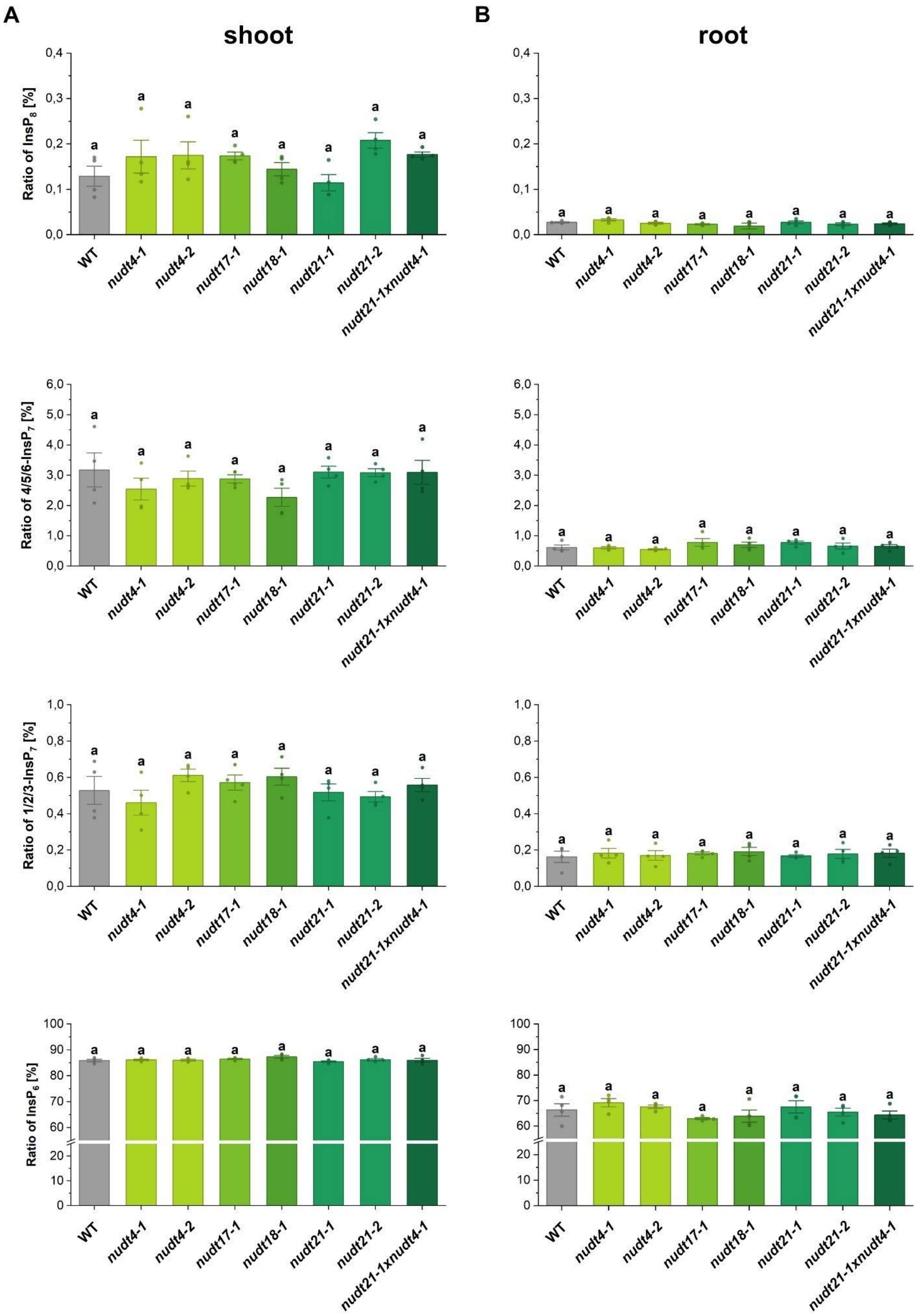

Figure S10: Single and double knockout mutants do not show a significant (PP-)InsP increase. *Arabidopsis thaliana* T-DNA insertion lines were grown hydroponically for 5 weeks. (PP-)InsPs were purified with $TiO_2$, spiked with isotopic standards mixture ($[^{13}C_6]$ 1,5-$InsP_8$, $[^{13}C_6]$ 5-$InsP_7$, $[^{13}C_6]$ 1-$InsP_7$, $[^{13}C_6]$ $InsP_6$, $[^{13}C_6]$ 2-OH $InsP_5$) and subjected to CE-ESI-MS analyses. Different letters indicate values that are significantly different determined with one-way ANOVA followed by Dunn-Šidák test with a significance level of 0.05.

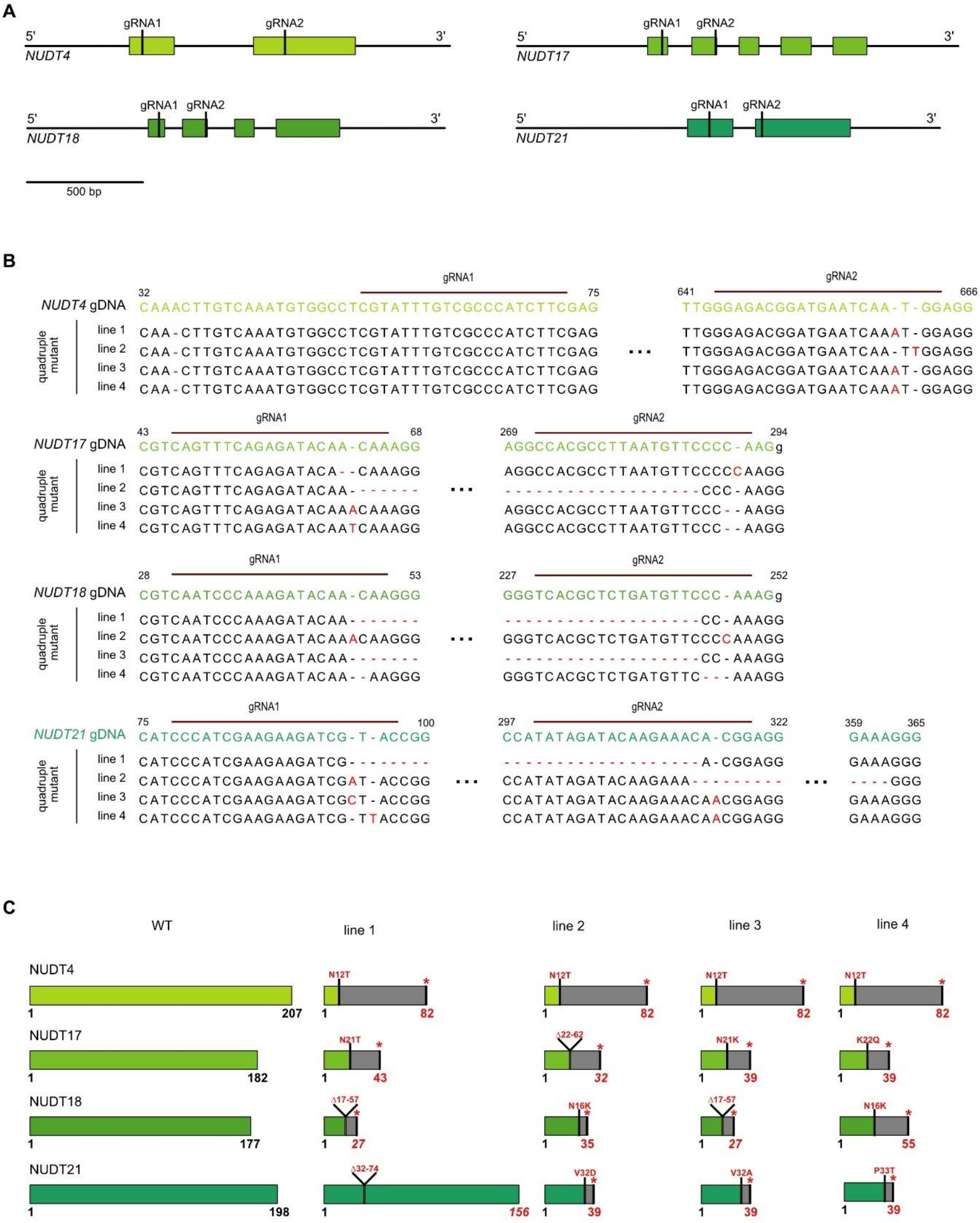

Figure S11: Schematic representation of the mutations in four subclade I *nudt4/17/18/21* mutants. (A) gRNA1 was designed to target the beginning of the open reading frame (ORF), while gRNA2 was designed to target the sequence encoding the NUDT motif (GX$_5$EX$_7$REUXEEXGU, where U represents a hydrophobic amino acid such as leucine, isoleucine or valine and X represents any amino acid), or its flanking region. (B) Observed deletions, insertions and point mutations in four *nudt4/17/18/21* mutants. (C) Predicted amino acids changes.

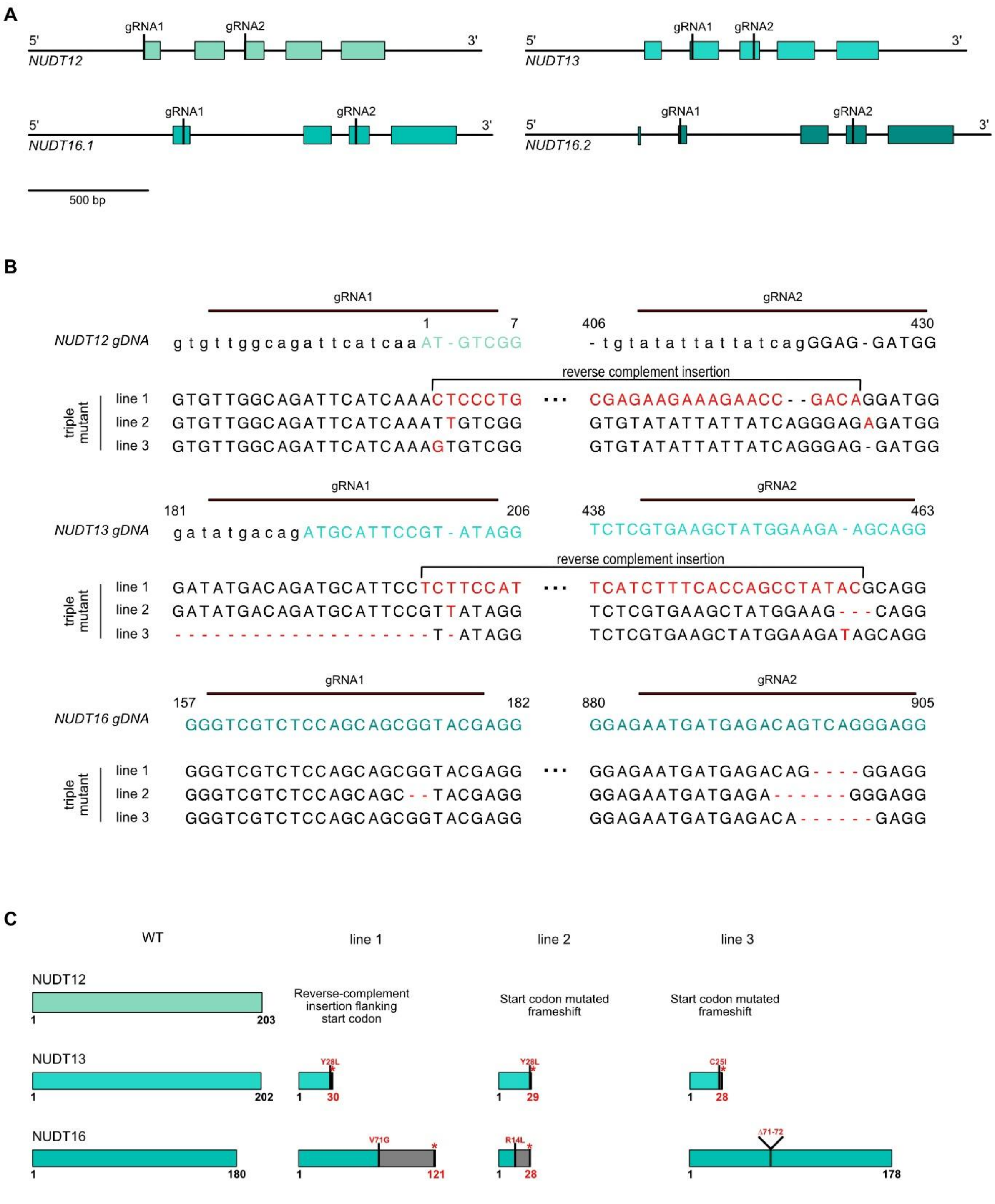

Figure S12: Schematic representation of the mutations in four subclade II *nudt12/13/16* mutants. (A) gRNA1 was designed to target the beginning of the open reading frame (ORF), while gRNA2 was designed to target the sequence encoding the NUDT motif ($GX_5EX_7REUXEEXGU$, where U represents a hydrophobic amino acid such as leucine, isoleucine or valine and X represents any amino acid), or its flanking region. (B) Observed deletions, insertions and point mutations in two *nudt12/13/16* mutants. (C) Predicted amino acids changes.

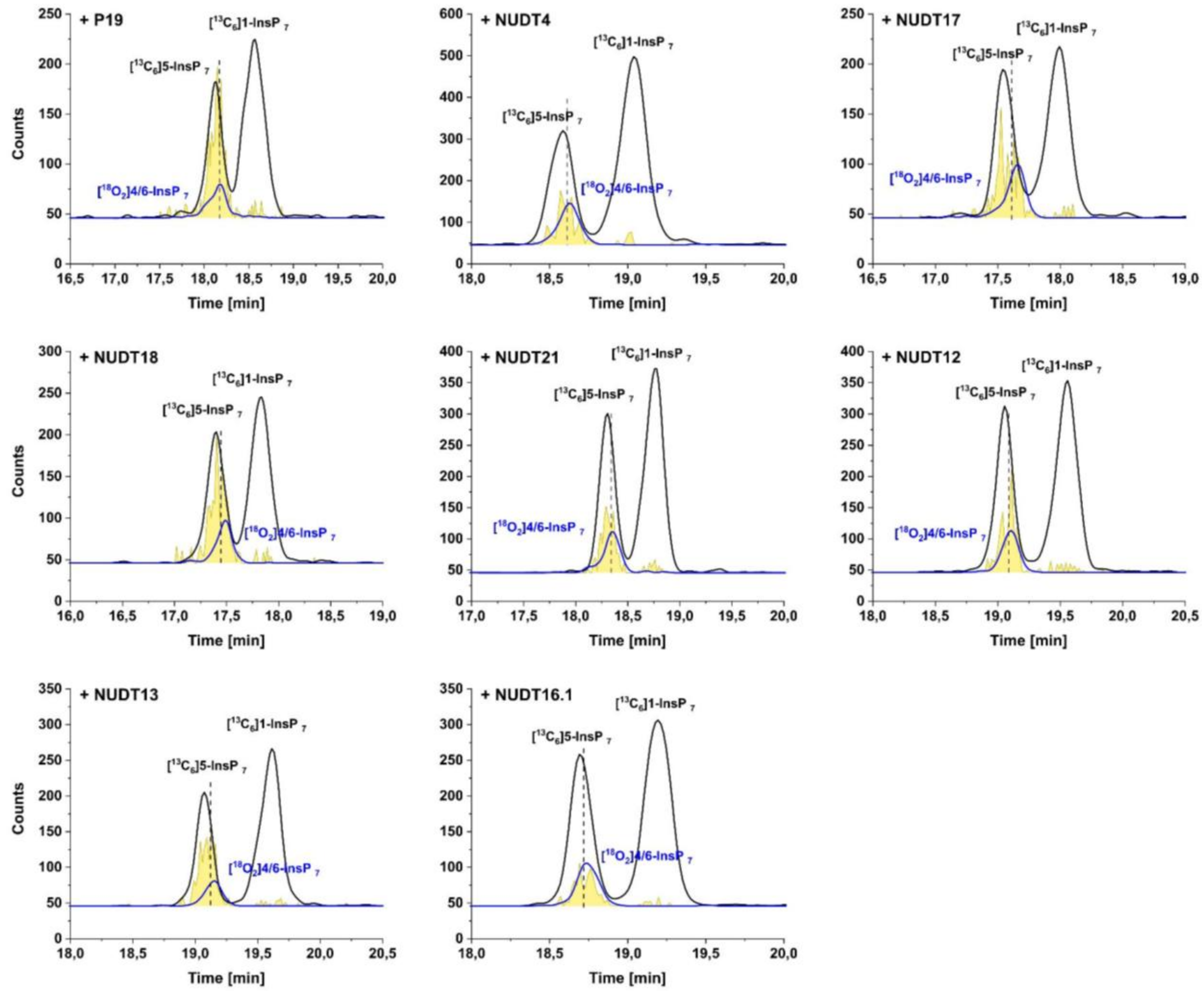

Figure S13: Transient expression of *NUDTs* in *N. benthamiana* reveals 4/6-Ins$P_7$ and 5-Ins$P_7$ turnover *in planta*.. The silencing inhibitor P19 alone or together with *NUDTs* was transiently expressed in *N. benthamiana* leaves. 2-3 days post infiltration (dpi) (PP-)InsPs were purified with $Nb_2O_5$ beads, spiked with isotopic standards mixture ([$^{13}C_6$] 1,5-Ins$P_8$, [$^{13}C_6$] 5-Ins$P_7$, [$^{18}O_2$] 4-Ins$P_7$, [$^{13}C_6$] 1-Ins$P_7$, [$^{13}C_6$] Ins$P_6$, [$^{13}C_6$] 2-OH Ins$P_5$) and subjected to CE-ESI-MS analyses.

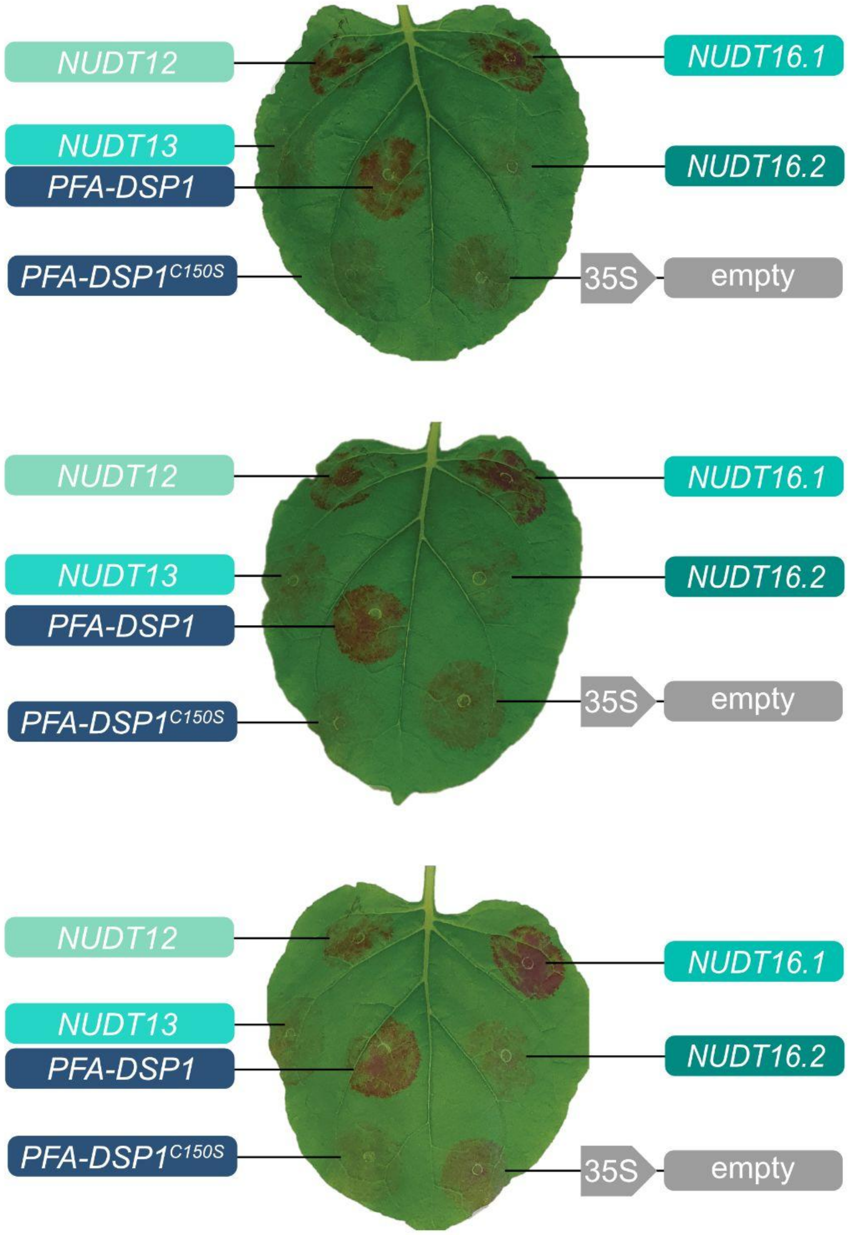

Figure S14: Transient co-expression of the *RUBY* reporter with subclade II *NUDT* hydrolase genes under the transcriptional control of the viral CaMV *35S* promoter. Co-expression with *PFA-DSP1* served as a positive control, while co-expression with the catalytic inactive *PFA-DSP1*$^{C150S}$ encoding the catalytic inactive protein or an empty vector served as negative controls. The picture was taken 3 days post infiltration.

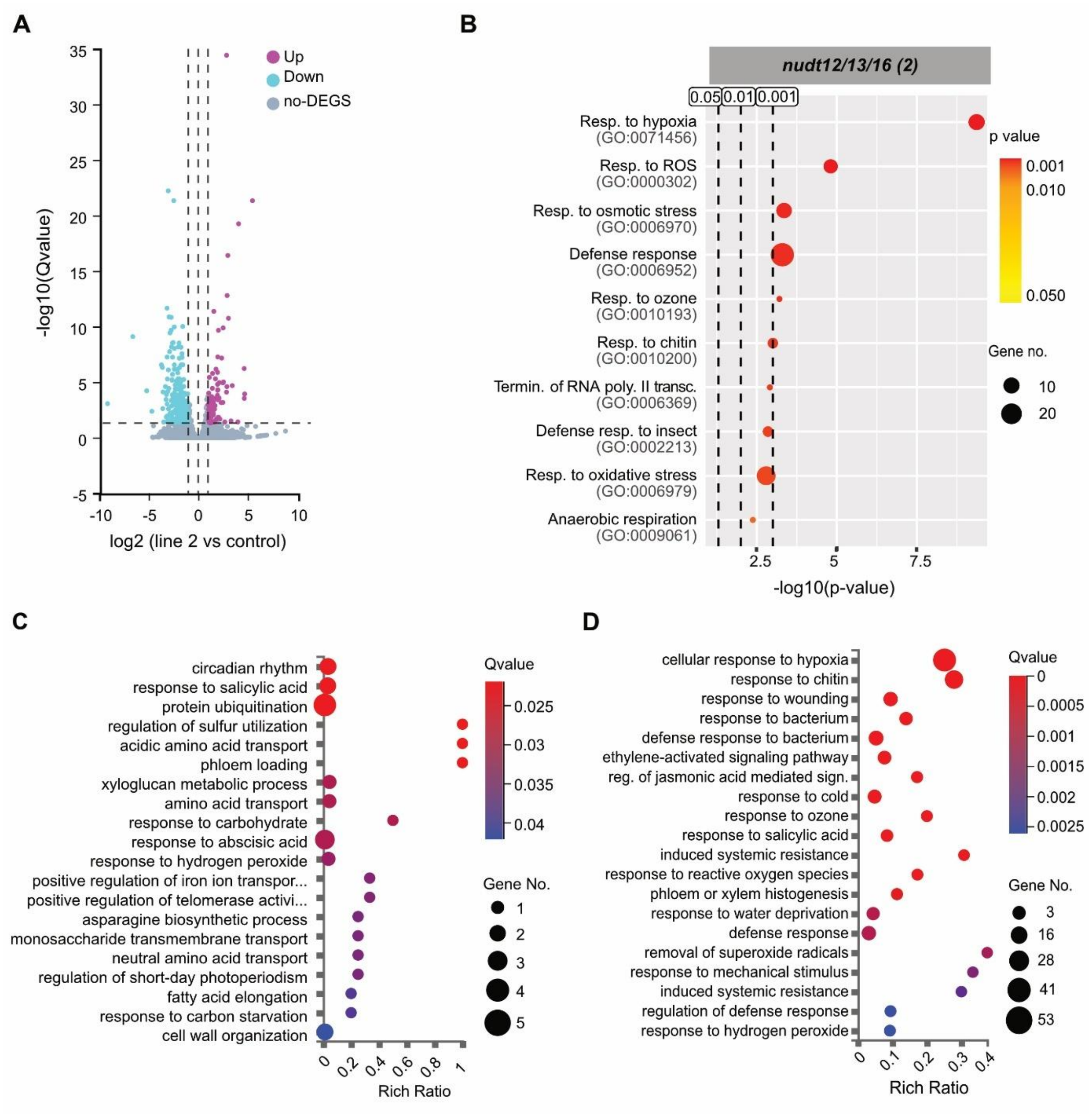

Figure S15: (A) Volcano plot comparing *nudt12/13/16* mutant line 2 to wild-type (WT). The X-axis represents the log2FC values, and the Y-axis shows -log10-transformed significance values. Magenta dots indicate upregulated DEGs, cyan dots indicate downregulated DEGs, and gray dots indicate non-DEGs. Gene Ontology (GO) enrichment analysis (B) suggests subclade II NUDT-dependent PP-InsPs regulate PSR-unrelated processes with numerous genes involved in plant defense and related GO terms. Shown is a GO enrichment analysis of DEGs with Q < 0.05 and (|log2FC| > 1), based on Biological Processes (BP) for *nudt12/13/16* triple mutant line 2. The X-axis represents the statistical significance of these GO terms in -log10 (*p*-value), while the Y-axis lists the GO terms with their respective ID. Bubble size indicates the number of DEGs annotated to each GO term, and the graph highlights three significance thresholds: 0.05, 0.01, and 0.001. The color of the bubbles corresponds to the *p*-value, with red indicating more significant enrichment. GO Enrichment bubble chart of DEGs for (C) upregulated and (D) downregulated genes with Q < 0.05 and (|log2FC| > 1), based on Biological Processes (BP) for triple mutant lines. The X-axis represents the enrichment ratio of genes, and the Y-axis represents the GO Term. The size of the bubble represents the number of differential genes annotated to a certain GO Term. The color represents the significance value of enrichment (Q-value), where red indicates smaller significance values. (Resp.: response, ROS: reactive oxygen species, syst.: systematic, termin.: termination, poly.: polymerase, transc.: transcription)

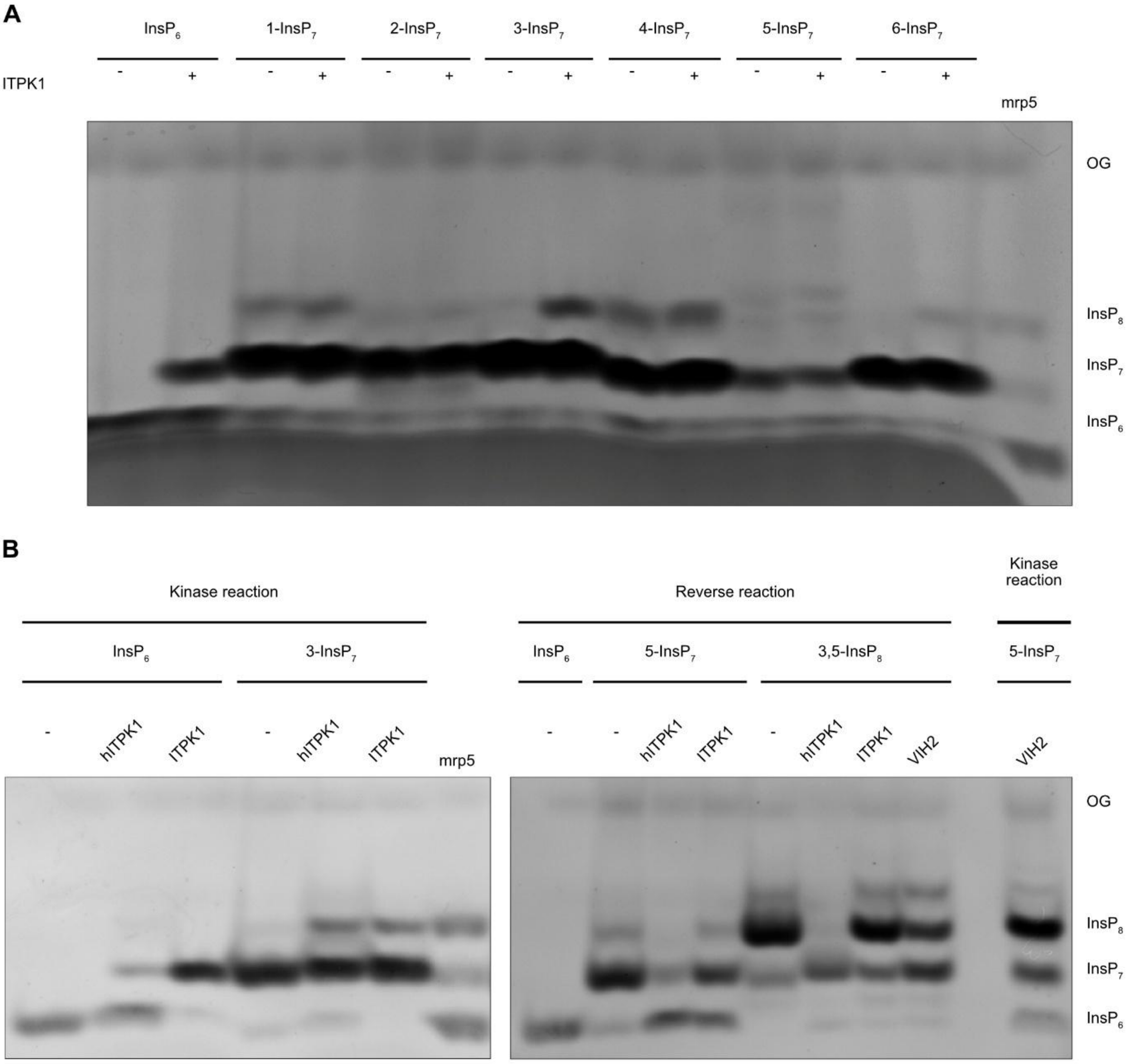

Figure S16: 3PP-InsP species are preferred substrates for evolutionarily conserved kinase and ADP phosphotransfer reactions. (A) Kinase assay with recombinant $His_8$-MBP-ITPK1 and $InsP_6$ or $InsP_7$ species as indicated. (B) Kinase assay (kinase reaction) and PP-InsP/ADP phosphotransfer assay (reverse reaction) with recombinant $His_8$-MBP-ITPK1, $His_8$-MBP-VIH2 and $His_8$-MBP-hsITPK1 and 1 mM of the indicated (PP)-InsP species. (A, B) After 6 h at 25 °C, the reaction products were separated by 33 % PAGE, and visualized by toluidine blue staining. $His_8$-MBP served as a negative control (indicated with the minus symbol). KD: kinase domain, OG: Orange G.

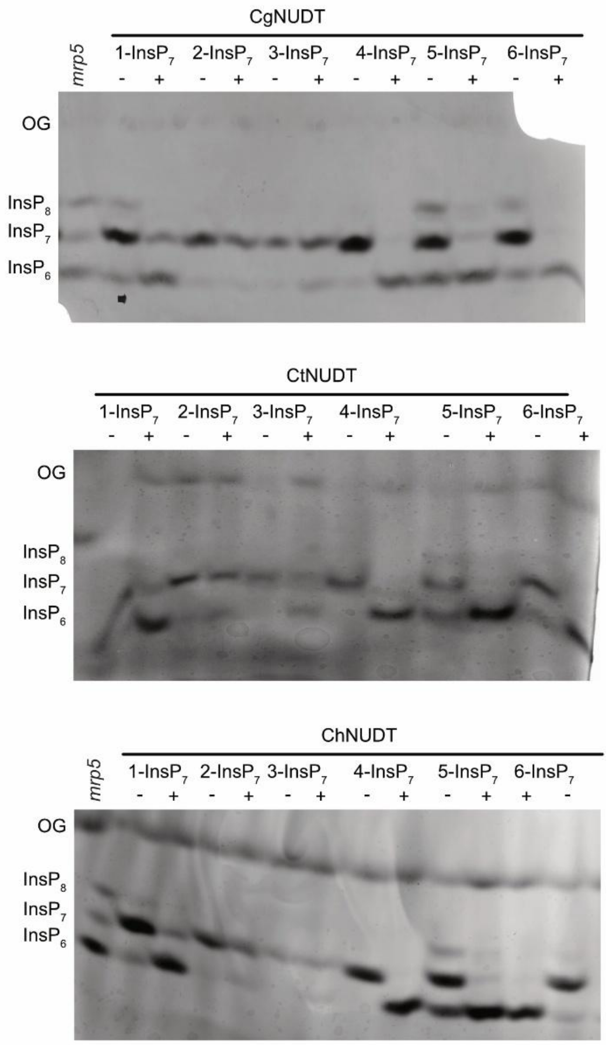

Figure S17: Fungal NUDT effectors show similar substrate specificities as subclade I NUDTs. Recombinant $His_6$-tagged CgNUDT, CtNUDT, or ChNUDT were incubated with 0.25 mM $InsP_7$. Lanes with a minus symbol show control reactions where no protein was added. After 45 min, the reaction products were separated by 33 % PAGE, and visualized by toluidine blue staining. A $TiO_2$-purified Arabidopsis *mrp5* seed extract was used as a marker for $InsP_6$, $InsP_7$ and $InsP_8$. OG: Orange G.

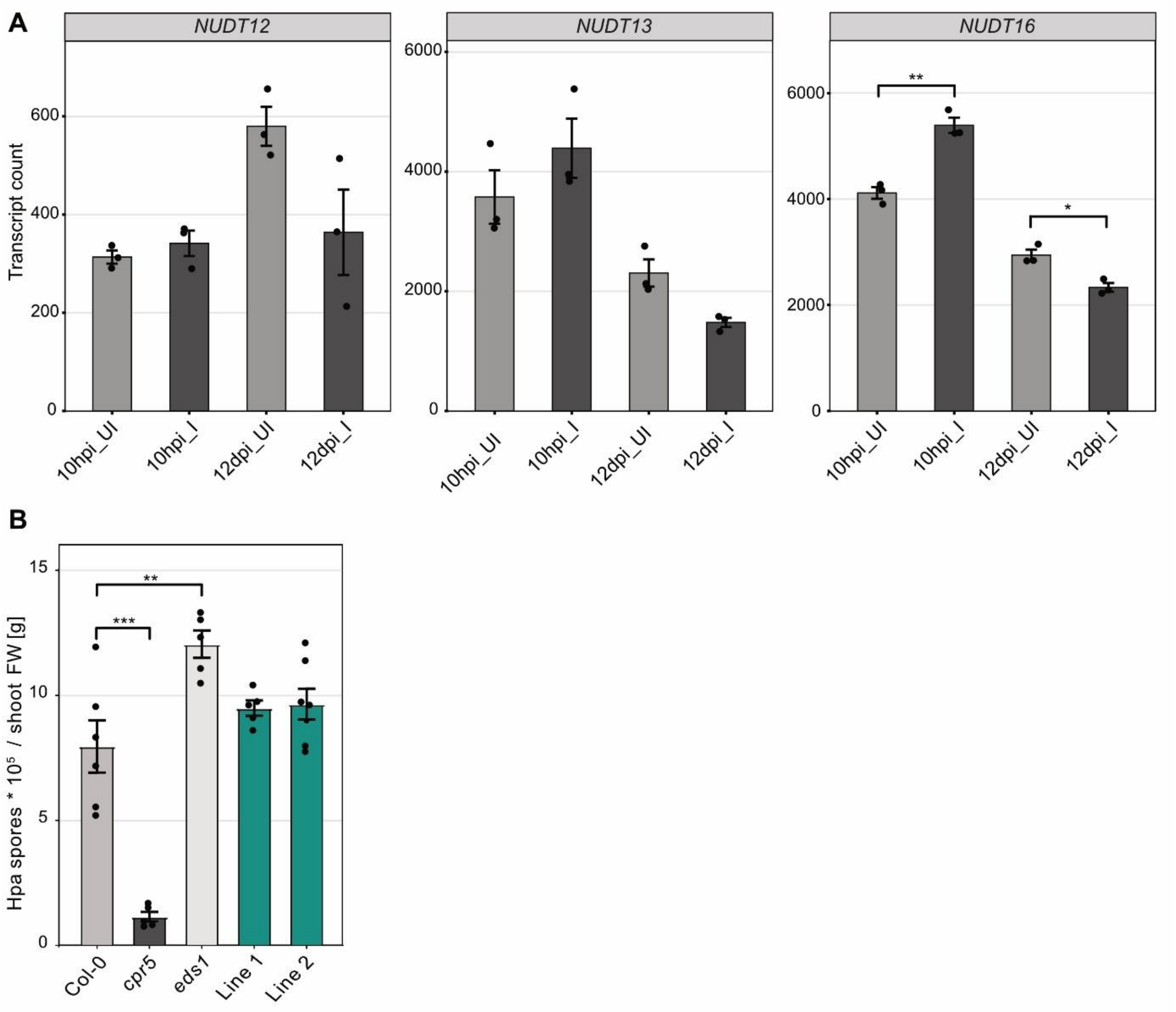

Figure S18: (A) Expression patterns of *NUDT12*, *NUDT13*, and *NUDT16* based on RNA-Seq data from Siddique et al. (2022) hpi: hours post-inoculation; dpi: days post-inoculation; UI: uninfected root tissue; I: infected root tissue. Statistical analysis was performed using the Wilcoxon test for unpaired samples ($p > 0.05$; *$p \leq 0.05$; **$p \leq 0.01$; ***$p \leq 0.001$). (B) Bar graphs represent the mean number of spores ± SEM per gram shoot fresh weight (FW) isolated from Arabidopsis wild-type (Col-0) or the indicated mutants 7 dpi with *Hyaloperonospora arabidopsidis* isolate Noco2. n = 5. Asterisks indicate significant differences to Col-0 determined by an ordinary one-way ANOVA with Dunnett's multiple comparisons test; **, $p < 0.01$; ***, $p < 0.001$.

Table S1:

List of known and putative Arabidopsis PP-InsP interactors and NUDT hydrolases identified via 5PCP-$InsP_5$ affinity pull-down. Numbers indicate identified peptides. Grouping of genes indicates that identified peptides derived from either of these two proteins.

| Gene Name | Accession Number | Shoot Beads | Shoot Eluate | Root Beads | Root Eluate |
|---|---|---|---|---|---|
| AFB1 | AT4G03190.1 | 8 | 20 | 9 | 20 |
| AFB2 | AT3G26810.1 | 5 | 3 | 1 | 2 |
| AFB3 | AT1G12820.1 | 6 | 3 | 1 | 1 |
| ASK1 | AT1G75950.1 | 5 | 9 | 4 | 10 |
| ASK2 | AT5G42190.1 | 0 | 7 | 0 | 7 |
| COI1 | AT2G39940.1 | 6 | 10 | 1 | 3 |
| CSN1 | AT3G61140.1 | 1 | 2 | 2 | 1 |
| CSN2 | AT2G26990.1 | 0 | 0 | 2 | 0 |
| CSN4 | AT5G42970.1 | 6 | 7 | 5 | 4 |
| CSN5A | AT1G22920.1<br>AT1G22920.2 | 0 | 2 | 1 | 2 |
| CUL1 | AT4G02570.1<br>AT4G02570.2<br>AT4G02570.3<br>AT4G02570.4 | 4 | 1 | 1 | 0 |
| CUL2 | AT1G43140.1 | 2 | 0 | 1 | 0 |
| IPK2α | AT5G07370.1<br>AT5G07370.2<br>AT5G07370.3<br>AT5G07370.4 | 0 | 0 | 3 | 5 |
| IPK2β | AT5G61760.1 | 0 | 2 | 4 | 3 |
| ITPK3 | AT4G08170.3 | 0 | 0 | 1 | 0 |
| MIPS3/MIPS1 | AT5G10170.1<br>AT4G39800.1 | 1 | 0 | 0 | 0 |
| NUDT16 | AT3G12600.1<br>AT3G12600.2 | 0 | 0 | 1 | 0 |
| NUDT17/NUDT18 | AT2G01670.1<br>AT1G14860.1 | 2 | 4 | 0 | 0 |
| NUDT18 | AT1G14860.1 | 0 | 0 | 0 | 1 |
| NUDT21/NUDT4 | AT1G73540.1<br>AT1G18300.1 | 5 | 7 | 0 | 0 |
| SPX1 | AT5G20150.1 | 1 | 0 | 0 | 0 |
| SPX2 | AT2G26660.1 | 0 | 0 | 2 | 0 |
| VIH1/VIH2 | AT5G15070.1<br>AT5G15070.2<br>AT3G01310.1<br>AT3G01310.2 | 4 | 4 | 13 | 7 |

Table S2: List of DEGs and no-DEGs related to P. Upregulated and downregulated genes have a |log2FC| > 1 and Q-values < 0.05. Non-significant genes have Q-values > 0.05.

| | Gene name | Gene ID | log2FC |
|---|---|---|---|
| **Upregulated** | *SDI1* | AT4G14070 | 3.07 |
| | *AAE15* | AT3G47340 | 2.80 |
| | *ASN1* | AT2G18700 | 2.17 |
| | *TPS9* | AT5G48850 | 1.67 |
| | *TPS11* | AT1G23870 | 1.64 |
| | *SDI2* | AT1G04770 | 1.11 |
| **Downregulated** | *DIC2* | AT5G04340 | -3.30 |
| | *ZAT6* | AT4G24570 | -3.06 |
| | *GLTP* | AT4G39670 | -2.98 |
| | *TIR-NBS9* | AT1G72920 | -2.37 |
| | | AT4G36010 | -2.03 |
| | *ACS6* | AT4G11280 | -1.99 |
| | *CaLB1* | AT4G34150 | -1.85 |
| | *DIC1* | AT2G22500 | -1.84 |
| | PR5 | AT1G75040 | -1.68 |
| | *MYB62* | AT1G68320 | -1.62 |
| | *PAP1* | AT2G01180 | -1.48 |
| | *ATL80* | AT1G20823 | -1.4 |
| | *CaLB domain* | AT3G16510 | -1.38 |
| | *SOT17* | AT1G18590 | -1.28 |
| | *GPAT6* | AT2G38110 | -1.21 |
| **Non significant** | *SPX1* | AT5G20150 | |
| | *BAH1* | AT1G02860 | |
| | *WRKY6* | AT1G62300 | |

| | |
|---|---|
| *VPT3* | AT4G22990 |
| *PHB* | AT2G34710 |
| *PHT4;5* | AT5G20380 |
| *LPR1* | AT1G23010 |
| *BAK1* | AT4G33430 |
| *PHR2* | AT2G47590 |
| *PAP10* | AT2G16430 |
| *SPX3* | AT2G45130 |
| *STOP1* | AT1G34370 |
| *SIZ1* | AT5G60410 |
| *PHL4* | AT2G20400 |
| *SPX2* | AT2G26660 |
| *PHR1* | AT4G28610 |
| *PEPR2* | AT1G17750 |
| *PBL12* | AT2G26290 |
| *PHO2* | AT2G33770 |
| *VPT1* | AT1G63010 |
| *MYB62* | AT1G68320 |
| *CERK1* | AT3G21630 |
| *PHT1;4* | AT2G38940 |
| *GLP1* | AT1G72610 |
| *bHLH050* | AT1G73830 |
| *VIH2* | AT3G01310 |
| *EAL1* | AT4G37650 |
| *PHT3;1* | AT5G14040 |
| *ALS3* | AT2G37330 |
| *MRP5* | AT1G04120 |
| *ALMT1* | AT1G08430 |

| | |
|---|---|
| *CLV2* | AT1G65380 |
| *LPR2* | AT1G71040 |
| *PAP12* | AT2G27190 |
| *PHT5* | AT2G32830 |
| *MOR1* | AT2G35630 |
| *SPDT* | AT3G15990 |
| *RALF23* | AT3G16570 |
| *PHO1* | AT3G23430 |
| *PHL2* | AT3G24120 |
| *PHT2;1* | AT3G26570 |
| *PHL3* | AT4G13640 |
| *ITPK2* | AT4G33770 |
| *IPS2* | AT5G03545 |
| *WRKY75* | AT5G13080 |
| *VIH1* | AT5G15070 |
| *SPX4* | AT5G15330 |
| *ITPK1* | AT5G16760 |
| *SEC12* | AT2G01470 |
| *PDR2* | AT5G23630 |

Table S3: Most highly downregulated and upregulated genes, defined as |log2FC| > 3.

| | Gene name | Gene ID | log2FC |
|---|---|---|---|
| **Downregulated** | *PAO2* | AT2G43020 | -9,57 |
| | U-box E3 ubiquitin | AT3G02840 | -4,09 |
| | *CMF4* | AT1G63820 | -3,88 |
| | *ATS40-2* | AT5G45630 | -3,87 |
| | *RAS1* | AT1G09950 | -3,83 |
| | *WRKY40* | AT1G80840 | -3,64 |
| | *DVL10* | AT4G13395 | -3,64 |
| | *ERF022* | AT1G33760 | -3,63 |
| | *ERF11* | AT1G28370 | -3,53 |
| | | AT4G29780 | -3,52 |
| | *IDL7* | AT3G10930 | -3,49 |
| | *CCR4* | AT5G47850 | -3,44 |
| | *DIC2* | AT4G24570 | -3,29 |
| | *DHYPRP1* | AT4G22470 | -3,27 |
| | *DTX50* | AT5G52050 | -3,10 |
| | | AT5G16200 | -3,07 |
| | *ZAT6* | AT5G04340 | -3,06 |
| | *ZAT10* | AT1G27730 | -3,01 |
| | *ERF6* | AT4G17490 | -3,01 |
| **Upregulated** | *SDI1* | AT5G48850 | 3,06 |
| | *IRONMAN 3* | AT2G30766 | 3,71 |
| | | AT2G05540 | 3,78 |
| | *LTP3* | AT5G59320 | 4,45 |
| | *IRP6* | AT5G05250 | 4,46 |
| | | AT5G35935 | 5,64 |

Table S4: List of DEGs and no-DEGs related to Fe. Upregulated and downregulated genes have a |log2FC| > 1 and Q-values < 0.05. Non-significant genes have Q-values > 0.05.

| | Gene name | Gene ID | log2FC |
|---|---|---|---|
| **Upregulated** | *IRP6* | AT5G05250 | 4.46 |
| | *IRONMAN 3* | AT2G30766 | 3.72 |
| **Downregulated** | *ZAT12* | AT5G35735 | -2.66 |
| | *FER1* | AT5G01600 | -2.02 |
| | *HYP1* | AT5G59820 | -1.30 |
| **Non- significant** | *NRAMP2* | AT1G47240 | |
| | *NAS1* | AT5G04950 | |
| | *NRAMP1* | AT1G80830 | |
| | *DEG18* | AT4G12980 | |
| | *CRR* | AT3G25290 | |
| | *YSL6* | AT3G27020 | |
| | *BHLH115* | AT1G51070 | |
| | *AHA2* | AT4G30190 | |
| | *BTS* | AT3G18290 | |
| | *AHA8* | AT3G42640 | |
| | *BHLH38* | AT3G56970 | |
| | *YSL2* | AT5G24380 | |
| | *AHA10* | AT1G17260 | |
| | *LPR1* | AT1G23010 | |
| | *IRT3* | AT1G60960 | |
| | *AIR12* | AT3G07390 | |
| | *COSY* | AT1G28680 | |
| | *VTL1* | AT1G21140 | |

| | |
|---|---|
| *MYB10* | AT3G12820 |
| *BHLH34* | AT3G23210 |
| *CIPK11* | AT2G30360 |
| *AHA1* | AT2G18960 |
| *VTL2* | AT1G76800 |
| *YSL5* | AT3G17650 |
| CYBDOMs | AT5G54830 |
| *YSL3* | AT5G53550 |
| *NAS4* | AT1G56430 |
| *AHA3* | AT5G57350 |
| *IRONMAN 2* | AT1G47395 |
| *BHLH121* | AT3G19860 |
| *BHLH29* | AT2G28160 |
| *BHLH105* | AT5G54680 |
| *PYE* | AT3G47640 |
| *AHA4* | AT3G47950 |
| *BHLH100* | AT2G41240 |
| CYBDOMs | AT5G47530 |
| *YSL4* | AT5G41000 |
| *YSL1* | AT4G24120 |
| *AHA5* | AT2G24520 |
| *AHA9* | AT1G80660 |
| *AHA11* | AT5G62670 |
| *YSL8* | AT1G48370 |
| *YSL7* | AT1G65730 |
| *IRONMAN 1* | AT1G47400 |
| *NRAMP6* | AT1G15960 |
| *BHLH39* | AT3G56980 |

| | |
|---|---|
| *NAS3* | AT1G09240 |
| *BTSL2* | AT1G18910 |
| *LPR2* | AT1G71040 |
| *BTSL1* | AT1G74770 |
| *NRAMP3* | AT2G23150 |
| CYBDOMs | AT3G07570 |
| *FRD3* | AT3G08040 |
| *F6'H1* | AT3G13610 |
| *VTL5* | AT3G25190 |
| *PDR9* | AT3G53480 |
| *MTP8* | AT3G58060 |
| *CYBDOMs* | AT3G59070 |
| *AHA7* | AT3G60330 |
| *BHLH104* | AT4G14410 |
| CYBDOMs | AT4G17280 |
| *NRAMP5* | AT4G18790 |
| *IRT1* | AT4G19690 |
| *BHLH101* | AT5G04150 |
| *ZIF1* | AT5G13740 |
| *NRAMP4* | AT5G67330 |

Table S5: List of primers used in this study.

**Cloning primer**

| **Target gene** | **Objective** | **Primer sequence 5'-3'** |
|---|---|---|
| *NUDT4* | attb1 cDNA | AAAAAGCAGGCTTC ATGACAGGGTTCTCTGTGTC |
| | attb1 promoter | AAAAAGCAGGCTTC GTCCGACTTTAAAGAGAAATTGAGG |
| | attb2 stop | AGAAAGCTGGGTC TCAGTTCCCACTTTCATCATCG |
| | attb2 no stop | AGAAAGCTGGGTC GTTCCCACTTTCATCATCGTC |
| | attb2 no stop V5 | ACCTCCTCCAGATCCGTTCCCACTTTCAT CATCGTC |
| | genotyping | F:AAAAAGCAGGCTTC GTCCGACTTTAAAGAGAAATTGAGG |
| | | R: AGAAAGCTGGGTC GTTCCCACTTTCATCATCGTC |
| | | Seq:ACTCTTTCTCTCTTGTTCTG |
| *NUDT17* | attb1 cDNA | AAAAAGCAGGCTTC ATGGGTGTTGAGAAAATGGTG |
| | attb1 promoter | AAAAAGCAGGCTTC GTCGTATTGTATGTTCCATGC |
| | attb2 stop | AGAAAGCTGGGTC TCAACACATTGTTTCAATAGAGATTG |
| | attb2 no stop | AGAAAGCTGGGTC ACACATTGTTTCAATAGAGATTGAC |
| | attb2 no stop V5 | ACCTCCTCCAGATCCACACATTGTTTCAA TAGAGATTGAC |
| | genotyping | F: AAAAAGCAGGCTTC GTCGTATTGTATGTTCCATGC |
| | | R:AGAAAGCTGGGTC TCAACACATTGTTTCAATAGAGATTG |
| | | Seq:CATCAGACCCTTCTCTTCTC |
| *NUDT18* | attb1 cDNA | AAAAAGCAGGCTTC ATGGTGTGTTTGGTCTCCC |
| | attb1 promoter | AAAAAGCAGGCTTC GAGCTTCTTCTAGATCGAGCTG |
| | attb2 stop | AGAAAGCTGGGTC TCAGTAGATAGAGATCAGTGGAAG |

| | | |
|---|---|---|
| | attb2 no stop | AGAAAGCTGGGTC GTAGATAGAGATCAGTGGAAGGTTC |
| | attb2 no stop V5 | ACCTCCTCCAGATCCGTAGATAGAGATC AGTGGAAGGTTC |
| | genotyping | F:GGATAAGCCGCAAGGAGG |
| | | R:TTTCCAAGTTGTCTCTGCACC |
| | | Seq:CTCACCCCATCATAACTTC |
| *NUDT21* | attb1 cDNA | AAAAAGCAGGCTTCATGATTTCTCTATTC ATCTCAAACTTTTC |
| | attb1 promoter | AAAAAGCAGGCTTC CTCCGTAAATACCGTGTTGG |
| | attb2 stop | AGAAAGCTGGGTC TTATTGGGTCTGGCATTTCC |
| | attb2 no stop | AGAAAGCTGGGTC TTGGGTCTGGCATTTCC |
| | attb2 no stop V5 | ACCTCCTCCAGATCCTTGGGTCTGGCAT TTCC |
| | genotyping | F: AAAAAGCAGGCTTC CTCCGTAAATACCGTGTTGG |
| | | R: AGAAAGCTGGGTC TTATTGGGTCTGGCATTTCC |
| | | Seq:ATAAAGACGCTCGCAAAC |
| *NUDT12* | attb1 cDNA | AAAAAGCAGGCTTC ATGTCGGTTCTTTCTTCTCG |
| | attb1 promoter | AAAAAGCAGGCTTC CCCTTCTGTTTCGTACATGC |
| | attb2 stop | AGAAAGCTGGGTC CTAGTTAACTACAAAACAGTACCAAGG |
| | attb2 no stop | AGAAAGCTGGGTCGTTAACTACAAAACA GTACCAAGG |
| | attb2 no stop V5 | ACCTCCTCCAGATCCGTTAACTACAAAA CAGTACCAAGG |
| | qPCR | F: AACTCGAGGATTGGCCAGAGCGA |
| | | R: CCGACAAAGCTCCAACGCTTCT |
| | genotyping | F: CCCTTCTGTTTCGTACATGC |
| | | R: CTAGTTAACTACAAAACAGTACCAAGG |
| | | Seq: CCCTGACTGTCTCATCATTC |
| *NUDT13* | attb1 cDNA | AAAAAGCAGGCTTC ATGTCGAATCTTTCTGCAAG |
| | attb1 promoter | AAAAAGCAGGCTTC GGGGACATTTGTTCTACACAG |

| | | |
|---|---|---|
| | attb2 stop | AGAAAGCTGGGTC TTAGACTACAAAGCAGTAGCG |
| | attb2 no stop | AGAAAGCTGGGTCGACTACAAAGCAGTAGCGAG |
| | attb2 no stop V5 | ACCTCCTCCAGATCCGACTACAAAGCAGTAGCGAG |
| | qPCR | F: AGGCTGGTGAAAGATGAAGAAGA |
| | | R: TCCCATCCTCCCTTTGGGAA |
| | genotyping | F: ATGTCGAATCTTTCTGCAAG |
| | | R: TTAGACTACAAAGCAGTAGCG |
| | | Seq: TTAGACTACAAAGCAGTAGCG |
| *NUDT16 both* | attb1 promoter | AAAAAGCAGGCTTC CAATTACCTGCGATCTCTCTCTG |
| | attb2 stop | AGAAAGCTGGGTC TCAATGTTCACCAGTTATCTCCTCTCC |
| | attb2 NS | AGAAAGCTGGGTC ATGTTCACCAGTTATCTCCTCTCC |
| | attb2 NS V5 | ACCTCCTCCAGATCCATGTTCACCAGTTATCTCCTCTCC |
| | qPCR | F: CGCCGGGTGTATTCCGTTTA |
| | | R: AGGTCCACTAGACGAGCTGA |
| | genotyping | F: CAATTACCTGCGATCTCTCTCTG |
| | | R: ATGTTCACCAGTTATCTCCTCTCC |
| | | Seq: AGCTATAGACAGACACACCC |
| *NUDT16.1* | attb1 cDNA | AAAAAGCAGGCTTCATGTGTGATTTGGTCGCGCG |
| | qPCR | F: TGTCTCCAAGCTGGGTGTTT |
| | | R: CGCGACCAAATCACACATGA |
| *NUDT16.2* | attb1 cDNA | AAAAAGCAGGCTTCATGGTAGAGCAGCGGTACGAG |
| *MoNUDT*$^{E79Q}$ | mutagenesis forward | CTGGGAAACCTGTGTGCGCCGTCAAGCGCGAGAAGAAGGCGGATTTACTTTG |
| *MoNUDT*$^{E79Q}$ | mutagenesis reverse | CAAAGTAAATCCGCCTTCTTCTCGCGCTTGACGGCGCACACAGGTTTCCCAG |
| General | attb1 adapter | GGGGACAAGTTTGTACAAAAAAGCAGGCTTC |
| | attb1 adapter | GGGGACCACTTTGTACAAGAAAGCTGGGTC |

| | | |
|---|---|---|
| | attb2 V5 adapter | GGGGACCACTTTGTACAAGAAAGCTGGG TCTTAC |
| | | GTAGAATCGAGACCGAGGAGAGGGTTA GGGATA |
| | | GGCTTACCTCCTCCAGATCC |

| **Line/Plasmid** | **Primer Sequence** |
|---|---|
| SALK_102051.27.90.x | F: CAACAACGTTTGGGCTTCTAG |
| | R: ATTGTGAATTTGAACAACGGC |
| SAIL_1211_A06 | F: TTGGTGGTTTACGAGTTGACC |
| | R: TCAACACGTTATTCCAAATTGC |
| SAIL_500_D10 | F: GAAAAATTGCTCCACACTTGC |
| | R: TGAAGAAGAACGTACCCAACG |
| SALKseq_117563.1 | F: AAACCAAACCGAAATCCAAAC |
| | R: AGATCCTCCTTCCCTTTAGCC |
| SALK_031788.56.00.x | F: TCACCCACTTGCTTGCTACTC |
| | R: TGCTTTTGTATTGCCATTTCC |
| SALKseq_132932.101 | F: AAATACCGTGTTGGCATGAAC |
| | R: TGCTTTTGTATTGCCATTTCC |
| oRU906 | GAGTCTATGATCAAGTAATTATGC |
| oRU908 | GCTTGCATGCCTGCAGGTCGACTCT |
| oRU385 | CAACGCGTTGGGAGCTCTCCCATATG |
| LBa1 SALK | TGGTTCACGTAGTGGGCC |
| LB2 SAIL | GCTTCCTATTATATCTTCCCAAATTACCAATACA |

| **qPCR** | **Primer Sequence 5'-3'** |
|---|---|
| *ACTIN2* | F: GACCAGCTCTTCCATCGAGAA |
| | R: CAAACGAGGGCTGGAACAAG |
| *UBQ2* | F: AGACGAACGCAAAGATGCAG |
| | R: CCGGCGAAGATCAACCTCTG |

Table S6: List of primers used to generate sgRNAs for the *NUDT* knockout mutants. Overhangs for oligo annealing are highlighted.

| Target gene | gRNA | Forward primer | Reverse primer |
|---|---|---|---|
| *NUDT17* | sgRNA1 | **ATTG** CAGTTTCAGAGATACAACAA | **AAAC** TTGTTGTATCTCTGAAACTG |
| *NUDT17* | sgRNA2 | **GTCA** CCACGCCTTAATGTTCCCCA | **AAAC** TGGGGAACATTAAGGCGTGG |
| *NUDT18* | sgRNA1 | **ATTG** CAATCCCAAAGATACAACAA | **AAAC** TTGTTGTATCTTTGGGATTG |
| *NUDT18* | sgRNA2 | **GTCA** TCACGCTCTGATGTTCCCAA | **AAAC** TTGGGAACATCAGAGCGTGA |
| *NUDT21* | sgRNA1 | **ATTG** CCCATCGAAGAAGATCGTAC | **AAAC** GTACGATCTTCTTCGATGGG |
| *NUDT21* | sgRNA2 | **GTCA** TATAGATACAAGAAACACGG | **AAAC** CCGTGTTTCTTGTATCTATA |
| *NUDT4* | sgRNA1 | **ATTG** GAAGATGGGCGACAAATACG | **AAAC** CGTATTTGTCGCCCATCTTC |
| *NUDT4* | sgRNA2 | **GTCA** GGAGACGGATGAATCAATGG | **AAAC** CCATTGATTCATCCGTCTCC |
| *NUDT12* | sgRNA1 | **ATTG** TTGGCAGATTCATCAAATGT | **AAAC** ACATTTGATGAATCTGCCAA |
| *NUDT12* | sgRNA2 | **GTCA** ATATTATTATCAGGGAGGAT | **AAAC** ATCCTCCCTGATAATAATAT |
| *NUDT13* | sgRNA1 | **ATTG** ATGACAGATGCATTCCGTAT | **AAAC** ATACGGAATGCATCTGTCAT |
| *NUDT13* | sgRNA2 | **GTCA** CGTGAAGCTATGGAAGAAGC | **AAAC** GCTTCTTCCATAGCTTCACG |
| *NUDT16* | sgRNA1 | **ATTG** TCGTCTCCAGCAGCGGTACG | **AAAC** CGTACCGCTGCTGGAGACGA |
| *NUDT16* | sgRNA2 | **GTCA** GAATGATGAGACAGTCAGGG | **AAAC** CCCTGACTGTCTCATCATTC |
